# The bacterial antiphage protein Tiamat is an ATPase and DNase whose homologs are found across the tree of life

**DOI:** 10.64898/2026.09.30.754977

**Authors:** Shirley Yuan, Yuri Rafael de Oliveira Silva, Balwina Koopal, Vinh Dang, Arav A. Nangia, Jack P.K. Bravo, Andrew Santiago-Frangos

**Affiliations:** Department of Biology, University of Pennsylvania, Philadelphia PA 19104, USA; Institute of Science and Technology Austria, Klosterneuburg 3400, Austria

**Author notes:** Current address: Department of Microbiology, Institute of Biomedical Sciences, University of São Paulo, São Paulo, Brazil (Yuri Rafael de Oliveira Silva). Current address: Sidney Kimmel Medical College, Thomas Jefferson University, Philadelphia, PA 19107, USA. Shirley Yuan and Yuri Rafael de Oliveira Silva contributed equally to this work.

## Abstract

Bacteria and archaea encode hundreds of antiviral defense systems, some with homologs found in eukaryotic innate immunity. GHL ATPases like plant MORCs and prokaryotic Tiamat defend against viruses and phage, though their architecture and biochemical activities have remained unclear. Searching thousands of proteomes, we find Tiamat is rare in prokaryotes yet enriched across eukaryotes, including green plants, aquatic chordates, sponges, placozoans, and water molds. We resolve the central DUF3684 and flanking region into three predicted folds (TPR-like repeats, an uncharacterized conserved fold, and an inactive metalloprotease-like fold) that occupy the substrate-binding position of other GHL proteins. We show that purified Bacillus cereus Tiamat hydrolyzes ATP, inefficiently nicks duplex DNA, and cleaves single-stranded DNA site-specifically. We identify Tiamat-like proteins whose C-terminal nuclease is often swapped for helicases, peptidases, or other effectors. Additionally, we show Tiamat genes neighbor helicases, DNA methyltransferases, and nucleases, suggesting some Tiamats degrade foreign nucleic acid within a multi-component complex. Eukaryotic Tiamat homologs like the close Arabidopsis homolog NO VEIN and the distant human homolog SACSIN have acquired critical non-defense roles, leaving open whether these homologs remain antiviral or have been exapted.

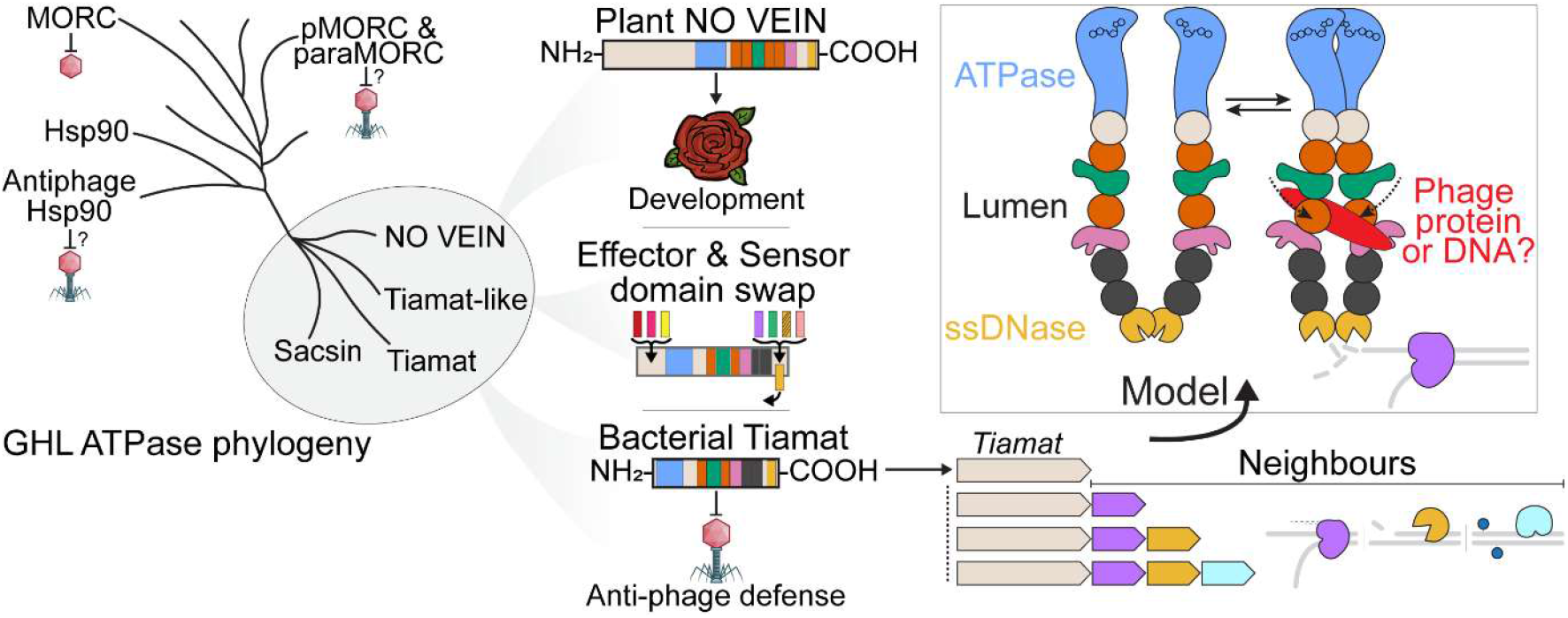
GRAPHICAL ABSTRACT.

## INTRODUCTION

Viral infection and parasitism have driven the evolution and diversification of ancient antiviral defense systems across the three domains of life, a pressure that likely dates back to the life’s origin (1). Beyond restriction-modification (R-M) and CRISPR-Cas systems, bacteria and archaea encode hundreds of antiviral defense systems (2–5). Consistent with this ancient origin, homologs of diverse antiviral defense systems are found across bacteria and eukaryotes (6, 7). Some of these widespread protein families have retained their antiviral function. For example, the interferon-inducible protein Viperin converts cytidine triphosphate to a chain terminator, inhibiting the replication of a broad range of mammalian viruses (8, 9), and bacterial Viperin homologs were similarly shown to defend against phage infection (10, 11). In fact, the antiviral function of Viperin is so strongly conserved that human Viperin heterologously expressed in *Escherichia coli* protects the bacterium from phage infection (11).

In contrast, other ancient defense proteins have diversified for a range of functions that include non-defense roles. For example, eukaryotic Argonaute proteins, originally discovered to regulate leaf development in *Arabidopsis thaliana* (12), silence host and viral mRNAs in an RNA-guided manner, and their activities are often diversified by long N-terminal extensions (13). Similarly, prokaryotic Argonaute homologs can have roles in antiviral defense via RNA-guided disruption of membrane integrity or depletion of key cellular metabolites (14–16), or non-antiviral roles that include DNA-guided disentanglement of circular chromosome replication (17). Modular swapping of effector domains onto a conserved Argonaute core and complex formation with proteins encoded nearby facilitate these diverse Argonaute activities (14–17). We hypothesize that other antiviral proteins have homologs with non-defense roles critical to cell or organismal health.

Recent studies have uncovered antiviral members of the GHKL ATPase superfamily, whose conventional members like Gyrase, Hsp90, Histidine Kinases, and MutL are best known for their non-defense roles in core cellular processes (3, 18–21). Hsp90, type II topoisomerases and MutL constitute a distinct subfamily, the GHL proteins, whose members assemble into homodimers or A_2_B_2_ heterotetramers (18). GHL proteins share an architecture that consists of a constitutive C-terminal dimerization domain, one or more middle domains that bind a substrate, and an N-terminal ATPase domain that dimerizes upon ATP binding, closing a clamp around substrate held in the lumen of the dimer (22–31). Within the middle region, an S5 domain 2-like fold (S5-fold) engages chemically distinct substrates in different members. Perhaps akin to defense system modularity, modular swapping of the domains C-terminal to a GHL ATPase domain enables this superfamily of proteins to engage diverse DNA and protein substrates and carry out critical but distinct biological functions.

For example, Hsp90 engages client polypeptides and co-chaperones (25, 32), whereas the topoisomerase IV subunit ParE binds a DNA duplex that must pass through a transient double-strand break (29, 33). The most well-studied example of GHL proteins that can have antiviral roles are the MORC (Microrchidia) family, which are nuclear proteins involved in transcriptional repression, RNA-mediated DNA methylation, and DNA repair (34–37). Recent studies have revealed that MORC homologs contribute to antiviral defense in plants, and are found in both bacteria and eukaryotes (21, 34–37). Antiphage activity, however, remains rare among the GHL protein family (2, 3, 5, 18, 20, 21, 38–43).

Tiamat is one of the few GHL proteins with a demonstrated antiphage role (3). It was identified in a large-scale computational analysis that found its gene enriched within “defense islands”, or genomic regions that contain many antiphage defense genes (3). While many defense systems are composed of more than one gene encoded adjacent to each other, a genomic neighborhood analysis suggested that Tiamat is a standalone defense protein (3). When expressed under its native promoter from a low-copy number plasmid, a *Bacillus cereus* Tiamat homolog provided modest defense against *Escherichia coli* phages T5 and T6 (3). Tiamat is a multidomain protein consisting of an N-terminal Hsp90-like ATPase and two domains of unknown function (DUFs), DUF3684 and DUF3883 (**Fig. 1a**). Tiamat-based defense requires an intact ATPase site, since the substitution E44A abolishes protection (3), yet Tiamat’s biochemical activities and the roles of its uncharacterized domains remain unknown.

**Figure 1:**
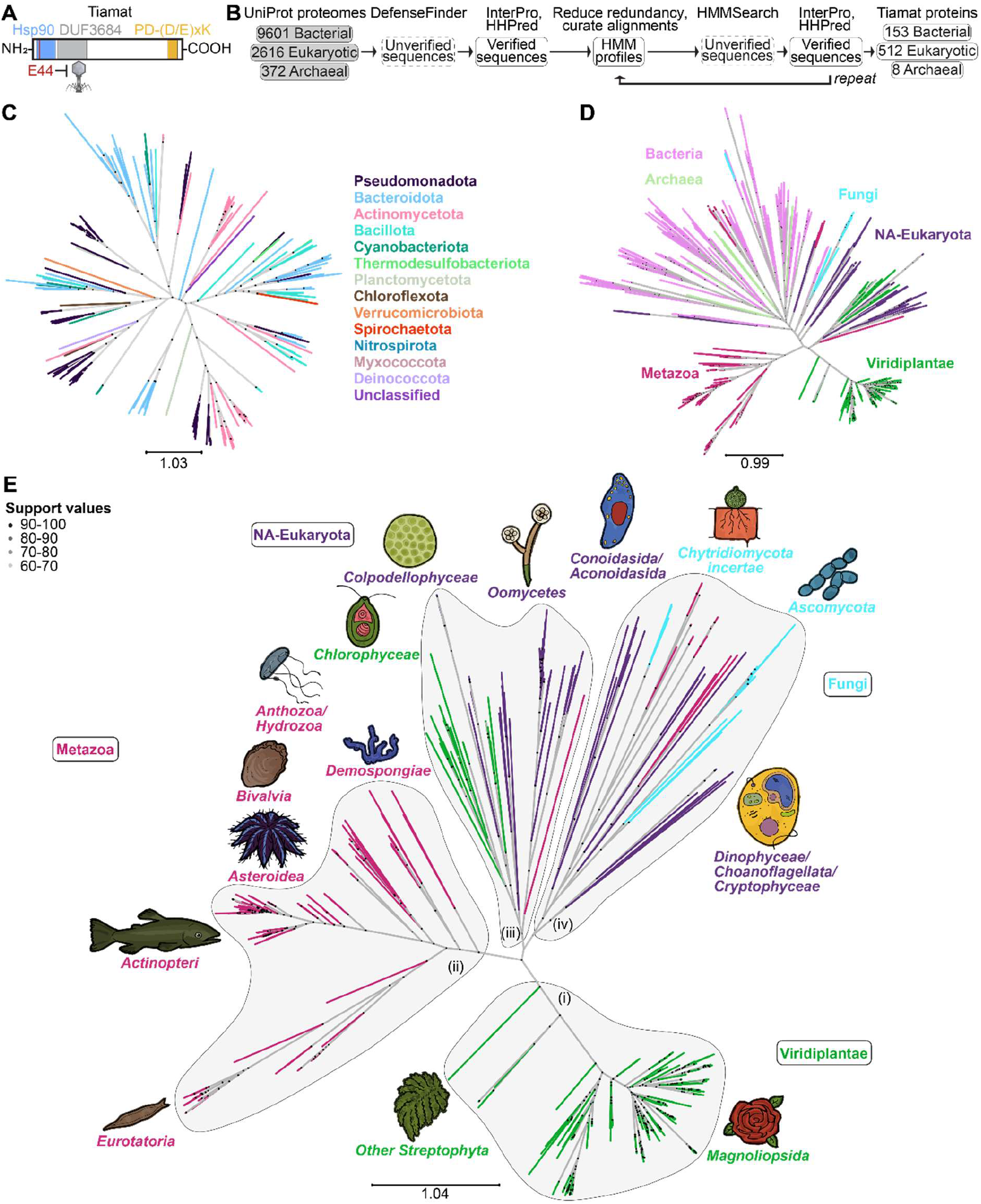
The prokaryotic antiphage protein Tiamat is conserved into eukaryotes. (**A**) 2-D representation of the antiviral Tiamat protein from B. cereus, with summary of known annotations and functional residues (3). (**B**) Workflow to systematically identify and curate Tiamat homolog candidates across bacterial, eukaryotic and archaeal proteomes in the UniProt reference dataset. (**C**) Maximum likelihood phylogenetic tree of bacterial Tiamat proteins generated using IQ-Tree 2. Tree branches are colored according to Phylum-level classifications. Support values are plotted as circles. (**D**) Maximum likelihood phylogenetic tree of bacterial, archaeal and eukaryotic Tiamat proteins generated by IQ-Tree 2. The tree branches are colored by Domains for Bacteria and Archaea, and by Kingdom for Eukaryota. “NA-Eukaryota” represents organisms that are not ranked at the Kingdom level by NCBI, and which here consists of many Oomycota, Apicomplexa, Chromists and two Choanoflagellates. (**E**) Maximum likelihood phylogenetic tree of eukaryotic Tiamat proteins generated using IQ-Tree 2. Tree branches are colored according to Kingdom-level classifications (78). The predominant positions of Classes are indicated with named illustrations. The models chosen for illustrations, listed clockwise, are as follows, *Magnoliopsida – Rosa chinensis* (China rose), Other *Streptophyta – Selaginella moellendorffii* (Lycophyte), *Eurotatoria – Rotaria sp*. (Bdelloid rotifer), *Actinopteri – Oncorhynchus tshawytscha* (Chinook salmon), *Asteroidea – Acanthaster planci* (Crown-of-thorns starfish), *Bivalvia – Magallana gigas* (Pacific oyster), *Anthozoa/Hydrozoa – Clytia hemisphaerica* (jellyfish and polyp), *Demospongiae – Amphimedon queenslandica* (sponge), *Chlorophyceae – Edaphochlamys debaryana* (algae), *Colpodellophyceae – Vitrella brassicaformis* (algae), *Oomycetes – Saprolegnia diclina* (water mold), *Conoidasida/Aconoidasida – Toxoplasma gondii* (single-celled parasite), *Chytridiomycota incertae – Neocallimastix californiae* (rumen fungus), *Ascomycota – Scytalidium dimidiatum* (fungus), *Dinophyceae/Choanoflagellata/Cryptophyceae – Guillardia theta* (algae).

The function of Tiamat’s DUF3684 is unclear, though the InterPro entry for this domain (IPR022155) reveals it frequently occurs C-terminal to an Hsp90-like ATPase domain, typically exceeds 1000 residues, and is predominantly found in eukaryotic proteins (44, 45). The ATPase-DUF3684 fusions frequently contain a C-terminal metallopeptidase domain, and these proteins sometimes have N-terminal nuclear localization signals. Remote homology searches place DUF3883 in the PD-(D/E)xK nuclease superfamily, and DUF3883 is commonly fused to helicases, methyltransferases, protein kinases and transcription factor domains (46). Recently, the isolated DUF3883 domain found in HmaA (of the helicase, methylase, ATPase system called Hma) was found to nick supercoiled DNA, which can exhibit short single-stranded stretches, but did not cleave double-stranded DNA (47–49).

Here we show that homologs of the prokaryotic antiviral protein Tiamat are highly prevalent in eukaryotes, encoded by more than half of green plants, water molds, molluscs, cnidarians, rotifers, placozoans and sponges. Analysis of predicted structures resolves DUF3684 and the unannotated middle region into conserved folded domains, including tetratricopeptide-like (TPR-like) repeats, an undetermined “Fold B” domain, and a metalloprotease-like fold, in place of the S5-fold that binds substrate in other GHL proteins. Biochemical assays show that *B. cereus* Tiamat is an ATPase with single-stranded DNase (ssDNase) activity. Eukaryotic homologs such as the *A. thaliana* developmental regulator NO VEIN (Nov) retain the active-site residues of both catalytic domains (50, 51). Beyond Nov, we show that Tiamat is related to the human protein SACSIN, which is mutated in the neurodegenerative disorder autosomal recessive spastic ataxia of Charlevoix-Saguenay (ARSACS) (52, 53). Tiamat is predicted to dimerize through both its ATPase and nuclease domains, reproducing the clamp architecture of other GHL proteins, with the nuclease taking the place of a dedicated dimerization domain. The nuclease domain itself is frequently replaced by other putative effectors, including helicases, peptidases and nucleic acid-binding domains. Tiamat is also frequently encoded adjacent to helicases, nucleases and methyltransferases. Together these data establish Tiamat as a modular GHL ATPase-nuclease conserved from bacteria to eukaryotes, with eukaryotic homologs that have critical non-defense functions. Whether these eukaryotic homologs also retain an antiviral role or are exapted remains an open question.

## MATERIALS AND METHODS

### Systematic Identification of Tiamat Homologs Across the Tree of Life

Reference and complete proteomes for bacteria (n = 9,601), archaea (n = 372), and eukaryotes (n = 2,616) were downloaded from UniProt (release 2024_03, May 29, 2024) using UniProt’s batch API scripts (54, 55). DefenseFinder v1.2.1 was used to identify homologs of known antiviral defense systems (56). This analysis defined the initial set of candidate Tiamat sequences in all reference proteomes (**Fig. S1 and S2**). Candidate Tiamat sequences were annotated using InterProScan v5.72-103.0-64 with all integrated databases enabled (45). Sequences were retained if they contained the ATPase domain (SUPERFAMILY/Gene3D: Histidine kinase/HSP90-like ATPase superfamily) and the NOV_C domain (Pfam: Protein NO VEIN, C-terminal). The NOV_C domain annotation overlaps with the region previously annotated as DUF3883, and from the Results section onwards is referred to in the text as a PD-(D/E)xK nuclease domain. Sequences lacking either domain were removed from the dataset at this stage. Tiamat sequences from each domain of life (bacteria, archaea, eukaryotes) were grouped and the redundancy of sequences in these groups was reduced by clustering with CD-HIT v4.8.1 using a 95% identity threshold (57). The Tiamat sequence from Bacillus cereus (JGI gene ID: 2641244160) was included in each dataset. Each set of sequences was aligned using MAFFT v7.520 (FFT-NS-2 strategy) (58). The ATPase and NOV_C domain boundaries in the *B. cereus* Tiamat were determined via HHPred (accessed 2025-03) using the PDB_mmCIF70, Pfam-A, CATH-S40, and UniProt-SwissProt databases, and used as a reference for trimming the alignments to the sequence coordinates spanning the ATPase to NOV_C regions (59). Gappy regions in the alignments were removed using ClipKit v2.3.0 with the smart-gap mode (60). Trimmed alignments were used to build profile HMMs (Hidden Markov Models) for each domain of life using HMMER v3.3.2 (hmmbuild) (61). These profiles were searched against their respective reference proteome sets using hmmsearch with default parameters. For bacterial and eukaryotic Tiamat proteins, InterProScan was used to filter out resulting protein sequences that lacked ATPase or NOV_C domains (45). For archaeal Tiamat proteins, however, both InterProScan and HHPred were used for domain annotation and to remove protein sequences that lacked ATPase or NOV_C domains (59). This search process was repeated iteratively until no new sequences with both domains were recovered (**Fig. S1**). Due to the small number of archaeal hits found (n = 6), a separate strategy was employed to identify additional archaeal Tiamat sequences. First, bacterial sequences identified by the approach above were aligned, and the regions of the alignment corresponding to the ATPase and NOV_C domains were extracted. Second, those regions were used to build custom HMMs for the individual domains, which were used to do an hmmsearch against the archaeal proteomes (61). Finally, sequences lacking either domain were excluded from the final dataset of archaeal Tiamat candidate sequences.

### Phylogenetic Analyses

To assess the phylogenetic relationships between Tiamat sequences, the sequence from the ATPase domain through to the C-terminal NOV_C domain inclusive was extracted and aligned with MAFFT v7.520 (E-INS-i strategy) (58). The alignments were trimmed using the gappyout method on TrimAl to remove gappy regions (62). Next IQ-Tree 2 was used to build maximum likelihood phylogenetic trees for either bacterial sequences alone, eukaryotic sequences alone, or a combination of bacterial, archaeal and eukaryotic sequences. In all cases, the substitution models were selected by IQ-Tree 2’s model selector (Q.pfam+F+R8 for Bacteria, JTT+F+R10 for Eukaryotes, JTT+F+R10 for combined dataset), and branch support was assessed using 1,000 Ultrafast bootstraps (63). Trees were visualized and annotated with TreeViewer (64). Protein BLASTs of eukaryotic sequences from clades 4, 5 and 6 (which contained mixtures of bacterial and eukaryotic sequences) were performed to determine if bacterial proteins could be independently identified as close homologs. BLASTs were performed using the blastp suite hosted at blast.ncbi.nlm.nih.gov, against the Non-redundant protein sequences (nr) database, and with “Max target sequences” set to 5000, all other parameters kept default.

To assess the phylogenetic relationship between Tiamat proteins, Tiamat-like proteins, classical GHL ATPase family members, bacterial MORC proteins, paraMORC proteins, and putative antiphage Hsp90-like proteins we collected and curated representative sequences of each group of proteins originating from archaea, bacteria and eukaryotes from previously published lists or from UniProt and NCBI databases directly (**Data S1**) (18, 20, 21, 65). The ATPase domains were annotated with the InterProScan webserver (accessed: 08/17/2026) (45). The CATH-Gene3D (G3DSA:3.30.565.10) annotation was used as the primary reference annotation, since it identified ATPase domains most consistently in the analyzed proteins (152 of 169 proteins). The amino acid coordinates of the ATPase domain for proteins which were instead annotated by other HMM profiles were calibrated to the G3DSA coordinates using empirically determined boundaries for the analyzed dataset of proteins, to ensure that the sequences extracted from each protein span approximately equivalent regions. GHL ATPase domains of each protein were extracted along with 50 upstream and 200 downstream flanking residues. These boundaries were determined empirically through successive sequence extraction, alignment, and trimming iterations. The final boundaries encompass the Bergerat fold and the adjacent C-terminal transducer or transducer-like region. Including greater than 50 residues N-terminal to the ATPase domain introduced columns that were below a 30% gap threshold. A region 100-150 residues C-terminal to the ATPase domain was just as, or slightly better occupied, than the ATPase domain itself and was populated by all protein families. Beyond 200 residues C-terminal of the ATPase domain we saw diminishing returns in the number of ungapped columns gained in multiple sequence alignments. Several proteins in the dataset are SACSIN and Tiamat-like proteins that appear to be SACSIN homologs, which have multiple ATPase domains. For these proteins, the most N-terminal ATPase domain was extracted. Sequences were aligned with the MAFFT webserver (accessed: 08/17/2026; E-INS-i strategy) (58). Sequences were subsequently trimmed locally with trimAl v1.5.rev1 using a 30% gap threshold (-gt 0.3) (62). Next, IQ-Tree v3.1.3 was used to build a phylogenetic tree using a substitution model selected by IQ-Tree’s ModelFinder (Q.PFAM+F+I+R7), and branch support was assessed using 1,000 ultrafast bootstraps with NNI optimization (66, 67). Trees were visualized and annotated with TreeViewer (64).

### Identification of Conserved Protein Folds and Active Sites

Representative Tiamat protein sequences were determined for each clade for further structural analyses and fold determination. First, the core region of each Tiamat homolog, spanning the ATPase domain to the NOV_C domain, was extracted, aligned with MAFFT v7.520 (E-INS-i strategy) and used to build clade-specific HMMs using hmmbuild (58, 61). The resulting HMM was used to search the same set of sequences, and results were ranked by score. The rank 1 sequence of each clade was chosen as the representative. Structures for selected sequences were predicted using the AlphaFold3 webserver, including only the region spanning from the start of the ATPase domain to the end of the NOV_C domain (68). TED (The Encyclopedia of Domains) was also used to annotate protein domains between the ATPase and NOV_C domain, by examining the best AFDB50 matches from Foldseek (69, 70). Individual putative domains from each clade representative were extracted and compared using FoldMason, which informed an iterative improvement of domain boundaries, and enabled the clustering of similar folds (**Fig. S9**) (71). Clade 8 sequences were excluded from this structure-based search because the proteins in this clade have many insertions of low complexity regions (e.g., poly-Ala, –Gly, –Gln, –Pro, –Ser) which decreased the confidence of predicted structures and inhibited the identification of domain boundaries. Foldseek identified high probability matches of the ATPase and NOV_C domain structures to experimentally solved structures of ATPase domains and nuclease domains (i.e., probability >0.3), but did not identify high probability matches for other predicted domains (e.g., probability <0.2). Dali was therefore applied to identify matches to representatives of the domains found between the ATPase and NOV_C domain. Dali identified likely structural matches for two types of domains, TPR-like folds and a metalloprotease-like fold (i.e., z-score >4). To further compare all putative protein domains, the predicted folds were compared using TM-align (**Fig. S9**) (72). InterProScan v5.72-103.0-64 and HHPred were used to identify additional putative protein domains (45, 59). Subcellular localization of candidate Tiamat homologs was predicted using ProtGPS (available at https://github.com/pgmikhael/protgps) with default parameters, and a confidence threshold of 0.8 was used to minimize false positives (73).

### Identification of Tiamat and Tiamat-like proteins and clade assignment

After an initial analysis of UniProt reference proteomes (**Fig. 1, 2, 3**), all complete RefSeq proteomes from bacteria (n = 6,259), archaea (n = 390) and eukaryotes (n = 2,256) were downloaded from NCBI and searched via HMMER v3.3.2 (hmmsearch) using the final curated profile HMMs generated from UniProt-derived Tiamat proteins. This HMM search recovered a list of 10,229 bacterial, 130 archaeal and 17,820 eukaryotic candidate Tiamat sequences. Candidate proteins were annotated using InterProScan v5.77-108.0 with the Pfam, SUPERFAMILY, SMART, and Gene3D databases enabled (45). A collection of strictly curated Tiamat proteins was generated which contained both the ATPase domain and PD-(D/E)xK nuclease domain (annotated as “Histidine kinase/HSP90-like ATPase domain” and “NO VEIN C-terminal domain” by InterProScan). The list of RefSeq-derived Tiamat sequences was combined with the original list of UniProt-derived Tiamat sequences, and redundancy was reduced by clustering with CD-HIT v4.8.1 using a 95% amino acid identity threshold, yielding a list of strictly curated and non-redundant Tiamat sequences, 918 of which were from eukaryotes, 221 of which were from bacteria, and 21 from archaea (**Fig. 4, S15, Table S1 and Data S2**). A clade-specific HMM classification strategy was used to assign the Tiamat proteins identified from RefSeq sequences to clades previously determined from UniProt-derived sequences (**Fig. 3, Table S1**).

**Figure 2.**
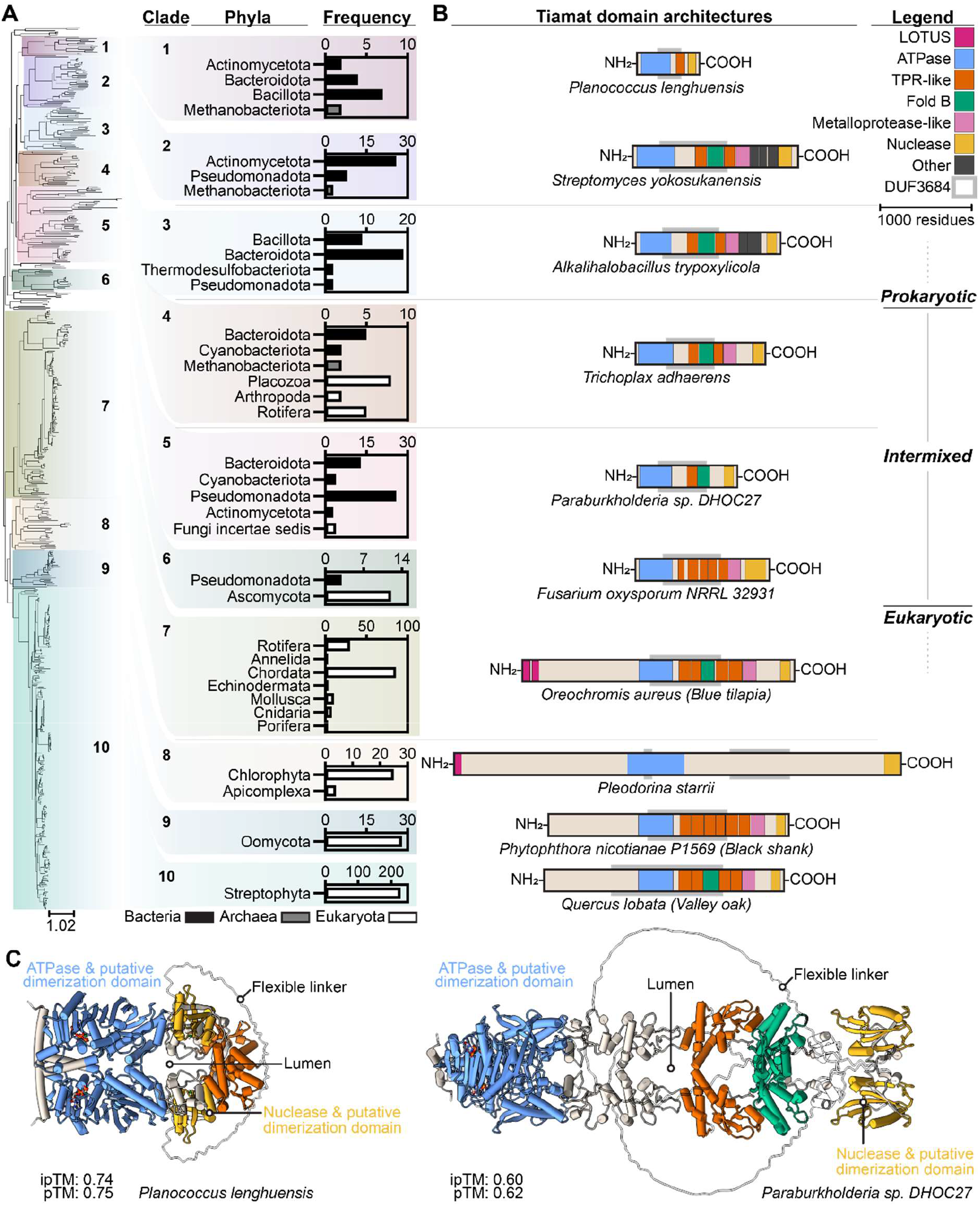
Tiamat homologs share previously undescribed domains conserved across the tree of life. (**A**) Maximum likelihood phylogenetic tree of bacterial, archaeal and eukaryotic Tiamat proteins generated using IQ-Tree 2, depicted in Figure 1 but here displayed with a midpoint root (left). The frequency of Phyla in clades of Tiamat proteins (right). For clarity, only Phyla that had more than one representative are plotted. (**B**) 2-D representations of example Tiamat protein architectures, centered on their ATPase domains. A sequence-based approach was used to determine a representative Tiamat sequence for each clade (**Figure S5**). LOTUS domains were identified by InterProScan (45). The extension of the ATPase fold was identified by The Encyclopedia of Domains (TED) (69) and by comparisons to experimental structures (**Figure S6 and S7**). The PD-(D/E)xK nuclease domain (previously annotated as DUF3883 or NOV_C domain) was identified by Foldseek (70) and by comparisons to experimental structures (**Figure S7 and S8**). TPR-like, Fold B, metalloprotease-like, and “Other” domains were identified with TED, HHPred, and Dali (59, 69, 82). TM-score was used to quantify the similarity of representatives of each domain to each other (**Figure S9**). (**C**) Structures of representative Tiamat proteins from Clades 1 and 5, predicted as dimers in the presence of two ATPs and six Mg^2+^ ions with the AlphaFold 3 webserver. The ATPase and nuclease domains of most representative Tiamat proteins are predicted to form stable dimers (**Figure S6 and S8**). The intermediate domains are predicted to be flexible or to hinge (**Figure S10 and S11**). Often the nuclease domain is tethered by a highly flexible linker, and moves relative to the ATPase. Residues and atoms with a pLDDT score lower than 0.5 are shown in 100% transparency.

**Figure 3.**
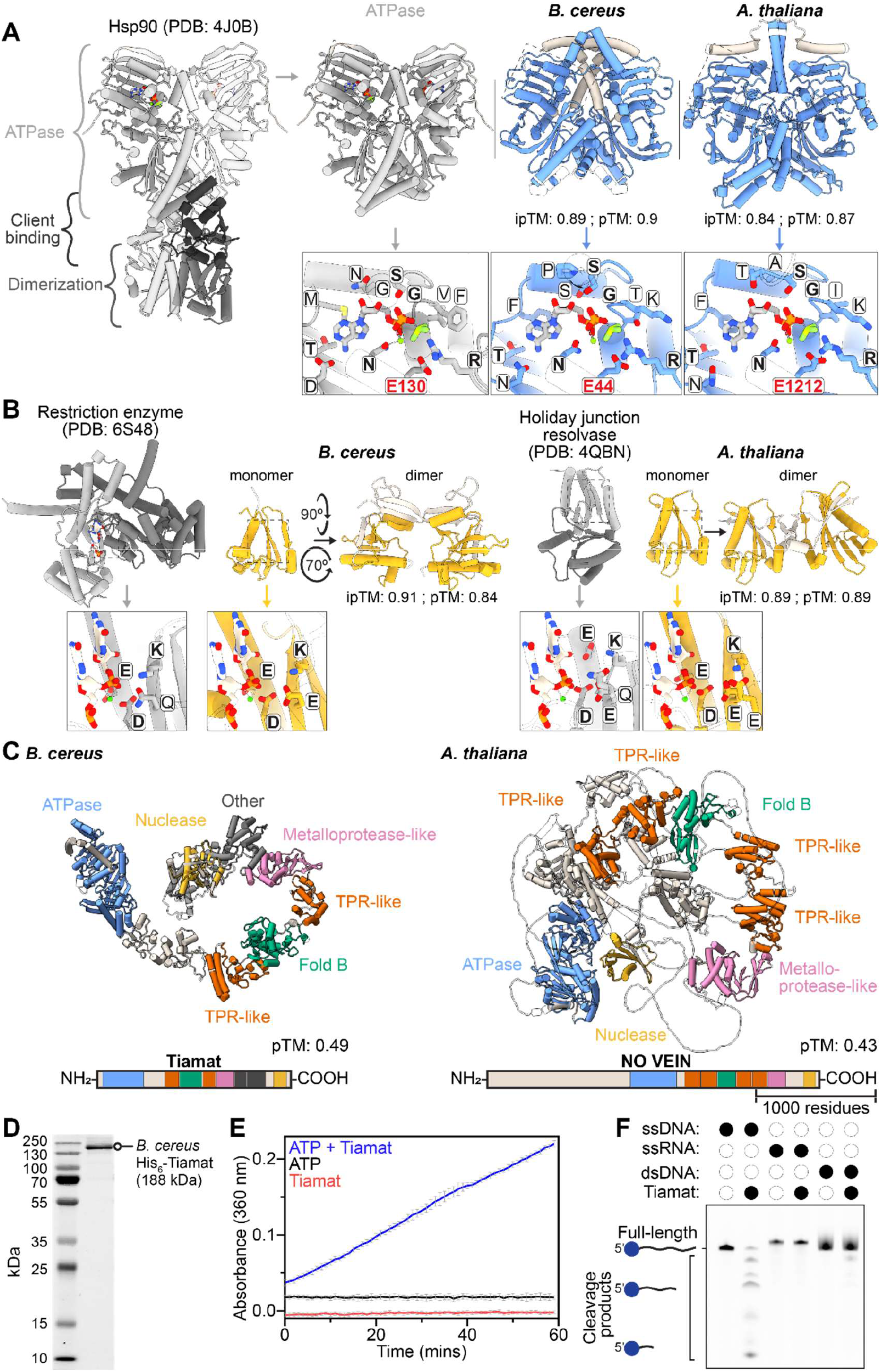
B. cereus Tiamat is an ATPase and ssDNase with catalytic residues conserved through to A. thaliana Nov protein. (**A**) A previously determined structure of zebrafish mitochondrial Hsp90 with ADP and BeF_3_ (greyscale, left, PDB: 4J0B) (87), shown alongside the AlphaFold3-predicted structures of dimers of the putative ATPase domains from B. cereus Tiamat and A. thaliana Nov proteins (blue). One monomer of PDB: 4J0B is shown semi-transparently. An overview (top) and a zoomed-in view of the active sites (bottom) are shown. 20-42 residues N-terminal to the domain were included (beige) and assist in forming the dimerization interface between the ATPase domains. To view how ATP might fit into the putative ATPase active sites, dimers of the B. cereus and A. thaliana domains were aligned to the Hsp90 structure and then all components of that structure were hidden except ADP, BeF_3_, and the Mg^2+^ ion. Active site residues are labeled. Residues conserved across known and putative active sites are bolded. A key active site residue required for ATP hydrolysis, and which has been shown to be important for B. cereus Tiamat antiviral activity, is labelled in bold and red (3). (**B**) Previously determined structures of dimeric bacterial restriction enzymes and Holliday junction resolvases (greyscale, PDB: 6S48 and 4QBN) are shown next to the putative PD-(D/E)xK nuclease domains from B. cereus Tiamat and A. thaliana Nov proteins (yellow). Predictions of monomers (left) and dimers (right) are shown. The C-terminal 12-46 residues (beige) appear necessary for prediction of stable nuclease dimers. An overview (top) and zoomed-in view of the active sites (bottom) are shown. A dinucleotide and metal co-factor from the 6S48 structure are shown overlaid on all domains similar to panel A. Active site residues are labelled. Residues that are conserved across known and putative active sites are bolded. (**C**) Full-length predicted structures of monomeric B. cereus Tiamat and A. thaliana Nov proteins are shown, alongside a 2-D scheme of the protein structures. The proteins are colored in beige, with putative biochemically active domains and annotated folds given additional colors. Regions of the predicted protein structures with a pLDDT score less than 0.5 are shown in 100% transparency. Predicted structures of full-length dimeric B. cereus Tiamat and A. thaliana Nov are shown in **Figure S11**. (**D**) SDS-PAGE gel of purified B. cereus Tiamat protein. (**E**) ATPase activity monitored using the EnzChek Phosphate Assay Kit. The blank-subtracted average of 3 replicates for reactions containing both ATP and Tiamat, or negative-controls containing either ATP alone or Tiamat alone are shown, along with standard deviations (**Fig. S12**). (**F**) Nuclease assay with 54-nt Cy5-labeled, ssDNA, ssRNA or dsDNA substrates in the presence or absence of Tiamat, resolved on a denaturing polyacrylamide gel. One replicate of two is shown (**Fig. S12**).

**Figure 4.**
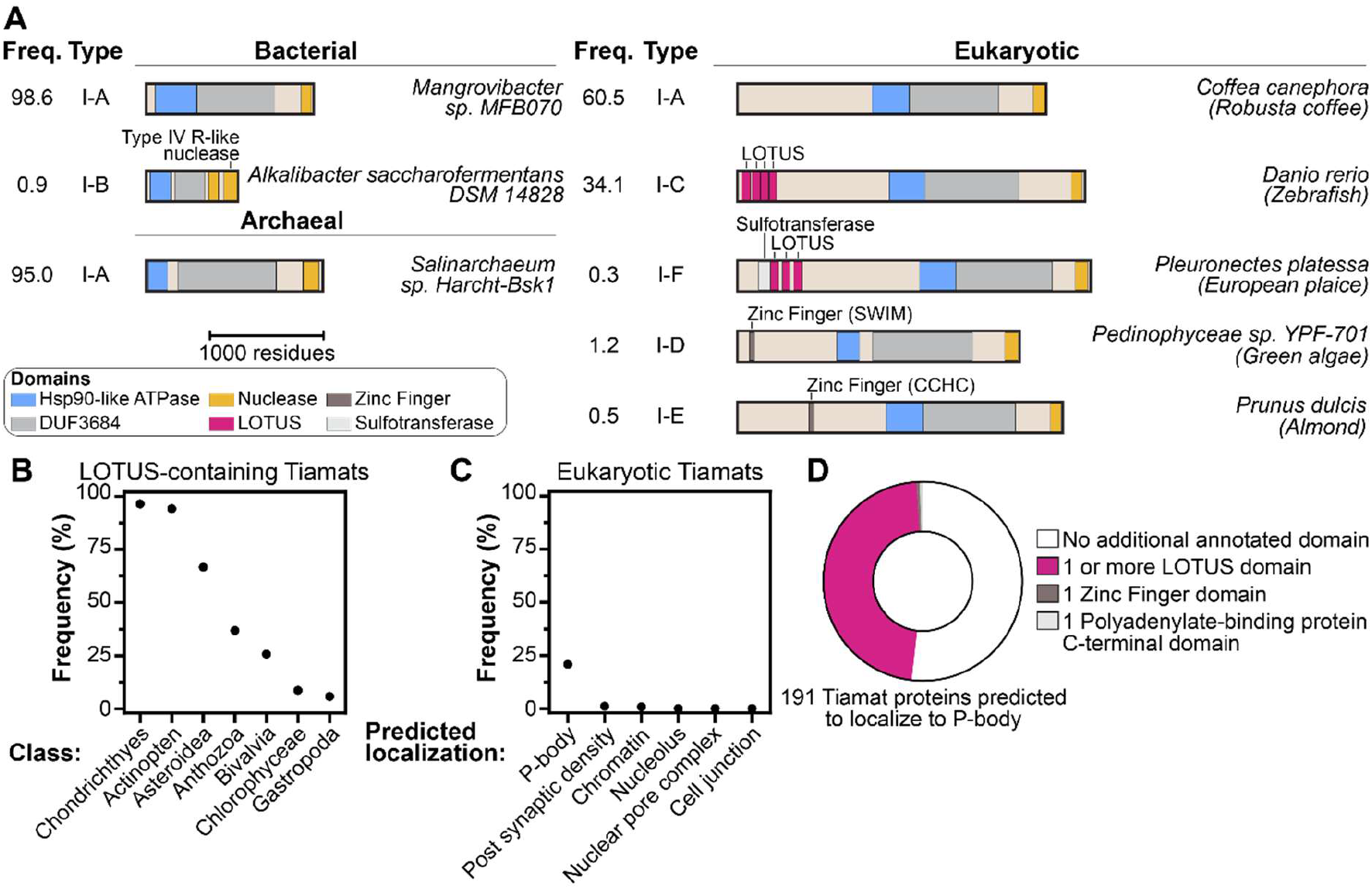
Eukaryotic Tiamat proteins frequently contain LOTUS domains and are predicted to localize to processing bodies. (**A**) The percent frequency of Tiamat proteins, identified in UniProt and RefSeq datasets, with unique domain architectures as annotated by InterProScan (45) and HHPred (59) are shown. For clarity, only domains that occurred in at least 2 bacterial or archaeal proteins, or 3 eukaryotic proteins are depicted here (**Data S5, S6**). Unless otherwise noted, the blue, grey and gold domains correspond to previously described ATPase, DUF3684 and PD-(D/E)xK nuclease domains. Tiamat homologs were assigned subtype classifications based on unique sequence-based annotations. Tiamat sequences originated from UniProt reference proteome and RefSeq databases. (**B**) The Class distribution of Tiamat proteins which are predicted to contain a LOTUS domain. For clarity, percent frequencies are shown only for taxonomic classes that have 4 or more Tiamat proteins. (**C**) 918 eukaryotic Tiamat protein sequences were analyzed with ProtGPS (73) to predict their subcellular compartment localization (**Data S7**). The percentage of Tiamat proteins with a given predicted localization is plotted. For clarity, only subcellular localizations with frequencies greater than 0% are plotted (**Fig. S14**). (**D**) Plot of domain annotations in Tiamat proteins predicted to localize to the P-body. Tiamat proteins predicted to localize to the P-body are enriched in those annotated with LOTUS domains.

Sequences that lacked either the ATPase or PD-(D/E)xK domains but which might play an analogous role in antiviral defense were curated in multiple steps. Sequences lacking either of these two domains, but which had high similarity to an HMM derived from Tiamat proteins (**Fig. S15**), were collated to generate an initial list of 21 archaeal, 497 bacterial and 1,948 eukaryotic sequences (**Data S3, S4**). InterProScan-based annotations of these “Tiamat-like” sequences were used to sort and cluster sequences (**Data S5, S6**) prior to additional manual curation of related clades of sequences using HHPred. Sequences with a header containing the descriptor “(fragment)” or “LOW QUALITY PROTEIN”, and those shorter than 900 residues were removed from further analyses.

### Neighborhood analysis

RefSeq IDs were converted to UniProt IDs using the UniProt ID mapping tool (https://www.uniprot.org/id-mapping). FASTA files of archaeal and bacterial strictly curated Tiamat proteins or Tiamat-like proteins were submitted to the EFI-EST (Enzyme Function Initiative-Enzyme Similarity Tool) webserver to cluster potentially isofunctional proteins together and to retrieve the genome context for each protein (74). The output from EFI-EST was submitted to the EFI-GNT (Enzyme Function Initiative-Genome Neighborhood Tool) to generate genome neighborhood networks and retrieve genome neighborhood diagrams, consisting of 10 genes upstream and downstream of the Tiamat or Tiamat-like protein.

### AlphaFold3 predictions

Most AlphaFold3 predictions were conducted via the webserver (https://alphafoldserver.com/) which has a 5,000 token limit. Homodimers of many eukaryotic representatives are larger than 5,000 residues in length. Therefore, the predictions of full-length representative Tiamat homodimers from each clade were performed using a local copy of AlphaFold3 run on the University of Pennsylvania’s high-performance computing and artificial intelligence cluster “Betty”. A custom cache was built to increase the token limit to 8,300. For predictions performed on “Betty”, three specified seeds were used for each complex (70507726, 1592465280, 1640934929) and the most confident of these three results was included in this manuscript (**Fig. S11**).

### Expression and purification of B. cereus Tiamat

BL21(DE3) cells (Invitrogen) harboring plasmids encoding His_6_-tagged B. cereus Tiamat were grown in three liters of LB media supplemented with ampicillin at 37 °C to OD₆₀₀ of roughly 0.4, cooled to 18 °C, and induced with 0.25 mM IPTG for roughly 16 hours while shaking at 150 RPM. Cells were harvested (4,500 × g, 15 min, 4 °C), flash-frozen, and resuspended in lysis buffer (20 mM HEPES–KOH pH 7.4, 500 mM NaCl, 10 mM imidazole, cOmplete protease inhibitor; Roche). Lysates were generated by sonication (QSONICA Q700A-220 sonicator with ½” tip; amplitude 40%, 2s ON/8s OFF, 5 minutes total ON time) and clarified by centrifugation (40,000 × g, 30 minutes, 4 °C).

Supernatants were applied to a HisTrap FF column (Cytiva), washed with five column volumes washing buffer (20 mM HEPES–KOH pH 7.4, 500 mM NaCl, 20 mM imidazole), then washed with two column volumes of high salt washing buffer (20 mM HEPES–KOH pH 7.4, 1.2 M NaCl, 20 mM imidazole), followed by one column volume of regular washing buffer until the A280 stabilized. The protein was eluted by increasing the imidazole concentration in one step (20 mM HEPES–KOH pH 7.4, 500 mM NaCl, 250 mM imidazole). After pooling the fractions containing the protein of interest the salt concentration was lowered to 250 mM NaCl by diluting with 20 mM HEPES-KOH pH 7.4. Subsequently, samples were applied to a Heparin HiTrap HP column (Cytiva). The column was washed using a low salt washing buffer (20 mM HEPES–KOH pH 7.4, 250 mM KCl, 0.5 mM TCEP) until the A280 was roughly zero, and eluted by applying a linear gradient from 250 mM to 1 M KCl, over 60 mL. Peak fractions containing the protein of interest were pooled, concentrated using a centrifugal filter (Amicon), and loaded on a Superose 6 HiLoad 16/600 column (Cytiva) for size exclusion chromatography (SEC). The protein was eluted with SEC buffer (20 mM HEPES–KOH pH 7.4, 500 mM KCl, 0.5 mM TCEP) and the fractions containing the protein of interest were combined. Subsequently, the sample was concentrated again, aliquoted, flash frozen in liquid nitrogen, and stored at –70 °C.

### ATPase assay

ATP hydrolysis by *B. cereus* Tiamat was monitored using the EnzChek Phosphate Assay Kit (Invitrogen) according to the manufacturer’s protocol. The provided reaction buffer was adapted to contain 150 mM KCl and 10 mM MgCl₂, with 1 mM ATP and 100 nM Tiamat. Reactions were assembled in 96-well plates and incubated at 25 °C, and phosphate release was quantified by reading absorbance at 360 nm in a CLARIOstar plate reader (BMG). Reactions were performed as technical triplicates (**Fig. S12**).

### Nucleic acid cleavage assay

Cleavage assays were performed using *B. cereus* Tiamat and Cy5-labeled nucleic acid substrates (DNA: JBO-0081 with complementary strand JBO-0062; RNA: JBO-0084). Final concentrations of each component were 1 µM *B. cereus* Tiamat monomer (0.5 µM dimer), 100 nM labeled nucleic acid, 20 mM HEPES pH 7.4, and 250 mM KCl. Oligonucleotide sequences are as follows, JBO-0081: Cy5-TGAAAAAGGAAGAGTATGAGTATTCAACATTTCCGTGTCGCCCTTATTCCCTTT; JBO-0062: AAAGGGAATAAGGGCGACACGGAAATGTTGAATACTCATACTCTTCCTTTTTCA; JBO-0084: Cy5-UGAAAAAGGAAGAGUAUGAGUAUUCAACAUUUCCGUGUCGCCCUUAUUCCCUUU.

Following 1 hour of incubation at 37 °C the reactions were quenched by the addition of 0.4 µl 0.5 M EDTA and 1 µl Proteinase K and incubated for 30 min at room temperature in the dark. Samples were mixed with 1× denaturing loading dye, heated, and resolved on 15% TBE urea–PAGE gels. Gels were pre-run for 30 min and electrophoresed at 180 V for 1 hour at 47 °C. Fluorescence was detected using a Chemidoc imager (Bio-Rad). Reactions were performed in duplicate using independent master mixes (**Fig. S12**).

## RESULTS

### Homologs of the antiviral protein Tiamat are abundant in eukaryotes

To determine whether eukaryotes encode homologs of prokaryotic antiphage defense systems, we searched 2,616 reference eukaryotic proteomes from the UniProt database with DefenseFinder, a tool that uses a library of Hidden Markov Model (HMM) profiles to identify distant homologs of known prokaryotic antiviral defense proteins (**Fig. 1b, S1, and S2**) (54, 56, 75, 76). UniProt reference proteomes provide a representative cross-section of taxonomic diversity and high coverage of protein-space for a given species. In accordance with previous observations, the two most abundant eukaryotic families were Argonaute and Viperin, with 6,637 and 890 sequences respectively (**Fig. S2**) (10, 77). To our surprise, the third most abundant family comprised homologs of the antiphage protein Tiamat, with 359 sequences identified at this stage, prior to curation and redundancy filtering (**Fig. 1a,b**) (3). Below Tiamat, family sizes fell by an order of magnitude (28 or fewer homologs), and these families were not analyzed or verified further (**Fig. S2**).

Many putative homologs recovered by DefenseFinder lacked either the GHKL ATPase domain, which is required for antiphage activity, or the NO VEIN C-terminal domain (NOV_C) (3). The NOV_C annotation overlaps the previous DUF3883 annotation, and both place the domain in the PD-(D/E)xK phosphodiesterase superfamily, whose enzymes function in DNA replication, phage restriction, DNA repair and tRNA-intron splicing (**Fig. 1a**) (46). We therefore refer to DUF3883 and NOV_C hereafter as the PD-(D/E)xK, or nuclease, domain. Variants lacking either domain may well be antiviral, but we conservatively required both in our search, retaining the proteins most likely to share the biochemical activities of the validated antiphage Tiamat from B. cereus (3). Two iterations of custom HMM building and searching saturated the search, yielding a strictly curated set of 153 bacterial, 8 archaeal and 512 eukaryotic non-redundant sequences (**Fig. 1b, S1, Table S1 and Data S2**).

Within this curated set of identified homologs, the bacterial antiphage protein Tiamat is surprisingly rare among prokaryotic phyla. It occurs in less than 5% of prokaryotic proteomes, and most of those encode a single Tiamat (**Fig. S2b,c**). In certain eukaryotic phyla, by contrast, Tiamat homologs are highly prevalent. For example, 78% of proteomes from Streptophyta (green plants) encode a Tiamat homolog, and 60% of these encode more than one. Every rotifer, placozoan and sponge proteome analyzed contains a Tiamat homolog, although this observation may be skewed by the low sample size of 10, 2 and 2 proteomes respectively (**Fig. S2b,c**). Water molds, cnidarians and molluscs also carry Tiamat homologs in 50–80% of proteomes. Tiamat homologs are comparatively rare in the chordates (13% of proteomes), though still more common than in any bacterial phylum. Strikingly, all chordate Tiamat homologs come from aquatic species, including 80 from ray-finned fish, 3 from lancelets, 2 from cartilaginous fishes and 1 from a bichir (**Fig. S2c, Data S2**). No tetrapod proteome analyzed encodes a Tiamat homolog.

Extending this HMM-based search to the RefSeq complete genomes database increased the number of Tiamat homologs identified by 1.4–2.6-fold per domain of life, to 221 bacterial, 21 archaeal and 918 eukaryotic sequences. The phylogenetic and structural analyses below use the UniProt-derived data (**Fig. 1, 2, 3**), while the analyses of terminal extensions, localization, domain shuffling and gene neighborhoods use the combined UniProt– and RefSeq-derived data (**Fig. 4, 5, 6 and Data S2 and S3**).

**Figure 5.**
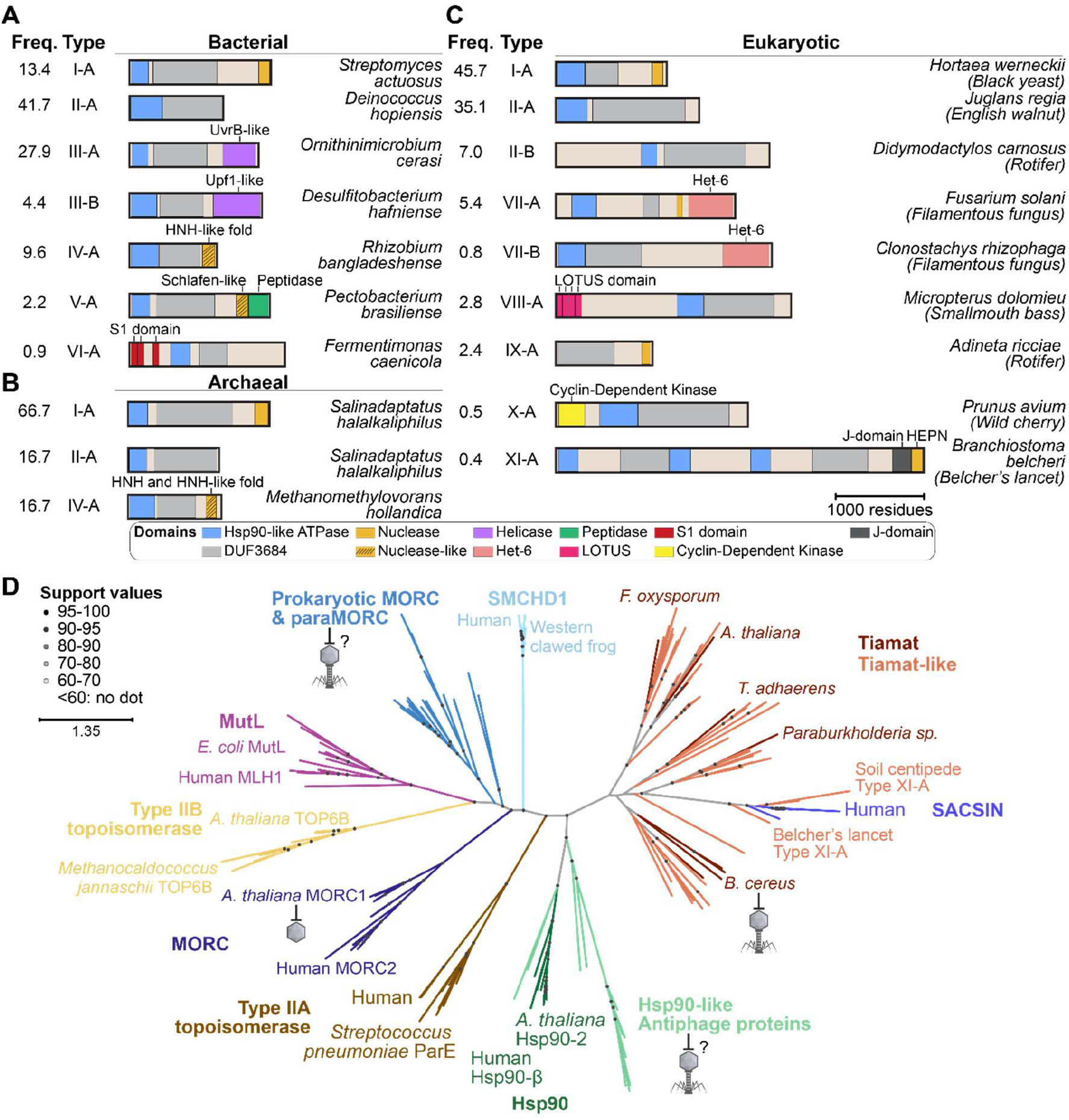
Tiamat-like homologs have evolved via domain shuffling, truncations and fusions and are related to human SACSIN. Tiamat-like proteins from (**A**) bacteria, (**B**) archaea and (**C**) eukaryotes that did not pass initial sequence curation either share domain architectures with strictly curated Tiamat proteins, possess novel domains in place of the PD-(D/E)xK nuclease domain, or are truncated. Domain types or activities are color-coded. Tiamat-like proteins were assigned type and subtype classifications based on unique sequence-based annotations. The frequency of Tiamat sequences within a given architecture, calculated as a percent out of all bacterial, archaeal or eukaryotic Tiamat-like sequences is shown. Tiamat sequences originated from UniProt reference proteome and RefSeq databases. (**D**) Unrooted maximum likelihood phylogenetic tree of 169 bacterial, archaeal and eukaryotic GHL ATPase proteins generated by IQ-Tree 3. Branches are colored by protein family. Protein family is labelled in bold font, example proteins are labelled in normal and italic font. Node labels show UFBoot support values from 1000 replicates. The GHL superfamily divides into a Tiamat, Tiamat-like and SACSIN clades on one side, and a clade containing all classical GHL families, and other antiphage GHL ATPases on the other side. The scale bar shows substitutions per site. Schemes for antiphage and antiviral activity are depicted for B. cereus Tiamat, A. thaliana MORC1, and for putative antiphage Hsp90-like proteins, and prokaryotic MORC and paraMORC proteins.

**Figure 6.**
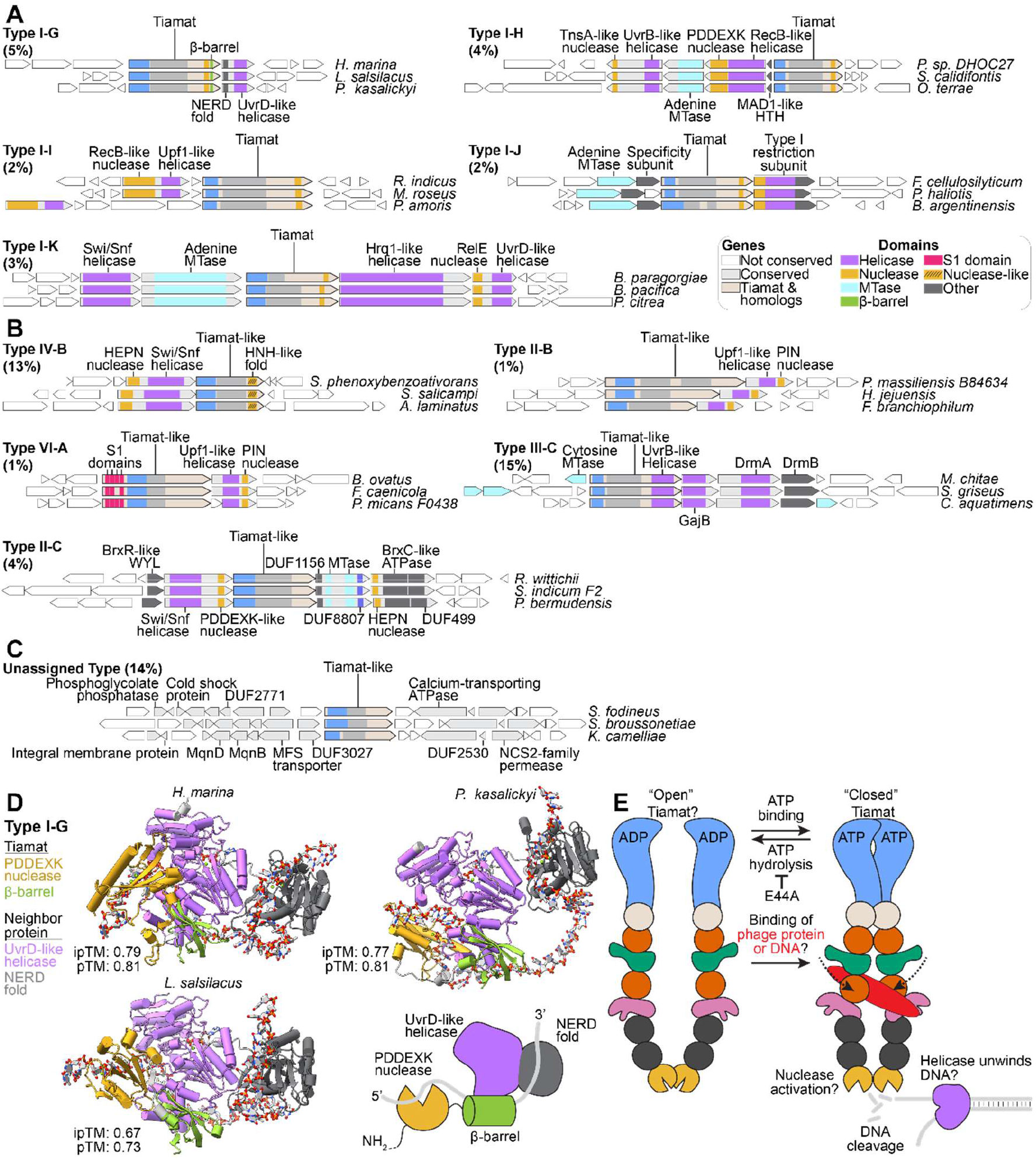
Tiamat and Tiamat-like proteins are frequently encoded near helicases, nucleases and DNA methylases. (**A**) Gene-neighborhood analysis of strictly curated prokaryotic Tiamat homologs. Tiamat and associated neighbor proteins were assigned subtype classifications based on unique sequence-based annotations. Schemes of Tiamat and associated genes are shown in a minimum of 3 different genetic contexts. Subtype and percent frequency of neighborhood occurrence out of all strictly curated bacterial Tiamat proteins are noted. Percentages were rounded to the nearest whole number. (**B**) Gene-neighborhood analysis of prokaryotic Tiamat-like homologs. Subtype and percent frequency of neighborhood occurrence out of all bacterial Tiamat-like proteins are noted. (**C**) A group of Tiamat-like proteins is associated with vitamin synthesis and membrane transport genes. (**D**) Predicted structures of the C-terminus of Type I-G Tiamat proteins with the associated NERD-helicase protein, in the presence of one ATP, 40-nt long T-rich DNA, and five Mg^2+^ ions. In all cases the unique β-barrel fused to the C-terminus of Type I-G Tiamat proteins appears to recruit the NERD-helicase protein by binding the same surface of the helicase (**Fig. S16**). A scheme summarizing the predicted protein-protein and protein-DNA interactions is shown (bottom right). (**E**) Speculative model for antiphage surveillance and activity of Tiamat. Like other GHKL-like ATPases (**Fig. S17**), proteins or DNA may bind to the middle region of an ADP-bound Tiamat in an “open” state. Subsequent exchange of ADP for ATP may trigger dimerization of the ATPase domains and closing of the Tiamat dimer. Capture of phage-derived proteins or DNA may trigger a conformational change that enhances nuclease activity. Tiamat may nick dsDNA in the cell or cooperate with a host or co-encoded helicase to unwind and cleave dsDNA into fragments, to degrade the phage genome specifically or all DNA in the cell. Tiamat may slowly hydrolyze ATP to ADP to relax to an inactive state and open into a conformation amenable to binding targets. The loss of defense that has been previously associated with the E44A mutation in *B. cereus* Tiamat suggests ATP hydrolysis (and the associated opening of the complex), not solely ATP binding, is critical to defense (3, 158, 159).

### Diverse Tiamat homologs bridge the gap between prokaryotic and eukaryotic clades

To analyze the phylogenetic relationships of bacterial, archaeal and eukaryotic Tiamat proteins, we extracted and aligned their core regions, defined here as the sequence from the ATPase to the PD-(D/E)xK domain inclusive. Bacterial Tiamat sequences are highly divergent, and their phylogeny does not recapitulate bacterial taxonomy (**Fig. 1c**). Instead, clades often contain sequences from bacteria in different phyla (**Fig. 1c**). This pattern could indicate that bacterial Tiamat genes are often horizontally transferred between distantly related bacteria, and implies that bacterial Tiamat either does not require phylum-specific host factors for defense, or evolves readily to interface with diverse host proteins, like some CRISPR systems (79–81).

The eukaryotic Tiamat tree, by contrast, shows little intermixing, such that proteins coming from a given taxonomic group often cluster (**Fig. 1e**). Two of the four main branches are taxonomically coherent: (i) a metazoan branch spanning sponges, cnidarians, molluscs, annelids, echinoderms, lancelets, fishes and rotifers, (ii) a branch of Streptophyta (green plants) alone, with many closely related but non-redundant members. While the other two are disparate: (iii) a branch consisting largely of green algae, dinoflagellates and water molds, (iv) a branch exhibiting substantial intermixing, combining fungi, some rotifers, apicomplexans (like the malaria parasite) and cryptophyte algae (**Fig. 1e, Table S1**). The coherence among animals and plants suggests that eukaryotic Tiamat genes have been predominantly inherited vertically since their acquisition into these two taxonomic groups, unlike the frequent horizontal transfer inferred among bacteria.

The combined phylogenetic tree of bacterial, archaeal and eukaryotic Tiamat proteins reveals that a minority of eukaryotic sequences share sequence and structural similarities with prokaryotic Tiamat sequences (**Fig. 1d, 2**). Tiamat proteins cluster into ten main clades, which include three prokaryotic (clades 1–3), three intermixed (clades 4–6), and four eukaryotic (clades 7–10) (**Fig. 2a**). Specifically, several eukaryotic Tiamat sequences from the phyla Ascomycota, Placozoa, Rotifera and Arthropoda have core regions resembling Tiamat proteins from Pseudomonadota, Bacteroidota, Cyanobacteria, Actinomycetota and Methanobacteriota (**Fig. 2a**).

Two independent lines of evidence support the clading between several bacterial and eukaryotic sequences. First, protein BLAST searches seeded with representative eukaryotic sequences from the intermixed clades 4 and 5 retrieve bacterial Tiamat sequences alongside homologs from related eukaryotic species and genera. The best bacterial matches differ by lineage. The spider *Trichonephila inaurata madagascariensis* matches the endosymbiont *Candidatus* Rickettsiella viridis, the placozoan *Trichoplax adhaerens* matches *Clostridium* spp. and *Paenibacillus* spp., the rotifer *Adineta ricciae* matches *Ginsengibacter* spp., and the anaerobic fungus *Neocallimastix californiae* matches *Mangrovibacterium marinum* (query coverage 78–86%, sequence identity 27–46%). However, BLASTs seeded with representative clade 6 sequences did not retrieve confident bacterial matches, for example *Symbiodinium* spp. matched weakly to *Nostoc* spp. (query coverages 21%, sequence identities 28%). Second, inspection of the genomic context of representative eukaryotic and prokaryotic Tiamat proteins that clade together reveals that eukaryotic variants often contain introns and are found adjacent to intron-containing genes, whereas the prokaryotic variants contain no introns and are adjacent to intron-less genes that encode prokaryotic proteins (**Fig. S3**). These differences suggest that the closely related bacterial and eukaryotic sequences do not arise from misannotation or sequence contamination. In sum, most eukaryotic Tiamat sequences are distinct from prokaryotic ones, but a minority appear to bridge this transition (**Fig. 1, 2**).

### The uncharacterized core of Tiamat comprises repeated TPR-like folds, a novel fold and a metalloprotease-like fold

The heterogeneous region of the Tiamat core that bridges the ATPase and PD-(D/E)xK domains is annotated to varying degrees as “DUF3684” (3). To define this core region, we predicted structures for a representative sequence from each clade with AlphaFold3 (**Fig. 2b, S10 and S11**) (68). Manual inspection of these structures reveals that this region contains several conserved folds connected by flexible linkers, like beads-on-a-string. This interdomain flexibility causes whole-protein structural alignments to fail (**Fig. S10 and S11**). We therefore segmented the predictions with TED (<u>T</u>he <u>E</u>ncyclopedia of <u>D</u>omains), which uses an AI-based approach to detect domains and has classified 365 million domains in the AlphaFold Protein Structure Database (69). Tiamat sequences from clade 8 were excluded from this analysis because large and frequent low complexity insertions prevented confident structural prediction of most of the protein.

Structural segmentation and comparison to experimentally solved structures resolved the N– and C-terminal catalytic domains. First, we identified an extension of the previously annotated ATPase domain (**Fig. S6c**). A Foldseek search showed that these extensions resemble a region of Hsp90 homologs required for ATP binding and hydrolysis, termed the “transducer” (probability > 0.3). In Hsp90, this domain contributes a key arginine that contacts the γ-phosphate of ATP, and the domain communicates the structural changes that occur during ATP binding to the C-terminus (**Fig. 3a, S6 and S7**) (30, 83–87). The equivalent domain in *B. cereus* Tiamat contributes a similarly positioned arginine (**Fig. 3a, S6 and S7**). Second, the region previously annotated as a NOV_C or DUF3883 domain shares structural and active-site homology with experimentally verified PD-(D/E)xK-family nucleases (**Fig. 3b, S7 and S8**). The carboxylate residues of the PD-(D/E)xK motif typically coordinate a metal ion that acts as a cofactor, while the lysine residue positions a water molecule for an in-line attack on the scissile phosphodiester bond. Most studied members of this nuclease family cleave DNA although some variants cleave RNA (46). Together these assignments suggest that Tiamat possesses both ATPase and nuclease activity.

Both catalytic domains of Tiamat are predicted to homodimerize, as in other GHL proteins (32–36). Highly confident predictions have an interface predicted template modelling (ipTM) scores >0.8, whereas predictions that may be correct or incorrect had a score between 0.6-0.8 (medium confidence), and predictions that likely failed have a score <0.6 (low confidence) (88). The predictions of ATPase dimers for Tiamats from B. cereus, A. thaliana and nearly all clades were highly confident, whereas clade 8 was medium confidence (**Fig. S6**). The predictions of nuclease dimers for B. cereus, A. thaliana and seven clades were highly confident, clade 7 was medium confidence, and clade 4 was not confident (**Fig. S8**). In both domains, flanking residues assist in forming the dimerization interface (**Fig. 3a,b, S6 and S8**). The 20–42 residues N-terminal to the ATPase domain are predicted to behave differently across homologs. In some cases the N-terminal flanking residues form a pair of α-helices that wrap around each other, in others a single α-helix reaches from one subunit across the apex of the other, and in one homolog it is unclear whether these residues contribute to the dimer interface. The 12–36 residues C-terminal to the nuclease are predicted to form one to four additional β-strands that pair edge-to-edge with each other or with β-strands of the PD-(D/E)xK core, to form β-sheets. A disordered linker typically tethers the C-terminal nuclease domain to the rest of the protein. Because the nuclease domain and its C-terminal extension dimerize, they may take the role played by the dedicated dimerization domain of Hsp90. Most full-length Tiamat representatives are predicted to dimerize through both the ATPase and nuclease domains, and to possess a flexible intervening region that sometimes forms a lumen, though the central region is highly flexible and negatively impacts the ipTM score such that most are low-confidence and the lumen is occluded (**Fig. 2c, S6, S8, S11**). The full-length Tiamat representatives of clade 1 and clade 5 were predicted with medium confidence (**Fig. 2c**).

Between the ATPase and nuclease domains, we find copies of at least three distinct folds (**Fig. 2b and S9**). Representatives of the first identifiable fold C-terminal to the ATPase commonly matched TPR or Armadillo (ARM) folds in searches with the structure comparison tool Dali (**Fig. 2b and S9**) (82). Tiamat TPR-like folds typically occur in 1–6 copies and often sandwich a second fold, which we term “Fold B” because it could not be confidently matched to any experimental structure (**Fig. 2b and S9**). TPR is a helix-turn-helix fold that often occurs in tandem arrays of 3–16 motifs, which pack into a super-helical structure presenting paired concave and convex surfaces that facilitate binding to target proteins (89, 90). Tiamat proteins may therefore use these surfaces to recognize viral antigens, although this remains to be tested. The third fold, found across many bacterial and eukaryotic Tiamat proteins, matched metalloprotease domains, and five of seven representative examples of this third fold matched to the eukaryotic metalloprotease Wss1, a critical genome maintenance enzyme that degrades covalent DNA-protein crosslinks (91, 92). However, the metalloprotease-like folds in Tiamat lack the canonical HExxH active site motif, and the residues that would coordinate active-site or structural metal ions are either absent or too far apart to coordinate a metal together, making proteolytic activity unlikely (**Fig. S9**) (93, 94). So called “pseudoenzyme” proteins and domains, which lack amino acids required for catalysis, but which are structurally homologous to active enzymes, exhibit a range of functions that include substrate sequestration, allosteric regulation, trafficking and others (95, 96). For example, the inactive homologs of rhomboid serine proteases (iRhoms) bind similar transmembrane client proteins as their active counterparts, but do not cleave and mature them (97, 98). Instead iRhoms bind and traffic their client proteins to target them to the ER degradation pathway. It’s possible the metalloprotease-like folds of Tiamat have similarly evolved a new role in binding to, but not cleaving, client proteins. Tiamat proteins from clades 2 and 3 sometimes contain additional “Other” folds that could not be confidently matched to known structures. Together, these assignments resolve the previously uncharacterized DUF3684 annotation, which spans the C-terminal half of the ATPase domain, the TPR-like folds and, where present, Fold B, but not the metalloprotease-like fold.

### The plant developmental protein NO VEIN is a Tiamat homolog with conserved active sites

Among the eukaryotic Tiamat homologs identified here is NO VEIN (Nov) from *A. thaliana*, named for the absent leaf veins in mutant plants. Nov was originally described as a “plant-specific” factor that mediates auxin-dependent cellular differentiation and patterning of the embryo, shoot and root, and whose loss is embryonic lethal (50, 51). It is strongly expressed across tissues and life stages, and a GFP-tagged fusion localizes to the nucleus (50, 51, 99). Our sequence and structural analyses identify Nov as a eukaryotic Tiamat homolog (**Fig. 3**). Nov possesses an Hsp90-like ATPase domain, and the residues required to bind and hydrolyze ATP, including the arginine contributed by the ATPase extension, are shared with those of *B. cereus* Tiamat and Hsp90 (**Fig. 3a**). Nov’s predicted nuclease domain likewise matches experimentally studied DNases, with the key active-site residues conserved (**Fig. 3b**).

*A. thaliana* Nov and *B. cereus* Tiamat also share structural similarity between the ATPase and nuclease domains. In both proteins a TPR-like fold flanks Fold B on either side, and in Nov this pair is flanked in turn by an additional pair of TPR-like folds. Additionally, both proteins share a metalloprotease-like fold that caps the most C-terminal TPR-like fold (**Fig. 3c**). However, these two proteins differ at their termini. *B. cereus* Tiamat carries two additional “Other” folds between the metalloprotease-like fold and the nuclease, a feature found in clade 2 sequences, while Nov carries a long N-terminal extension, as do most eukaryotic Tiamat homologs (**Fig. 2b, 3c**). Nov therefore shares not only both predicted active sites, but also the intervening architecture between these sites with the antiviral *B. cereus* Tiamat (**Fig. 3c**).

### *B. cereus* Tiamat is an active ATPase that cleaves ssDNA and nicks dsDNA

Given the conservation of ATPase and nuclease active-site residues across Tiamat homologs, we next asked whether the prototypical Tiamat from B. cereus, which was previously shown to provide phage defense, exhibits these activities in vitro. We over-expressed His_6_-tagged B. cereus Tiamat and purified it to homogeneity by nickel-affinity, heparin and size-exclusion chromatography (**Fig. 3d**). To monitor ATP hydrolysis we used a coupled colorimetric assay in which released inorganic phosphate drives the generation of 2-amino-6-mercapto-7-methylpurine, which absorbs strongly at 360 nm. Incubation of 100 nM Tiamat with 1 mM ATP produced a steady increase in absorbance over 1 hour, whereas ATP or Tiamat alone did not, indicating that B. cereus Tiamat is an active ATPase (**Fig. 3e**). The slightly higher baseline of ATP alone relative to Tiamat alone is consistent with a small amount of spontaneous hydrolysis in the ATP stock.

We next tested whether the putative C-terminal PD-(D/E)xK nuclease domain is active, and whether it preferentially cleaves 54-nt long ssDNA, ssRNA or dsDNA substrates (**Fig. 3f**). A five-fold excess of Tiamat dimer (0.5 µM) was incubated with each nucleic acid substrate (100 nM) for 1 hour at 37°C. Tiamat cleaved ssDNA efficiently, converting most of the substrate into approximately six discrete products, and did not detectably cleave ssRNA (**Fig. 3f and S12**). The ssDNA cleavage products corresponding to the most 5’– and most 3’-proximal cleavage positions are the most abundant, and those from intermediate positions are less so. Tiamat also cleaved the Cy5-labeled strand within the dsDNA substrate, though far less efficiently, generating three discrete products, of which the product corresponding to cleavage nearest the 3’ end is the most abundant (**Fig. 3f and S12**). These three dsDNA products appear to coincide with cleavage positions in the ssDNA substrate, suggesting Tiamat has a sequence preference. Collectively these data show that *B. cereus* Tiamat is an ATPase and a DNase that prefers ssDNA under the conditions tested, and that it can nick at least one strand of a duplex at defined positions. Given the conservation of active-site residues in *A. thaliana* Nov (**Fig. 3a,b**), Nov may share these activities.

### Eukaryotic Tiamat proteins carry N-terminal LOTUS domains and are predicted to localize to processing bodies

The eukaryotic and prokaryotic Tiamat proteins identified differ greatly at their N-termini. Prokaryotic Tiamat proteins, and the eukaryotic sequences that clade with them, have short N-terminal extensions preceding the ATPase domain (1–124 residues, median of 30 residues), whereas most other eukaryotic sequences have far longer extensions (3–2,047 residues, median of 1,139 residues) (**Fig. S4**). C-terminal extensions were less variable and typically short, both in eukaryotic clades (1–902 residues, median of 29 residues), and in intermixed and prokaryotic clades (2–192 residues, median of 38 residues) (**Fig. S4**).

To determine whether Tiamat proteins possess fusions that might impart additional biochemical activities, we annotated the UniProt– and RefSeq-derived sequences with InterProScan and HHPred, then manually curated the results (**Fig. 4, Data S3-S6**) (45, 59). Additional domains are rarely fused to prokaryotic Tiamat proteins (**Fig. 4a**). Two bacterial homologs possess an additional C-terminal nuclease domain similar to type IV restriction enzymes, which cleave methylated DNA (100). Single archaeal sequences carry a tRNA endonuclease-like domain or a type II restriction endonuclease domain (**Data S3, S5**). Fusions are common in eukaryotic Tiamat proteins, by contrast. N-terminal LOTUS domains occur in 316 of the 918 eukaryotic sequences (34.4%) (**Fig. 4a**). LOTUS domains occur frequently in proteins involved in RNA metabolism, where they bind RNA, or recruit RNA-interacting proteins such as the RNA helicase Vasa (101–104). A few LOTUS-containing Tiamat proteins also carry an N-terminal sulfotransferase domain, which modifies substrates ranging from steroids to proteins to alter their solubility or stability (105). A further 16 eukaryotic Tiamat proteins (1.7%) possess one of two N-terminal zinc finger domains, which are versatile nucleic acid-, protein– and ligand-binding folds (**Fig. 4a**) (106). The prevalence of LOTUS domains suggests that a subset of eukaryotic Tiamat proteins acts on or with RNA.

N– and C-terminal extensions often harbor signals that direct proteins to specific subcellular compartments. To ask whether eukaryotic Tiamat proteins carry such signals, we analyzed all 918 eukaryotic sequences with ProtGPS at a stringent probability threshold of 0.8 (**Fig. 4c, S13 and Data S7**) (73). ProtGPS is a protein language model trained on 5,480 human proteins annotated as localizing to 12 subcellular condensate compartments (73).

Therefore, the following predictions for non-human extrapolate past ProtGPS’s training data. More than 20% of eukaryotic Tiamat sequences (191 proteins) are predicted to localize to processing bodies (P-bodies), which are cytoplasmic ribonucleoprotein granules composed largely of translationally repressed mRNAs and proteins involved in mRNA decay (107). Because LOTUS domains also imply RNA-related functions, we asked whether these were the same proteins. Roughly half of the P-body-localizing proteins contain a LOTUS domain (**Fig. 4d**). Consistent with the N-terminal position of these domains, Tiamat proteins truncated at the N-terminus frequently lose their predicted P-body localization and are sometimes assigned to other compartments, whereas C-terminal truncations leave most predictions unchanged (**Fig. S14a**). Together these annotations suggest that the predicted localization of a subset of eukaryotic Tiamat proteins to an RNA-rich compartment derives from their N-terminal extensions, which often contain LOTUS domains.

The next most common predicted localizations are to the post-synaptic density and to chromatin, for 11 and 8 proteins respectively, with single proteins assigned to the nucleolus, the nuclear pore complex or the cell junction (**Fig. 4c and S14a**). Predicted P-body localization occurs across clades 5–10 (6–26% of sequences) but is absent from clade 4. The 13 athropod, placozoan and rotifer sequences in clade 4 instead contribute 3 of the 8 proteins predicted to localize to chromatin, the remaining 5 coming from clades 7 and 8 (**Fig. S14b**). Unexpectedly, all 11 proteins predicted to localize to the post-synaptic density (PSD) belong to clade 8 (15% of that clade) and originate from organisms that lack synapses, such as algae and dinoflagellates (**Table S1**). Some of these are likely false positives of the kind expected in large datasets, but ProtGPS may also be detecting features shared with membrane-associated condensates more generally. Consistent with the latter possibility, 4 of the 11 PSD-localizing proteins carry additional domains, and 3 of those domain types are enriched among the more than 700 proteins found at the PSD (**Fig. S14c**) (108–110).

We next asked whether the LOTUS fusions identified above occur sporadically across eukaryotic Tiamat proteins or are concentrated in particular taxonomic groups. In the UniProt– and RefSeq-derived dataset, LOTUS fusions appear restricted to aquatic organisms, though these span a wide phylogenetic range that includes green algae of the class Chlorophyceae, sponges, cnidarians, molluscs, echinoderms and both jawed and jawless fishes (**Fig. 4b**). Intriguingly, despite the abundance of Tiamat homologs found in green plants, none have LOTUS fusions. Most LOTUS domain-containing Tiamat proteins, 257 out of 316 proteins, come from ray-finned fish (class Actinopteri), the same lineage that supplies most of the aquatic chordates encoding a Tiamat homolog (**Fig. 4b**). These proteins can possess several LOTUS domains, and the Tiamat from the model organism zebrafish has four (**Fig. 4a**). Little is known about the function of these LOTUS-containing Tiamat proteins, though their genes are highly expressed in many tissues. For example, zebrafish Tiamat (Bgee gene identifier: ENSDARG00000096562) and several close fish homologs share high expression in the testes, digestive system, hepatobiliary system, brain and musculoskeletal system (111).

### Tiamat-like proteins swap the PD-(D/E)xK nuclease for alternative effectors

Antiviral defense systems often evolve by shuffling sensor and effector domains into new combinations (112). Because our requirement for both an Hsp90-like ATPase and a PD-(D/E)xK domain would have excluded “Tiamat-like” proteins that had undergone such shuffling, we re-examined the sequences this curation step removed, retaining those sequences with HMM scores comparable to curated Tiamat proteins (**Fig. S15**). Across kingdoms the PD-(D/E)xK nuclease of Tiamat-like proteins is frequently replaced by a wide range of alternative effector and recognition domains, appended to either the N– or C-terminus.

Across bacteria, archaea and eukaryotes, some Tiamat-like proteins have divergent PD-(D/E)xK domains that InterProScan missed but that HHPred identifies (Type I-A) (45, 59). Others lack the PD-(D/E)xK domain and consist of just an ATPase and DUF3684 domain (Type II-A) (**Fig. 5**). Unique to bacteria, the PD-(D/E)xK domain is often replaced by one of two helicases, a UvrB-like (Superfamily 2; SF2) helicase that may act on DNA, and a Upf1-like (Superfamily 1; SF1) helicase that may unwind RNA (Types III-A and III-B respectively) (**Fig. 5a**). The prototypical UvrB helicase is a translocase that localizes to damaged DNA and forms a stable DNA-protein complex that recruits the UvrC nuclease to excise the damage (113). While Upf1, the key effector for eukaryotic nonsense-mediated decay of mRNAs, can translocate on ssRNA and DNA, and unwind both RNA and DNA (114). Helicases play an important role in eukaryotic and prokaryotic antiviral immunity, often functioning as sensors or effectors (115–117).

In both bacteria and archaea, the PD-(D/E)xK domain is sometimes replaced by an HNH-like fold that typically lacks the catalytic residues required to cleave DNA (Type IV-A) (**Fig. 5a,b**) (118, 119). The absence of these catalytic residues does not preclude activity, since the HNH nuclease PacI cleaves DNA without them (118). These HNH-like folds may therefore be catalytically active, or may have diverged to a non-catalytic role in DNA binding. In another common bacterial Tiamat-like protein, the PD-(D/E)xK domain is replaced by two domains, a Schlafen-like RNase domain that lacks known catalytic residues (120, 121), and a peptidase that retains active-site residues (Type V-A) (**Fig. 5a**). Schlafen proteins, first described as regulators of mouse thymocyte development, restrict viruses by cleaving tRNAs and rRNAs, and bacterial Schlafen proteins cleave tRNAs to restrict phage infection (120, 122). Antiviral defense systems in both eukaryotes and prokaryotes frequently use peptidases to mature proproteins or proenzymes into their active forms once a viral antigen is recognized, which elicits downstream antiviral responses (123–126).

Both bacteria and eukaryotes possess Tiamat-like proteins that lack the PD-(D/E)xK domain and instead carry N-terminal fusions to domains involved in RNA metabolism. Bacterial Type VI-A proteins possess S1 domains typically found in ribosomal protein S1, which likely bind RNA directly (127). Eukaryotic Type VIII-A proteins contain LOTUS domains, which are reported to bind to G-rich sequences or RNA-acting enzymes like the Vasa helicase (102, 104) (**Fig. 5a,c**).

In eukaryotes such as filamentous fungi, Tiamat-like proteins with or without the PD– (D/E)xK domain carry a complete het-6-like protein fused to the C-terminus (Type VII-A and VII-B respectively) (**Fig. 5c**). Fungal filaments fuse to form a mycelial network that supports their growth, and Het-6 is one of several proteins that detect whether such a fusion has merged cells of two different strains. Het-6 senses which variant of the neighboring Un-24 protein is present in the cell, and triggers cell death if it detects a non-self variant (128–130). In discriminating one fungal strain from another, Het-6 confronts a self versus non-self recognition problem analogous to that of distinguishing host from virus.

Several eukaryotic Tiamat-like proteins are truncated at the N-terminus, lacking the ATPase domain but retaining DUF3684 and PD-(D/E)xK (Type IX-A). Some plant Tiamat-like proteins lack the C-terminal PD-(D/E)xK domain and instead carry a complete cyclin-dependent kinase (CDK) fused to the N-terminus (Type X-A). CDKs are serine/threonine protein kinases that regulate the cell cycle and are periodically activated by cyclins (131). The final group of eukaryotic Tiamat-like proteins identified here appears to comprise aquatic chordate homologs of the human protein SACSIN. Mutations in the human *SACSIN* gene cause a neurodegenerative disorder associated with impaired mitochondrial quality control (52, 132, 133). These proteins share SACSIN’s characteristic three ATPase domains followed by a C-terminal J-domain and HEPN domain, but lack the N-terminal ubiquitin-like and protein-binding XPCB domains of the human protein (Type XI-A) (**Fig. 5c**) (53). HEPN (Higher Eukaryotes and Prokaryotes Nucleotide-binding) RNases contribute to antiviral defense in both eukaryotes and prokaryotes, as well as to mRNA splicing and ribosomal RNA maturation, and must dimerize to form a composite catalytic center (134).

### Tiamat and Tiamat-like proteins are closely related to SACSIN

The identification of SACSIN proteins as putative Tiamat-like proteins (Type XI-A) (**Fig. 5c**) led us to probe the relative phylogenetic relationships between Tiamat homologs, SACSIN, and other GHL ATPases involved in defense and non-defense roles (**Fig. 5d**). The ATPase, N-terminal flank, and C-terminally adjacent transducer-like region in all selected GHL proteins were extracted and aligned to determine their relationship (**Data S1**). The first ATPase domain of the multiple repetitions found in SACSIN homologs was used for this analysis. The resulting phylogenetic analysis recapitulated previously established relationships. For example, MutL, gyrase (Type IIA topoisomerase), and Hsp90 enzymes form distinct clades, further, Type IIA topoisomerases form a clade distinct from Type IIB topoisomerases (18, 65, 135).

Tiamat, Tiamat-like and SACSIN proteins form a large diverse clade that is confidently separated from other classical GHL ATPase family members (Gyrase, Hsp90, MutL) and from other GHL ATPases with proposed or demonstrated antiphage and antiviral roles (prokaryotic MORC, paraMORC, Hsp90-like antiphage proteins, plant MORC) (18, 20, 21, 34, 35). Hsp90 and putative antiphage Hsp90-like proteins form an intermixed clade that is the next closest group of sequences related to the “Tiamat – Tiamat-like – SACSIN” clade. Prokaryotic MORC and paraMORC sequences appear to form a clade that is distinct from MORC and MutL, though the support values for these branches are lower. Collectively, proteins across the GHL ATPase family have been co-opted for defense, and of the families surveyed here, Tiamat is most closely related to SACSIN and Hsp90.

### Tiamat and Tiamat-like proteins are frequently encoded near nucleic acid-modifying enzymes

Many antiphage defense systems comprise multiple genes encoded next to one another, whose products form a complex or otherwise cooperate to sense and defend against phage infection. We therefore asked whether Tiamat co-occurs with particular defense genes by analyzing the ten genes flanking each prokaryotic Tiamat homolog with the Enzyme Function Initiative (EFI) webserver (**Fig. 6a**–**c**) (74).

Conserved proteins with putative defense roles were found within five genes of Tiamat (**Fig. 6a, b**). The most common conserved neighbor is a helicase, encoded beside roughly 14% of curated prokaryotic Tiamat proteins and roughly 33% of Tiamat-like proteins, found in all but one neighborhoods identified (Type I-G, I-H, I-I, I-J, I-K, II-B, II-C, III-C, IV-B, VI-A) (**Fig. 6a, b, c**). Associated helicases belong to either SF1 (UvrD, Upf1, RecB) or SF2 (Hrq1, Swi/Snf, T1R, UvrB) (136–139). The associated helicase is often fused to a nuclease domain (I-H, I-I, I-J, II-B, II-C, IV-B, VI-A). A DNA methyltransferase gene, predominantly targeting adenine and sometimes targeting cytosine, co-occurs only where a helicase is also present (Types I-H, I-J, I-K, II-C and III-C). In some neighborhoods a putative transcriptional regulator is also conserved, resembling those that regulate the MADS or BREX defense systems (MADS1-like HTH and BrxR-like WYL), both of which are large multi-gene systems that rely on encoded methyltransferases to distinguish self from foreign DNA (140–145) (**Fig. 6a, b**).

The shuffling of Tiamat with components of other defense systems is more nuanced than putative transcriptional regulators. The Type I-J neighborhood appears to be a type I R-M system with an embedded Tiamat gene (**Fig. 6a**). Type I R-M systems comprise a nuclease, a methylase and a specificity subunit that assemble into a large heterooligomeric complex, which methylates hemi-methylated DNA and cleaves unmethylated DNA (146). In the Type III-C neighborhood, a Tiamat-like protein carrying a C-terminal helicase in place of the canonical PD-(D/E)xK nuclease is associated with a cytosine methyltransferase and with three proteins drawn from other defense systems. These three proteins are GajB of Gabija, which hydrolyzes and depletes (d)A/(d)GTP molecules in the cell upon sensing DNA termini, and DrmA and DrmB of DISARM, a helicase complex proposed to recognize unmodified DNA and recruit DrmC or nucleases to degrade foreign DNA (2, 147–150).

The neighbors of Tiamat and Tiamat-like proteins also include helicase-nuclease fusions that may act on RNA. One helicase of the Type I-K system is fused to an N-terminal RelE-like nuclease domain (**Fig. 6a**). RelE toxins are ribosome-dependent endoribonucleases often involved in stress response, and a CRISPR-associated RelE fusion protein inhibits translation to provide antiphage defense (151, 152). In the Type II-B and VI-A systems the Tiamat-like proteins both lack the PD-(D/E)xK nuclease, and those of Type VI-A instead carry N-terminal S1 domains that likely bind RNA (**Fig. 6b**) (127). Both neighborhoods contain a protein that combines a Upf1-like RNA helicase domain often involved in eukaryotic nonsense-mediated decay, and a PIN nuclease domain that often cleaves RNA in the context of nonsense-mediated decay (153–155). In prokaryotes the PIN (PilT N-terminus) domain also frequently serves as the toxin of toxin-antitoxin systems, and a CRISPR-associated PIN fusion cleaves RNA in response to phage detection (154, 155).

Not all Tiamat-like proteins captured by the loosened curation criteria have a gene neighborhood consistent with a role in defense. Nearly 14% of Tiamat-like proteins, predominantly from Streptomyces, do not co-occur with known defense genes and are instead frequently encoded next to vitamin synthesis and membrane transporter genes (**Fig. 6c and S15**).

### Type I-G Tiamat proteins are predicted to form a complex with an associated NERD-helicase

Antiviral proteins encoded near each other and cooperating in defense often form complexes. Given this propensity, and the ssDNase activity of B. cereus Tiamat (**Fig. 3f**), we asked whether Tiamat and an associated helicase would be predicted to interact. We focused on the Tiamat and helicase in Type I-G systems because of their prevalence and minimal complexity. We modelled a fragment of Tiamat spanning the nuclease to the C-terminus together with the NERD-helicase protein, in the presence of 1 ATP for the helicase active site, a 40-nt ssDNA, and 5 Mg^2+^ ions. Tiamat C-terminus and NERD-helicase were modelled in a 1:1 stoichiometry because SF1 helicases (such as UvrD) act as monomers (138). AlphaFold3 returned consistent predictions for all 12 non-redundant homologs tested, with medium to high interface confidence values (ipTM 0.67–0.86, pTM 0.73–0.89) (**Fig. 6d and S16**).

NERDs (Nuclease-related domains) are enriched in prokaryotic defense islands, and co-occur with certain prokaryotic viperin proteins and with some reverse transcriptases that may both play a role in antiphage defense (11, 42, 156). Type I-G Tiamat proteins had longer extensions C-terminal to the PD-(D/E)xK nuclease domain (83-100 residues) than the clade representative Tiamat proteins (12-40 residues) (**Fig. 2**). Structure prediction reveals that the C-terminal extension of Type I-G Tiamat proteins folds into an SH3-type β-barrel (**Fig. 6d and S16**). This β-barrel is predicted to recruit the NERD-helicase protein by binding one specific face of the helicase domain (**Fig. 6d**). The predictions suggest that the NERD domain binds ssDNA and threads it through the helicase domain to feed ssDNA to the Tiamat nuclease, which efficiently cleaves ssDNA (**Fig. 3f, 6d**). Analogously, the bacterial GHKL ATPase involved in mismatch repair, MutL, recruits and loads the DNA helicase UvrD onto a nicked DNA site to initiate degradation of the mismatched strand (157).

Collectively, Tiamat proteins and Tiamat-like proteins are most commonly associated with helicases, which may generate the ssDNA that B. cereus Tiamat prefers (**Fig. 3f, 6e**). Tiamat proteins also frequently co-occur with DNA methyltransferases, which may be used to distinguish self from non-self, and with additional nucleases. The extra nucleases may enable the system to circumvent anti-defense genes that target a single nuclease, or may provide nuclease activity to a Tiamat-like partner protein that lacks the PD-(D/E)xK active site.

## DISCUSSION

Here we demonstrate that the antiphage protein Tiamat from *B. cereus* is an ATPase and ssDNase, with homologs found across the tree of life (**Fig. 1, 3**). Tiamat proteins share an N-terminal GHKL ATPase, a middle region (itself consisting of TPR-like folds, an unidentified fold we term “Fold B” and a metalloprotease-like fold that appears proteolytically inactive), and a C-terminal PD-(D/E)xK nuclease domain (**Fig. 2, 3**). This architecture resolves the previously uncharacterized DUF3684 annotation, which spans the C-terminal half of the ATPase, TPR-like folds and, where present, Fold B (**Fig. 2**). Both the middle region and the C-terminal effector vary among homologs (**Fig. 2, 5 and 6**). The *A. thaliana* developmental regulator Nov retains similar active-site residues in both catalytic domains and may therefore share biochemical activities with *B. cereus* Tiamat. Our findings raise the question of whether eukaryotic Tiamat homologs have been exapted solely for non-defense functions or retain their antiviral role.

Tiamat’s N-terminal Hsp90-like ATPase domain assigns it to the GHL protein family, whose roles in antiphage defense have emerged only recently, and Tiamat’s overall architecture mirrors that of well-characterized members (3, 18, 20, 21). This architecture includes an N-terminal GHKL ATPase domain, a set of middle domains of undetermined function, and a C-terminal nuclease domain predicted to dimerize (**Fig. 2, 3 and S17**). By analogy to the constitutive C-terminal dimerization of other GHL proteins, we propose that the C-terminal Tiamat dimerization interface is maintained throughout the ATPase cycle while the N-terminal domains cycle between dissociated (apo or ADP-bound) and associated (ATP-bound) states (**Fig. 6e**) (32–36). ATPase activity is required for defense, as the active-site substitution E44A abolishes the protection that Tiamat confers (3). Equivalent substitutions in chicken Hsp90 (E46A) and *E. coli* gyrase B (E42A) permit ATP binding but prevent hydrolysis, trapping those enzymes in an ATP-bound, closed state (158, 159). The failure of Tiamat^E44A^ to protect therefore suggests that ATP binding alone is not sufficient, and both the ATP turnover and cycling between closed and open states are required (**Fig. 6e**).

How substrate may be held during the predicted open-to-closed cycle is unclear. In other GHL proteins, an S5 domain 2-like fold (S5-fold) within the middle region of the protein engages diverse substrates (**Fig. S17**). In Hsp90, the S5-fold binds client polypeptides and co-chaperones, and in the topoisomerase IV subunit ParE this fold binds a DNA duplex that must pass through a transient double-stranded break (**Fig. S17**) (25, 29, 32, 33). In place of the S5-fold, Tiamat may use its TPR-like folds, Fold B or metalloprotease-like fold to capture viral proteins or nucleic acids within the lumen of the dimer. Tiamat’s middle region exhibits variability primarily in the number of TPR-like folds, but also in the presence of Fold B and the metalloprotease-like fold (**Fig. 2**). The low sequence conservation and frequent rearrangement of the middle region lead us to hypothesize that it is a rapidly evolving sensor, adapting to recognize new phage-encoded ligands, analogously to the middle regions of other GHL proteins that evolve to bind chemically diverse substrates (**Fig. 6e, S10 and S17**). However, the low predicted structural confidence of this region in most homologs prevent us from identifying a candidate binding surface (**Fig. 2, 3 and S10, S11**). The substrate-bound state of dimeric human MORC2 remains structurally uncertain as well, as MORC2’s lumen is thought to trap dsDNA through positively charged residues within a disordered linker, an arrangement our predictions do not recapitulate (**Fig. S17**) (22, 23).

Whatever substrate is captured by Tiamat, closure of the N-terminal ATPase domains around it may strain the dimerization interfaces and modulate ATPase or nuclease activity, as previously seen in Hsp90 and proposed for type II topoisomerases (160, 161). Such long-range coupling is also proposed in a different family of prokaryotic Hsp90-like defense proteins that carry N-terminal HD or calcineurin-like phosphoesterase domains and rapidly evolving, modular C-terminal domains. The C-terminal domains of these Hsp90-like proteins are thought to bind foreign or endogenous molecules which signal an active infection, thereby triggering a conformational change that activates the fused nucleotide-hydrolyzing effector (20, 32).

Tiamat’s genomic context suggests that it does not always act alone. Tiamat and Tiamat-like genes are frequently encoded adjacent to genes coding for DNA helicases, methyltransferases, and additional nucleases (**Fig. 6a, b**). Helicases and NTPases are frequently found in prokaryotic and eukaryotic antiviral defense systems, where they act as sensors of phage infection, unwind viral nucleic acid, or both (2, 3, 75, 115–117, 146, 162, 163). For example, the Hachiman system, composed of the PD-(D/E)xK nuclease HamA and the SF2 helicase HamB cooperate to sense, unwind and degrade viral dsDNA (116, 117, 164). While, in the case of PARIS, the AriA ATPase subunit sequesters the AriB nuclease subunits in an inactive state, in a complex (AriA6-AriB3). Binding of phage “trigger” proteins stimulates AriA ATPase activity and the release of AriB to cleave host tRNAs, stalling translation in the infected cell (165). Methylation may mark self-DNA and inhibit DNA sensing or cleavage, analogously to traditional restriction-modification systems and has been hypothesized or in some cases demonstrated for other recently described multigene defense systems such as BREX, MADS, DISARM and Hma (48, 140, 144, 146, 149).

Prokaryotic MORCs and paraMORCs, two other types of GHL proteins, are also frequently neighbored by helicases, nucleases and methyltransferases (**Fig. 6a,b**) (21). Based on these genomic associations, prokaryotic MORCs and paraMORCs have been proposed to mediate a restriction-modification-like defense in which they sense unmethylated or hemimethylated DNA and recruit nucleases and helicases to nick, reel, unwind and then degrade foreign nucleic acid (21). Given its domain architecture and comparable genomic associations, we speculate that Tiamat acts analogously (**Fig. 6e**). The resemblance may be convergent since the methyltransferases encoded near Tiamat predominantly modify adenine, whereas those near prokaryotic MORCs and paraMORCs are predominantly predicted to target cytosine (**Fig. 6a, b**) (21).

Since Tiamat nicks dsDNA inefficiently but preferentially degrades ssDNA, we speculate and propose a model that Tiamat introduces an initial nick, a partner helicase then loads onto the nick and unwinds the duplex, and the exposed ssDNA is reeled in and degraded (**Fig. 3f and 6e**). Several Tiamat proteins (14%) are encoded next to a putative helicase or translocase and may directly interact with these for defense, but the majority are not encoded adjacent to a conserved helicase and may instead act with other endogenous helicases in the cell, or even act alone (**Fig. 6a**). Tiamat’s ATPase cycle may regulate this process, with closure capturing substrate within the lumen and hydrolysis driving opening and product dissociation, so that the enzyme can act on successive nucleic acid substrates. Alternatively, substrate binding within the Tiamat lumen may trigger a conformational change that modulates ssDNase and dsDNase activities. The additional nucleases and multiple helicases found in some Tiamat systems may broaden the range of substrates or motifs that can be cleaved, or provide redundancy against the anti-defense proteins phages frequently deploy to evade immunity (166). A caveat is that our model remains untested, since we have not reconstituted a Tiamat-helicase complex, the interaction is supported only by structure prediction (**Fig. 6d, e**), and the dsDNA nicking on which the model depends is inefficient under the *in vitro* conditions used (**Fig. 3f**). Establishing the model will require showing that Tiamat and a helicase act on the same substrate in tandem, and may require the identification of a substrate or ligand that binds the Tiamat lumen to activate the complex.

Our model of Tiamat function can be extended to many homologs but will require modifications for several Tiamat-like proteins. Antiviral defense systems often evolve by modular shuffling of sensor and effector domains to generate new protein fusions or new protein complexes, and Tiamat is no exception (**Fig. 6**) (112, 167, 168). Examples of such extensive shuffling of sensor and effector modules have been described for CRISPR, BREX and Menshen defense systems, to name a few (43, 163, 169–171). Among these systems, sensors include NTPases, β-sandwiches, inactive nucleases, winged helix-turn-helix motifs, CARF and SAVED domains, whereas effectors can range from DNases and RNases to proteases, proteins that form membranes pores, ATP deaminases, adenylate cyclases and nitrilases. Tiamat’s C-terminal effector is interchangeable. Relaxing our curation thresholds identifies many Tiamat-like proteins in which the nuclease is truncated or replaced by helicases that might unwind foreign nucleic acid, peptidases that may trigger a proteolytic cascade, nuclease-like folds that could bind or cleave DNA, or, in Het-6, a combined sensor and effector of cell death (**Fig. 5**) (172). Strikingly, Tiamat-like proteins that lack a C-terminal nuclease and have no annotated replacement are frequently encoded beside a putative RNA helicase-nuclease, suggesting that the effector may not need to be fused to the putative sensor domain but can be supplied in trans by a neighboring gene (**Fig. 6b**).

Modular addition also occurs N-terminal to the ATPase domain, most frequently in eukaryotes. Many Tiamat homologs from aquatic eukaryotes have long N-terminal extensions containing LOTUS domains and are predicted to localize to P-bodies, a prediction dependent on the N-terminal sequence (**Fig. 4, S13 and S14**). Together, these annotations suggest that a subset of eukaryotic Tiamat proteins may sense or act on RNA through their N-termini while retaining a C-terminal DNase. The phylogenetic distribution of LOTUS-Tiamat fusions is itself striking. LOTUS-Tiamat fusions are confined to aquatic organisms, spanning green algae to fishes, and are absent from land plants despite the abundance of Tiamat homologs in that taxonomic group (**Fig. 4b, S2**). Among those land plant homologs is the best-studied eukaryotic Tiamat, *A. thaliana* Nov, which is a nuclear-localized regulator of auxin-dependent patterning whose loss is embryonic lethal, and which nonetheless retains the active-site residues of both the ATPase and the PD-(D/E)xK nuclease (**Fig. 3a, b**) (50, 51). It is possible that Nov’s activities have been exapted for a purely developmental role, or Nov may simultaneously have both defense and non-defense roles, similar to Argonaute (12, 13).

The eukaryotic clades also include one whose members might be mistaken for pseudogenes. Clade 8 has sequences from Chlorophyceae, Dinophyceae and Conoidasida (green algae, dinoflagellates, and apicomplexans like the parasite Toxoplasma gondii), which contain many low-complexity insertions (**Fig. 2, S6, S8**). These insertions consist of poly-Ala, –Gly, –Gln, –Pro, and –Ser tracts predicted to form disordered loops between and within domains, including the ATPase and nuclease domains. Such low-complexity tracts are common in this part of the eukaryotic tree. An estimated 49–90% of the proteome of *Plasmodium falciparum*, a relative of *T. gondii*, consists of low-complexity regions (173). These insertions are therefore unlikely to be inactivating on their own, since most of the *P. falciparum* proteome would otherwise be non-functional. Although these insertions reduce overall prediction confidence enough that we excluded clade 8 from structural segmentation, the ATPase and nuclease active sites themselves appear unperturbed in the predicted structures (**Fig. 3, S6 and S8**). Therefore, we interpret this clade as reflecting lineage-specific low-complexity region expansion rather than inactivation of Tiamat.

Eukaryotes acquire genes by horizontal transfer from other eukaryotes and from prokaryotes, and these events contribute to important evolutionary adaptations (7, 174–177). MORCs are thought to have originated in bacteria and to have been acquired by eukaryotes in a single such event (21). Tiamat’s history appears less tidy. Its homologs are widespread across eukaryotes and enriched in plants and fish, and their respective eukaryotic parasites, including a clade dominated by Oomycota plant pathogens. Subsets from Placozoa, Arthropoda, Rotifera and Fungi, however, fall into several separate groups that clade with bacterial rather than eukaryotic sequences (**Fig. 1, 2, Table S1 and Data S2**). Three observations argue that these bacterial-like sequences are genuine eukaryotic genes rather than contamination, misannotation, or assembly artifacts. First, they contain introns or are encoded next to intron-containing genes and are flanked by eukaryotic genes, whereas the closest bacterial homologs lack introns and are flanked by bacterial genes. Second, they have close homologs in related eukaryotic species and genera, and third, they derive from large scaffolds, contigs or complete genomes rather than short fragments (**Fig. S3**).

The nearest bacterial Tiamat relatives differ according to the eukaryotic variant. The Trichonephila Tiamat is most like that of the endosymbiont Candidatus Rickettsiella viridis, the *Trichoplax* adhaerens protein to Clostridium and Paenibacillus spp. homologs, and the rotifer and anaerobic-fungal proteins to Bacteroidota homologs (Ginsengibacter and Mangrovibacterium spp. respectively) (**Fig. S3, Data S2**). Because a single transfer into a eukaryotic ancestor would place these sequences together rather than beside different bacterial lineages, we propose that Tiamat has been exchanged between prokaryotic and eukaryotic genomes on several independent occasions. The *Trichoplax* genes make one such acquisition look recent. The closest Trichoplax Tiamat relatives outside the genus are bacterial rather than eukaryotic, they are structurally more similar to bacterial Tiamats than to most metazoan and plant homologs, and they carry an intron density of 0.2 introns/kb (one intron in a 4.3–4.5 kb gene), far below the Trichoplax genome average of 7.6 introns/kb (**Fig. 2, S3 and Data S2**) (178). We therefore speculate that these prokaryotic-like Tiamats were acquired from bacterial prey by bacterivorous predators such as Placozoa and Rotifera, or from bacterial endosymbionts by their arthropod hosts, a route for which the *Rickettsiella* homolog identifies a plausible donor.

Here we define the previously undescribed biochemistry, architecture, modularity, and genomic contexts of Tiamat and its homologs. The functions of eukaryotic Tiamat and Tiamat-like homologs, like certain SACSIN proteins, nevertheless remain unclear. Eukaryotic homologs may have retained an antiviral role, been exapted for other biological purposes, or both, as in plant MORCs and Argonautes. We have not tested any eukaryotic homolog for antiviral activity, therefore our findings do not distinguish among these possibilities. Given that close Tiamat homologs are present in staple crops, commercially important fish, eukaryotic pathogens and parasites, and in the model organisms A. thaliana and D. rerio, these proteins merit further investigation (**Fig. 1, S2 and Data S2**). A key step toward resolving their function will be to identify what bacterial and eukaryotic Tiamat proteins capture within their lumens.

## ACKNOWLEDGEMENTS

We thank Alaíde Cordova for the illustrations of organisms used in Figure 1. The authors gratefully acknowledge the Penn Advanced Research Computing Center (PARCC) core facility (parcc.upenn.edu) for providing the computational infrastructure used in this study. PARCC is supported in part by the Office of the Vice Provost for Research at the University of Pennsylvania. We also thank members of the A.S-.F. laboratory for feedback and discussions.

## AUTHOR CONTRIBUTIONS

Shirley Yuan, Yuri Rafael de Oliveira Silva: Data Curation, Formal Analysis, Investigation, Methodology, Validation, Visualization, Software, Writing – original draft. Vinh Dang, Arav A. Nangia: Data Curation, Investigation, Methodology. Balwina Koopal: protein purification and *in vitro* assays. Jack Bravo: supervision, writing – review & editing. Andrew Santiago-Frangos: Conceptualization, Formal Analysis, Funding acquisition, Investigation, Methodology, Project administration, Resources, Supervision, Visualization, Software, Writing – original draft, review & editing.

## SUPPLEMENTARY DATA

### Supplementary Data statement

Supplementary Data, including supplementary figures, FASTA files of GHL ATPases, FASTA files all Tiamat and Tiamat-like sequences analyzed, InterProScan annotations of Tiamat proteins, ProtGPS annotations of Tiamat proteins, and a spreadsheet of Tiamat protein IDs along with their taxonomic and clade data are available at NAR online.

## DATA AVAILABILITY

Data has been made available as supplementary files published alongside the article.

## REFERENCES

1. Nasir, A., Kim, K.M. and Caetano-Anollés, G. (2012) Viral evolution: Primordial cellular origins and late adaptation to parasitism. Mob. Genet. Elem., 2, 247–252.

2. Doron, S., Melamed, S., Ofir, G., Leavitt, A., Lopatina, A., Keren, M., Amitai, G. and Sorek, R. (2018) Systematic discovery of antiphage defense systems in the microbial pangenome. Science, 359, eaar4120.

3. Millman, A., Melamed, S., Leavitt, A., Doron, S., Bernheim, A., Hör, J., Garb, J., Bechon, N., Brandis, A., Lopatina, A., et al. (2022) An expanded arsenal of immune systems that protect bacteria from phages. Cell Host Microbe, 30, 1556–1569.e5.

4. Gao, L.A., Wilkinson, M.E., Strecker, J., Makarova, K.S., Macrae, R.K., Koonin, E.V. and Zhang, F. (2022) Prokaryotic innate immunity through pattern recognition of conserved viral proteins. Science, 377, eabm4096.

5. Gao, L., Altae-Tran, H., Böhning, F., Makarova, K.S., Segel, M., Schmid-Burgk, J.L., Koob, J., Wolf, Y.I., Koonin, E.V. and Zhang, F. (2020) Diverse enzymatic activities mediate antiviral immunity in prokaryotes. Science, 369, 1077–1084.

6. Wein, T. and Sorek, R. (2022) Bacterial origins of human cell-autonomous innate immune mechanisms. Nat. Rev. Immunol., 10.1038/s41577-022-00705-4.

7. Culbertson, E.M. and Levin, T.C. (2023) Eukaryotic CD-NTase, STING, and viperin proteins evolved via domain shuffling, horizontal transfer, and ancient inheritance from prokaryotes. PLOS Biol., 21, e3002436.

8. Gizzi, A.S., Grove, T.L., Arnold, J.J., Jose, J., Jangra, R.K., Garforth, S.J., Du, Q., Cahill, S.M., Dulyaninova, N.G., Love, J.D., et al. (2018) A naturally occurring antiviral ribonucleotide encoded by the human genome. Nature, 558, 610–614.

9. Rivera-Serrano, E.E., Gizzi, A.S., Arnold, J.J., Grove, T.L., Almo, S.C. and Cameron, C.E. (2020) Viperin Reveals Its True Function. Annu. Rev. Virol., 7, 421–446.

10. Fenwick, M.K., Li, Y., Cresswell, P., Modis, Y. and Ealick, S.E. (2017) Structural studies of viperin, an antiviral radical SAM enzyme. Proc. Natl. Acad. Sci., 114, 6806–6811.

11. Bernheim, A., Millman, A., Ofir, G., Meitav, G., Avraham, C., Shomar, H., Rosenberg, M.M., Tal, N., Melamed, S., Amitai, G., et al. (2021) Prokaryotic viperins produce diverse antiviral molecules. Nature, 589, 120–124.

12. Bohmert, K., Camus, I., Bellini, C., Bouchez, D., Caboche, M. and Benning, C. (1998) AGO1 defines a novel locus of Arabidopsis controlling leaf development. EMBO J., 17, 170–180.

13. Hutvagner, G. and Simard, M.J. (2008) Argonaute proteins: Key players in RNA silencing. Nat. Rev. Mol. Cell Biol., 9, 22–32.

14. Zaremba, M., Dakineviciene, D., Golovinas, E., Zagorskaitė, E., Stankunas, E., Lopatina, A., Sorek, R., Manakova, E., Ruksenaite, A., Silanskas, A., et al. (2022) Short prokaryotic Argonautes provide defence against incoming mobile genetic elements through NAD+ depletion. Nat. Microbiol., 7, 1857–1869.

15. Koopal, B., Potocnik, A., Mutte, S.K., Aparicio-Maldonado, C., Lindhoud, S., Vervoort, J.J.M., Brouns, S.J.J. and Swarts, D.C. (2022) Short prokaryotic Argonaute systems trigger cell death upon detection of invading DNA. Cell, 10.1016/j.cell.2022.03.012.

16. Zeng, Z., Chen, Y., Pinilla-Redondo, R., Shah, S.A., Zhao, F., Wang, C., Hu, Z., Wu, C., Zhang, C., Whitaker, R.J., et al. (2022) A short prokaryotic Argonaute activates membrane effector to confer antiviral defense. Cell Host Microbe, 30, 930–943.e6.

17. Jolly, S.M., Gainetdinov, I., Jouravleva, K., Zhang, H., Strittmatter, L., Bailey, S.M., Hendricks, G.M., Dhabaria, A., Ueberheide, B. and Zamore, P.D. (2020) Thermus thermophilus Argonaute Functions in the Completion of DNA Replication. Cell, 182, 1545–1559.e18.

18. Dutta, R. and Inouye, M. (2000) GHKL, an emergent ATPase/kinase superfamily. Trends Biochem. Sci., 25, 24–28.

19. Bergerat, A., Gadelle, D. and Forterre, P. (1994) Purification of a DNA topoisomerase II from the hyperthermophilic archaeon Sulfolobus shibatae. J. Biol. Chem., 269, 27663–27669.

20. Nicastro, G.G., Burroughs, A.M., Iyer, L.M. and Aravind, L. (2023) Functionally comparable but evolutionarily distinct nucleotide-targeting effectors help identify conserved paradigms across diverse immune systems. Nucleic Acids Res., 10.1093/nar/gkad879.

21. Iyer, L.M., Abhiman, S. and Aravind, L. (2008) MutL homologs in restriction-modification systems and the origin of eukaryotic MORC ATPases. Biol. Direct, 3, 8.

22. Fendler, N.L., Ly, J., Welp, L., Lu, D., Schulte, F., Urlaub, H. and Vos, S.M. (2025) Identification and characterization of a human MORC2 DNA binding region that is required for gene silencing. Nucleic Acids Res., 53, gkae1273.

23. Tan, W., Park, J., Venugopal, H., Lou, J., Dias, P.S., Baldoni, P.L., Moon, K.-W., Dite, T.A., Keenan, C.R., Gurzau, A.D., et al. (2025) MORC2 is a phosphorylation-dependent DNA compaction machine. Nat. Commun., 16, 5606.

24. Karagöz, G.E., Duarte, A.M.S., Akoury, E., Ippel, H., Biernat, J., Morán Luengo, T., Radli, M., Didenko, T., Nordhues, B.A., Veprintsev, D.B., et al. (2014) Hsp90-Tau Complex Reveals Molecular Basis for Specificity in Chaperone Action. Cell, 156, 963–974.

25. Verba, K.A., Wang, R.Y.-R., Arakawa, A., Liu, Y., Shirouzu, M., Yokoyama, S. and Agard, D.A. (2016) Atomic structure of Hsp90-Cdc37-Cdk4 reveals that Hsp90 traps and stabilizes an unfolded kinase. Science, 352, 1542–1547.

26. Guarné, A., Ramon-Maiques, S., Wolff, E.M., Ghirlando, R., Hu, X., Miller, J.H. and Yang, W. (2004) Structure of the MutL C-terminal domain: a model of intact MutL and its roles in mismatch repair. EMBO J., 23, 4134–4145.

27. Ban, C. and Yang, W. (1998) Crystal Structure and ATPase Activity of MutL. Cell, 95, 541–552.

28. Xin, Y., Xian, R., Liu, C., Zhang, O., Rao, Z., Li, X. and Chen, Y. (2025) Direct trapping of the transport-segment DNA by the central domain of type IIA topoisomerases. Sci. Adv., 11, eadw2839.

29. Laponogov, I., Pan, X.-S., Veselkov, D.A., Skamrova, G.B., Umrekar, T.R., Fisher, L.M. and Sanderson, M.R. (2018) Trapping of the transport-segment DNA by the ATPase domains of a type II topoisomerase. Nat. Commun., 9, 2579.

30. Wigley, D.B., Davies, G.J., Dodson, E.J., Maxwell, A. and Dodson, G. (1991) Crystal structure of an N-terminal fragment of the DNA gyrase B protein. Nature, 351, 624–629.

31. Kim, H., Yen, L., Wongpalee, S.P., Kirshner, J.A., Mehta, N., Xue, Y., Johnston, J.B., Burlingame, A.L., Kim, J.K., Loparo, J.J., et al. (2019) The Gene-Silencing Protein MORC-1 Topologically Entraps DNA and Forms Multimeric Assemblies to Cause DNA Compaction. Mol. Cell, 75, 700–710.e6.

32. Pearl, L.H. and Prodromou, C. (2006) Structure and Mechanism of the Hsp90 Molecular Chaperone Machinery. Annu. Rev. Biochem., 75, 271–294.

33. Corbett, K.D. and Berger, J.M. (2004) Structure, Molecular Mechanisms, and Evolutionary Relationships in DNA Topoisomerases. Annu. Rev. Biophys. Biomol. Struct., 33, 95–118.

34. Kang, H.-G., Woo Choi, H., Von Einem, S., Manosalva, P., Ehlers, K., Liu, P.-P., Buxa, S.V., Moreau, M., Mang, H.-G., Kachroo, P., et al. (2012) CRT1 is a nuclear-translocated MORC endonuclease that participates in multiple levels of plant immunity. Nat. Commun., 3, 1297.

35. Kang, H.-G., Kuhl, J.C., Kachroo, P. and Klessig, D.F. (2008) CRT1, an Arabidopsis ATPase that Interacts with Diverse Resistance Proteins and Modulates Disease Resistance to Turnip Crinkle Virus. Cell Host Microbe, 3, 48–57.

36. Li, D.-Q., Nair, S.S. and Kumar, R. (2013) The MORC family: New epigenetic regulators of transcription and DNA damage response. Epigenetics, 8, 685–693.

37. Watson, M.L., Zinn, A.R., Inoue, N., Hess, K.D., Cobb, J., Handel, M.A., Halaban, R., Duchene, C.C., Albright, G.M. and Moreadith, R.W. (1998) Identification of *morc* (*microrchidia*), a mutation that results in arrest of spermatogenesis at an early meiotic stage in the mouse. Proc. Natl. Acad. Sci., 95, 14361–14366.

38. Mordret, E., Hervé, A., Tesson, F., Vaysset, H., Clabby, T., Loubat, A., Shomar, H., Planel, R., Lavenir, R., Cury, J., et al. (2026) Protein and genomic language models uncover the unexplored diversity of bacterial immunity. Science, 392, eadv8275.

39. DeWeirdt, P.C., Mahoney, E.M. and Laub, M.T. (2026) DefensePredictor: A machine learning model to discover prokaryotic immune systems. Science, 392, eadv7924.

40. Millman, A., Melamed, S., Amitai, G. and Sorek, R. (2020) Diversity and classification of cyclic-oligonucleotide-based anti-phage signalling systems. Nat. Microbiol., 5.

41. Tal, N. and Sorek, R. (2022) SnapShot: Bacterial immunity. Cell, 185, 578–578.e1.

42. Mestre, M.R., Gao, L.A., Shah, S.A., López-Beltrán, A., González-Delgado, A., Martínez-Abarca, F., Iranzo, J., Redrejo-Rodríguez, M., Zhang, F. and Toro, N. (2022) UG/Abi: a highly diverse family of prokaryotic reverse transcriptases associated with defense functions. Nucleic Acids Res., 50, 6084–6101.

43. Altae-Tran, H., Kannan, S., Suberski, A.J., Mears, K.S., Demircioglu, F.E., Moeller, L., Kocalar, S., Oshiro, R., Makarova, K.S., Macrae, R.K., et al. (2023) Uncovering the functional diversity of rare CRISPR-Cas systems with deep terascale clustering. Science, 382, eadi1910.

44. Blum, M., Andreeva, A., Florentino, L.C., Chuguransky, S.R., Grego, T., Hobbs, E., Pinto, B.L., Orr, A., Paysan-Lafosse, T., Ponamareva, I., et al. (2025) InterPro: the protein sequence classification resource in 2025. Nucleic Acids Res., 53, D444–D456.

45. Jones, P., Binns, D., Chang, H.-Y., Fraser, M., Li, W., McAnulla, C., McWilliam, H., Maslen, J., Mitchell, A., Nuka, G., et al. (2014) InterProScan 5: genome-scale protein function classification. Bioinforma. Oxf. Engl., 30, 1236–1240.

46. Steczkiewicz, K., Muszewska, A., Knizewski, L., Rychlewski, L. and Ginalski, K. (2012) Sequence, structure and functional diversity of PD-(D/E)XK phosphodiesterase superfamily. Nucleic Acids Res., 40, 7016–7045.

47. Payne, L.J., Todeschini, T.C., Wu, Y., Perry, B.J., Ronson, C.W., Fineran, P.C., Nobrega, F.L. and Jackson, S.A. (2021) Identification and classification of antiviral defence systems in bacteria and archaea with PADLOC reveals new system types. Nucleic Acids Res., 10.1093/nar/gkab883.

48. Liu, T., Gao, X., Chen, R., Tang, K., Liu, Z., Wang, P. and Wang, X. (2024) A nuclease domain fused to the Snf2 helicase confers antiphage defence in coral-associated *Halomonas meridiana*. Microb. Biotechnol., 17, e14524.

49. Fogg, J.M., Judge, A.K., Stricker, E., Chan, H.L. and Zechiedrich, L. (2021) Supercoiling and looping promote DNA base accessibility and coordination among distant sites. Nat. Commun., 12, 5683.

50. Tsugeki, R., Ditengou, F.A., Sumi, Y., Teale, W., Palme, K. and Okada, K. (2009) NO VEIN Mediates Auxin-Dependent Specification and Patterning in the *Arabidopsis* Embryo, Shoot, and Root. Plant Cell, 21, 3133–3151.

51. Tsugeki, R., Anicet Ditengou, F., Palme, K. and Okada, K. (2010) *NO VEIN* facilitates auxin-mediated development in Arabidopsis. Plant Signal. Behav., 5, 1249–1251.

52. Engert, J.C., Bérubé, P., Mercier, J., Doré, C., Lepage, P., Ge, B., Bouchard, J.-P., Mathieu, J., Melançon, S.B., Schalling, M., et al. (2000) ARSACS, a spastic ataxia common in northeastern Québec, is caused by mutations in a new gene encoding an 11.5-kb ORF. Nat. Genet., 24, 120–125.

53. Anderson, J.F., Siller, E. and Barral, J.M. (2010) The Sacsin Repeating Region (SRR): A Novel Hsp90-Related Supra-Domain Associated with Neurodegeneration. J. Mol. Biol., 400, 665–674.

54. The UniProt Consortium, Bateman, A., Martin, M.-J., Orchard, S., Magrane, M., Adesina, A., Ahmad, S., Bowler-Barnett, E.H., Bye-A-Jee, H., Carpentier, D., et al. (2025) UniProt: the Universal Protein Knowledgebase in 2025. Nucleic Acids Res., 53, D609–D617.

55. Ahmad, S., Jose da Costa Gonzales, L., Bowler-Barnett, E.H., Rice, D.L., Kim, M., Wijerathne, S., Luciani, A., Kandasaamy, S., Luo, J., Watkins, X., et al. (2025) The UniProt website API: facilitating programmatic access to protein knowledge. Nucleic Acids Res., 53, W547–W553.

56. Tesson, F., Planel, R., Egorov, A.A., Georjon, H., Vaysset, H., Brancotte, B., Néron, B., Mordret, E., Atkinson, G.C., Bernheim, A., et al. (2024) A Comprehensive Resource for Exploring Antiphage Defense: DefenseFinder Webservice, Wiki and Databases. Peer Community J., 4, e91.

57. Fu, L., Niu, B., Zhu, Z., Wu, S. and Li, W. (2012) CD-HIT: accelerated for clustering the next-generation sequencing data. Bioinformatics, 28, 3150–3152.

58. Katoh, K. and Standley, D.M. (2013) MAFFT Multiple Sequence Alignment Software Version 7: Improvements in Performance and Usability. Mol. Biol. Evol., 30, 772–780.

59. Soding, J., Biegert, A. and Lupas, A.N. (2005) The HHpred interactive server for protein homology detection and structure prediction. Nucleic Acids Res., 33, W244–W248.

60. Steenwyk, J.L., Buida, T.J., Li, Y., Shen, X.-X. and Rokas, A. (2020) ClipKIT: A multiple sequence alignment trimming software for accurate phylogenomic inference. PLOS Biol., 18, e3001007.

61. Finn, R.D., Clements, J. and Eddy, S.R. (2011) HMMER web server: interactive sequence similarity searching. Nucleic Acids Res., 39, W29–W37.

62. Capella-Gutiérrez, S., Silla-Martínez, J.M. and Gabaldón, T. (2009) trimAl: a tool for automated alignment trimming in large-scale phylogenetic analyses. Bioinformatics, 25, 1972–1973.

63. Nguyen, L.-T., Schmidt, H.A., Von Haeseler, A. and Minh, B.Q. (2015) IQ-TREE: A Fast and Effective Stochastic Algorithm for Estimating Maximum-Likelihood Phylogenies. Mol. Biol. Evol., 32, 268–274.

64. Bianchini, G. and Sánchez-Baracaldo, P. (2024) TREEVIEWER: Flexible, modular software to visualise and manipulate phylogenetic trees. Ecol. Evol., 14, e10873.

65. Allen, A.M.B. and Maxwell, A. (2024) Phylogenetic distribution of DNA topoisomerase VI and its distinction from SPO11. NAR Genomics Bioinforma., 6, lqae085.

66. Wong, T.K.F., Ly-Trong, N., Ren, H., Demotte, P., Baños, H., Roger, A.J., Susko, E., Bielow, C., De Maio, N., Goldman, N., et al. (2026) IQ-TREE 3: phylogenomic inference software using complex evolutionary models. Mol. Biol. Evol., 43, msag117.

67. Hoang, D.T., Chernomor, O., Von Haeseler, A., Minh, B.Q. and Vinh, L.S. (2018) UFBoot2: Improving the Ultrafast Bootstrap Approximation. Mol. Biol. Evol., 35, 518–522.

68. Abramson, J., Adler, J., Dunger, J., Evans, R., Green, T., Pritzel, A., Ronneberger, O., Willmore, L., Ballard, A.J., Bambrick, J., et al. (2024) Accurate structure prediction of biomolecular interactions with AlphaFold 3. Nature, 10.1038/s41586-024-07487-w.

69. Lau, A.M., Bordin, N., Kandathil, S.M., Sillitoe, I., Waman, V.P., Wells, J., Orengo, C.A. and Jones, D.T. (2024) Exploring structural diversity across the protein universe with The Encyclopedia of Domains. Science, 386, eadq4946.

70. Van Kempen, M., Kim, S.S., Tumescheit, C., Mirdita, M., Lee, J., Gilchrist, C.L.M., Söding, J. and Steinegger, M. (2023) Fast and accurate protein structure search with Foldseek. Nat. Biotechnol., 10.1038/s41587-023-01773-0.

71. Gilchrist, C.L.M., Mirdita, M. and Steinegger, M. (2026) Multiple protein structure alignment at scale with FoldMason. Science, 391, 485–488.

72. Zhang, Y. (2005) TM-align: a protein structure alignment algorithm based on the TM-score. Nucleic Acids Res., 33, 2302–2309.

73. Kilgore, H.R., Chinn, I., Mikhael, P.G., Mitnikov, I., Van Dongen, C., Zylberberg, G., Afeyan, L., Banani, S.F., Wilson-Hawken, S., Lee, T.I., et al. (2025) Protein codes promote selective subcellular compartmentalization. Science, 387, 1095–1101.

74. Oberg, N., Zallot, R. and Gerlt, J.A. (2023) EFI-EST, EFI-GNT, and EFI-CGFP: Enzyme Function Initiative (EFI) Web Resource for Genomic Enzymology Tools. J. Mol. Biol., 435, 168018.

75. Tesson, F., Hervé, A., Mordret, E., Touchon, M., D’Humières, C., Cury, J. and Bernheim, A. (2022) Systematic and quantitative view of the antiviral arsenal of prokaryotes. Nat. Commun., 13, 2561.

76. Eddy, S.R. (1998) Profile hidden Markov models. Bioinformatics, 14, 755–763.

77. Makarova, K.S., Wolf, Y.I., van der Oost, J. and Koonin, E.V. (2009) Prokaryotic homologs of Argonaute proteins are predicted to function as key components of a novel system of defense against mobile genetic elements. Biol. Direct, 4, 29.

78. Ruggiero, M.A., Gordon, D.P., Orrell, T.M., Bailly, N., Bourgoin, T., Brusca, R.C., Cavalier-Smith, T., Guiry, M.D. and Kirk, P.M. (2015) A Higher Level Classification of All Living Organisms. PLOS ONE, 10, e0119248.

79. Santiago-Frangos, A., Buyukyoruk, M., Wiegand, T., Krishna, P. and Wiedenheft, B. (2021) Distribution and phasing of sequence motifs that facilitate CRISPR adaptation. Curr. Biol., 10.1016/j.cub.2021.05.068.

80. Buyukyoruk, M., Krishna, P., Santiago-Frangos, A. and Wiedenheft, B. (2025) Discovery of Diverse CRISPR Leader Motifs, Putative Functions, and Applications for Enhanced CRISPR Detection and Subtype Annotation. CRISPR J., 10.1089/crispr.2024.0093.

81. Nuñez, J.K., Bai, L., Harrington, L.B., Hinder, T.L. and Doudna, J.A. (2016) CRISPR Immunological Memory Requires a Host Factor for Specificity. Mol. Cell, 62, 824–833.

82. Holm, L. (2022) Dali server: structural unification of protein families. Nucleic Acids Res., 50, W210–W215.

83. Ban, C., Junop, M. and Yang, W. (1999) Transformation of MutL by ATP Binding and Hydrolysis. Cell, 97, 85–97.

84. Corbett, K.D. and Berger, J.M. (2003) Structure of the topoisomerase VI-B subunit: implications for type II topoisomerase mechanism and evolution. EMBO J., 22, 151–163.

85. Oestergaard, V.H., Bjergbaek, L., Skouboe, C., Giangiacomo, L., Knudsen, B.R. and Andersen, A.H. (2004) The Transducer Domain Is Important for Clamp Operation in Human DNA Topoisomerase IIα. J. Biol. Chem., 279, 1684–1691.

86. Gruszczyk, J., Grandvuillemin, L., Lai-Kee-Him, J., Paloni, M., Savva, C.G., Germain, P., Grimaldi, M., Boulahtouf, A., Kwong, H.-S., Bous, J., et al. (2022) Cryo-EM structure of the agonist-bound Hsp90-XAP2-AHR cytosolic complex. Nat. Commun., 13, 7010.

87. Lavery, L.A., Partridge, J.R., Ramelot, T.A., Elnatan, D., Kennedy, M.A. and Agard, D.A. (2014) Structural Asymmetry in the Closed State of Mitochondrial Hsp90 (TRAP1) Supports a Two-Step ATP Hydrolysis Mechanism. Mol. Cell, 53, 330–343.

88. Evans, R., O’Neill, M., Pritzel, A., Antropova, N., Senior, A., Green, T., Žídek, A., Bates, R., Blackwell, S., Yim, J., et al. (2021) Protein complex prediction with AlphaFold-Multimer. 10.1101/2021.10.04.463034.

89. Blatch, G.L. and Lässle, M. (1999) The tetratricopeptide repeat: a structural motif mediating protein-protein interactions. BioEssays, 21, 932–939.

90. Zeytuni, N. and Zarivach, R. (2012) Structural and Functional Discussion of the Tetra-Trico-Peptide Repeat, a Protein Interaction Module. Structure, 20, 397–405.

91. Reinking, H.K., Hofmann, K. and Stingele, J. (2020) Function and evolution of the DNA-protein crosslink proteases Wss1 and SPRTN. DNA Repair, 88, 102822.

92. Duxin, J.P., Dewar, J.M., Yardimci, H. and Walter, J.C. (2014) Repair of a DNA-Protein Crosslink by Replication-Coupled Proteolysis. Cell, 159, 346–357.

93. Vallee, B.L. and Auld, D.S. (1990) Zinc coordination, function, and structure of zinc enzymes and other proteins. Biochemistry, 29, 5647–5659.

94. Jongeneel, C.V., Bouvier, J. and Bairoch, A. (1989) A unique signature identifies a family of zinc-dependent metallopeptidases. FEBS Lett., 242, 211–214.

95. Murphy, J.M., Mace, P.D. and Eyers, P.A. (2017) Live and let die: insights into pseudoenzyme mechanisms from structure. Curr. Opin. Struct. Biol., 47, 95–104.

96. Jeffery, C.J. (2019) The demise of catalysis, but new functions arise: pseudoenzymes as the phoenixes of the protein world. Biochem. Soc. Trans., 47, 371–379.

97. Adrain, C. and Freeman, M. (2012) New lives for old: evolution of pseudoenzyme function illustrated by iRhoms. Nat. Rev. Mol. Cell Biol., 13, 489–498.

98. Zettl, M., Adrain, C., Strisovsky, K., Lastun, V. and Freeman, M. (2011) Rhomboid Family Pseudoproteases Use the ER Quality Control Machinery to Regulate Intercellular Signaling. Cell, 145, 79–91.

99. Klepikova, A.V., Kasianov, A.S., Gerasimov, E.S., Logacheva, M.D. and Penin, A.A. (2016) A high resolution map of the *Arabidopsis thaliana* developmental transcriptome based on RNA –seq profiling. Plant J., 88, 1058–1070.

100. Loenen, W.A.M. and Raleigh, E.A. (2014) The other face of restriction: modification-dependent enzymes. Nucleic Acids Res., 42, 56–69.

101. Jeske, M., Müller, C.W. and Ephrussi, A. (2017) The LOTUS domain is a conserved DEAD-box RNA helicase regulator essential for the recruitment of Vasa to the germ plasm and nuage. Genes Dev., 31, 939–952.

102. Ding, D., Wei, C., Dong, K., Liu, J., Stanton, A., Xu, C., Min, J., Hu, J. and Chen, C. (2020) LOTUS domain is a novel class of G-rich and G-quadruplex RNA binding domain. Nucleic Acids Res., 48, 9262–9272.

103. Callebaut, I. and Mornon, J.-P. (2010) LOTUS, a new domain associated with small RNA pathways in the germline. Bioinformatics, 26, 1140–1144.

104. Jeske, M., Bordi, M., Glatt, S., Müller, S., Rybin, V., Müller, C.W. and Ephrussi, A. (2015) The Crystal Structure of the Drosophila Germline Inducer Oskar Identifies Two Domains with Distinct Vasa Helicase– and RNA-Binding Activities. Cell Rep., 12, 587–598.

105. Pedersen, L.C., Yi, M., Pedersen, L.G. and Kaminski, A.M. (2022) From Steroid and Drug Metabolism to Glycobiology, Using Sulfotransferase Structures to Understand and Tailor Function. Drug Metab. Dispos., 50, 1027–1041.

106. Krishna, S.S. (2003) Structural classification of zinc fingers: SURVEY AND SUMMARY. Nucleic Acids Res., 31, 532–550.

107. Luo, Y., Na, Z. and Slavoff, S.A. (2018) P-Bodies: Composition, Properties, and Functions. Biochemistry, 57, 2424–2431.

108. Chen, S., Cai, Q., Peng, H., Yang, Y., Wu, Z. and Zhang, M. (2026) Native postsynaptic density is a functional condensate formed via phase separation. Cell Rep., 45, 116723.

109. Bayés, À., Collins, M.O., Croning, M.D.R., Van De Lagemaat, L.N., Choudhary, J.S. and Grant, S.G.N. (2012) Comparative Study of Human and Mouse Postsynaptic Proteomes Finds High Compositional Conservation and Abundance Differences for Key Synaptic Proteins. PLoS ONE, 7, e46683.

110. Sheng, M. and Hoogenraad, C.C. (2007) The Postsynaptic Architecture of Excitatory Synapses: A More Quantitative View. Annu. Rev. Biochem., 76, 823–847.

111. Bastian, F.B., Cammarata, A.B., Carsanaro, S., Detering, H., Huang, W.-T., Joye, S., Niknejad, A., Nyamari, M., Mendes de Farias, T., Moretti, S., et al. (2025) Bgee in 2024: focus on curated single-cell RNA-seq datasets, and query tools. Nucleic Acids Res., 53, D878–D885.

112. Rousset, F. and Sorek, R. (2023) The evolutionary success of regulated cell death in bacterial immunity. Curr. Opin. Microbiol., 74, 102312.

113. Jaciuk, M., Swuec, P., Gaur, V., Kasprzak, J.M., Renault, L., Dobrychłop, M., Nirwal, S., Bujnicki, J.M., Costa, A. and Nowotny, M. (2020) A combined structural and biochemical approach reveals translocation and stalling of UvrB on the DNA lesion as a mechanism of damage verification in bacterial nucleotide excision repair. DNA Repair, 85, 102746.

114. Mocquet, V. and Fiorini, F. (2026) UPF1 at Work: Structural and Mechanistic Insights Into a Master Regulator of Nonsense-Mediated mRNA Decay. J. Mol. Biol., 10.1016/j.jmb.2026.169913.

115. Ahmad, S. and Hur, S. (2015) Helicases in Antiviral Immunity: Dual Properties as Sensors and Effectors. Trends Biochem. Sci., 40, 576–585.

116. Tuck, O.T., Adler, B.A., Armbruster, E.G., Lahiri, A., Hu, J.J., Zhou, J., Pogliano, J. and Doudna, J.A. (2024) Genome integrity sensing by the broad-spectrum Hachiman antiphage defense complex. Cell, 187, 6914–6928.e20.

117. Ragucci, A.E., Antine, S.P., Leviss, E.M., Mooney, S.E., Garcia, J.M., Shyrokova, L., Hauryliuk, V., Lee, A.S.Y. and Kranzusch, P.J. (2026) Nuclease–NTPase antiphage defence systems use conserved molecular features to control bacterial immunity. Nat. Microbiol., 11, 1424–1436.

118. Shen, B.W., Heiter, D.F., Chan, S.-H., Wang, H., Xu, S.-Y., Morgan, R.D., Wilson, G.G. and Stoddard, B.L. (2010) Unusual Target Site Disruption by the Rare-Cutting HNH Restriction Endonuclease PacI. Structure, 18, 734–743.

119. Pastor, M., Czapinska, H., Helbrecht, I., Krakowska, K., Lutz, T., Xu, S. and Bochtler, M. (2021) Crystal structures of the EVE-HNH endonuclease VcaM4I in the presence and absence of DNA. Nucleic Acids Res., 49, 1708–1723.

120. Yang, J.-Y., Deng, X.-Y., Li, Y.-S., Ma, X.-C., Feng, J.-X., Yu, B., Chen, Y., Luo, Y.-L., Wang, X., Chen, M.-L., et al. (2018) Structure of Schlafen13 reveals a new class of tRNA/rRNA-targeting RNase engaged in translational control. Nat. Commun., 9, 1165.

121. Hou, P., Hao, W., Qin, B., Li, M., Zhao, R. and Cui, S. (2023) Structural and biochemical characterization of Schlafen11 N-terminal domain. Nucleic Acids Res., 51, 7053–7070.

122. Perez Taboada, V., Wu, Y., Cassidy, R., Medvedev, K.E., Loeff, L., Nemudraia, A. and Nemudryi, A. (2026) Bacterial Schlafen proteins mediate phage defence. Nat. Microbiol., 10.1038/s41564-026-02277-8.

123. Steens, J.A., Bravo, J.P.K., Salazar, C.R.P., Yildiz, C., Amieiro, A.M., Köstlbacher, S., Prinsen, S.H.P., Andres, A.S., Patinios, C., Bardis, A., et al. (2024) Type III-B CRISPR-Cas cascade of proteolytic cleavages. Science, 383, 512–519.

124. Hu, C., van Beljouw, S.P.B., Nam, K.H., Schuler, G., Ding, F., Cui, Y., Rodríguez-Molina, A., Haagsma, A.C., Valk, M., Pabst, M., et al. (2022) Craspase is a CRISPR RNA-guided, RNA-activated protease. Science, 377, 1278–1285.

125. Tuck, O.T., Hu, J.J., Lopez, S.C., Adler, B.A., O’Brien, C.E., Hsieh, K., Meredith, C., Loi, K.J., Yoon, P.H., Doherty, E.E., et al. (2026) Recurrent acquisition of nuclease-protease pairs in antiviral immunity. Science, 391, 195–201.

126. Chen, H., Ning, X. and Jiang, Z. (2017) Caspases control antiviral innate immunity. Cell. Mol. Immunol., 14, 736–747.

127. Draper, D.E. and Reynaldo, L.P. (1999) RNA binding strategies of ribosomal proteins. Nucleic Acids Res., 27, 381–388.

128. Roper, M., Simonin, A., Hickey, P.C., Leeder, A. and Glass, N.L. (2013) Nuclear dynamics in a fungal chimera. Proc. Natl. Acad. Sci., 110, 12875–12880.

129. Roca, M.G., Kuo, H.-C., Lichius, A., Freitag, M. and Read, N.D. (2010) Nuclear Dynamics, Mitosis, and the Cytoskeleton during the Early Stages of Colony Initiation in Neurospora crassa. Eukaryot. Cell, 9, 1171–1183.

130. Smith, M.L., Micali, O.C., Hubbard, S.P., Mir-Rashed, N., Jacobson, D.J. and Glass, N.L. (2000) Vegetative Incompatibility in the *het-6* Region of *Neurospora crassa* Is Mediated by Two Linked Genes. Genetics, 155, 1095–1104.

131. Loog, M. and Morgan, D.O. (2005) Cyclin specificity in the phosphorylation of cyclin-dependent kinase substrates. Nature, 434, 104–108.

132. Francis, V., Alshafie, W., Kumar, R., Girard, M., Brais, B. and McPherson, P.S. (2022) The ARSACS disease protein sacsin controls lysosomal positioning and reformation by regulating microtubule dynamics. J. Biol. Chem., 298, 102320.

133. Bradshaw, T.Y., Romano, L.E.L., Duncan, E.J., Nethisinghe, S., Abeti, R., Michael, G.J., Giunti, P., Vermeer, S. and Chapple, J.P. (2016) A reduction in Drp1-mediated fission compromises mitochondrial health in autosomal recessive spastic ataxia of Charlevoix Saguenay. Hum. Mol. Genet., 25, 3232–3244.

134. Pillon, M.C., Gordon, J., Frazier, M.N. and Stanley, R.E. (2021) HEPN RNases–an emerging class of functionally distinct RNA processing and degradation enzymes. Crit. Rev. Biochem. Mol. Biol., 56, 88–108.

135. Gadelle, D., Filée, J., Buhler, C. and Forterre, P. (2003) Phylogenomics of type II DNA topoisomerases. BioEssays, 25, 232–242.

136. Fairman-Williams, M.E., Guenther, U.-P. and Jankowsky, E. (2010) SF1 and SF2 helicases: family matters. Curr. Opin. Struct. Biol., 20, 313–324.

137. Byrd, A., K. (2012) Superfamily 2 helicases. Front. Biosci., 17, 2070.

138. Singleton, M.R., Dillingham, M.S. and Wigley, D.B. (2007) Structure and Mechanism of Helicases and Nucleic Acid Translocases. Annu. Rev. Biochem., 76, 23–50.

139. Gorbalenya, A.E. and Koonin, E.V. (1993) Helicases: amino acid sequence comparisons and structure-function relationships. Curr. Opin. Struct. Biol., 3, 419–429.

140. Maestri, A., Pons, B.J., Pursey, E., Chong, C.E., Gandon, S., Custodio, R., Olina, A., Agapov, A., Chisnall, M.A.W., Grasso, A., et al. (2024) The bacterial defense system MADS interacts with CRISPR-Cas to limit phage infection and escape. Cell Host Microbe, 32, 1412–1426.e11.

141. Luyten, Y.A., Hausman, D.E., Young, J.C., Doyle, L.A., Higashi, K.M., Ubilla-Rodriguez, N.C., Lambert, A.R., Arroyo, C.S., Forsberg, K.J., Morgan, R.D., et al. (2022) Identification and characterization of the WYL BrxR protein and its gene as separable regulatory elements of a BREX phage restriction system. Nucleic Acids Res., 50, 5171–5190.

142. Picton, D.M., Harling-Lee, J.D., Duffner, S.J., Went, S.C., Morgan, R.D., Hinton, J.C.D. and Blower, T.R. (2022) A widespread family of WYL-domain transcriptional regulators co-localizes with diverse phage defence systems and islands. Nucleic Acids Res., 50, 5191–5207.

143. Drobiazko, A., Adams, M.C., Skutel, M., Potekhina, K., Kotovskaya, O., Trofimova, A., Matlashov, M., Yatselenko, D., Maxwell, K.L., Blower, T.R., et al. (2025) Molecular basis of foreign DNA recognition by BREX anti-phage immunity system. Nat. Commun., 16, 1825.

144. Goldfarb, T., Sberro, H., Weinstock, E., Cohen, O., Doron, S., Charpak-Amikam, Y., Afik, S., Ofir, G. and Sorek, R. (2015) BREX is a novel phage resistance system widespread in microbial genomes. EMBO J., 34, 169–183.

145. Readshaw, J.J., Doyle, L.A., Puiu, M., Kelly, A., Nelson, A., Kaiser, A.J., McGuire, S.F., Peralta Acosta, J., Smith, D.L., Stoddard, B.L., et al. (2025) PglZ from Type I BREX phage defence systems is a metal-dependent nuclease that forms a sub-complex with BrxB. Nucleic Acids Res., 53, gkaf540.

146. Murray, N.E. (2000) Type I Restriction Systems: Sophisticated Molecular Machines (a Legacy of Bertani and Weigle). Microbiol. Mol. Biol. Rev., 64, 412–434.

147. Cheng, R., Huang, F., Lu, X., Yan, Y., Yu, B., Wang, X. and Zhu, B. (2023) Prokaryotic Gabija complex senses and executes nucleotide depletion and DNA cleavage for antiviral defense. Cell Host Microbe, 31, 1331–1344.e5.

148. Cheng, R., Huang, F., Wu, H., Lu, X., Yan, Y., Yu, B., Wang, X. and Zhu, B. (2021) A nucleotide-sensing endonuclease from the Gabija bacterial defense system. Nucleic Acids Res., 49, 5216–5229.

149. Ofir, G., Melamed, S., Sberro, H., Mukamel, Z., Silverman, S., Yaakov, G., Doron, S. and Sorek, R. (2017) DISARM is a widespread bacterial defence system with broad anti-phage activities. Nat. Microbiol., 3, 90–98.

150. Bravo, J.P.K., Aparicio-Maldonado, C., Nobrega, F.L., Brouns, S.J.J. and Taylor, D.W. (2022) Structural basis for broad anti-phage immunity by DISARM. Nat. Commun., 13, 2987.

151. Gerdes, K. (2025) Mono– and multidomain defense toxins of the RelE/ParE superfamily. mBio, 16, e00258–25.

152. Mogila, I., Tamulaitiene, G., Keda, K., Timinskas, A., Ruksenaite, A., Sasnauskas, G., Venclovas, Č., Siksnys, V. and Tamulaitis, G. (2023) Ribosomal stalk-captured CARF-RelE ribonuclease inhibits translation following CRISPR signaling. Science, 382, 1036–1041.

153. Kervestin, S. and Jacobson, A. (2012) NMD: a multifaceted response to premature translational termination. Nat. Rev. Mol. Cell Biol., 13, 700–712.

154. Senissar, M., Manav, M.C. and Brodersen, D.E. (2017) Structural conservation of the PIN domain active site across all domains of life. Protein Sci., 26, 1474–1492.

155. Wang, F., Zhao, P., Bi, X., Zheng, R., Tian, X., Xu, J., Jiang, S., Li, G., Shen, Y. and She, Q. (2025) Cyclic tetraadenylate binding induces dimerization of protein dimers to activate a CRISPR-associated PIN nuclease. Nucleic Acids Res., 53, gkaf744.

156. Makarova, K.S., Wolf, Y.I., Snir, S. and Koonin, E.V. (2011) Defense Islands in Bacterial and Archaeal Genomes and Prediction of Novel Defense Systems. J. Bacteriol., 193, 6039–6056.

157. Matson, S.W. and Robertson, A.B. (2006) The UvrD helicase and its modulation by the mismatch repair protein MutL. Nucleic Acids Res., 34, 4089–4097.

158. Xu, W., Beebe, K., Chavez, J.D., Boysen, M., Lu, Y., Zuehlke, A.D., Keramisanou, D., Trepel, J.B., Prodromou, C., Mayer, M.P., et al. (2019) Hsp90 middle domain phosphorylation initiates a complex conformational program to recruit the ATPase-stimulating cochaperone Aha1. Nat. Commun., 10, 2574.

159. Grenert, J.P., Johnson, B.D. and Toft, D.O. (1999) The Importance of ATP Binding and Hydrolysis by Hsp90 in Formation and Function of Protein Heterocomplexes. J. Biol. Chem., 274, 17525– 17533.

160. Qu, X., Wang, S., Zhao, S., Wan, C., Xu, W. and Huang, C. (2024) The dynamic triage interplay of Hsp90 with its chaperone cycle and client binding. Nat. Commun., 15, 10661.

161. Corbett, K.D. and Berger, J.M. (2005) Structural Dissection of ATP Turnover in the Prototypical GHL ATPase TopoVI. Structure, 13, 873–882.

162. Payne, L.J., Meaden, S., Mestre, M.R., Palmer, C., Toro, N., Fineran, P.C. and Jackson, S.A. (2022) PADLOC: a web server for the identification of antiviral defence systems in microbial genomes. Nucleic Acids Res., 50, W541–W550.

163. Makarova, K.S., Wolf, Y.I., Iranzo, J., Shmakov, S.A., Alkhnbashi, O.S., Brouns, S.J.J.J., Charpentier, E., Cheng, D., Haft, D.H., Horvath, P., et al. (2020) Evolutionary classification of CRISPR–Cas systems: a burst of class 2 and derived variants. Nat. Rev. Microbiol., 18, 67–83.

164. Cui, Y., Dai, Z., Ouyang, Y., Fu, C., Wang, Y., Chen, X., Yang, K., Zheng, S., Wang, W., Tao, P., et al. (2025) Bacterial Hachiman complex executes DNA cleavage for antiphage defense. Nat. Commun., 16, 2604.

165. Burman, N., Belukhina, S., Depardieu, F., Wilkinson, R.A., Skutel, M., Santiago-Frangos, A., Graham, A.B., Livenskyi, A., Chechenina, A., Morozova, N., et al. (2024) Viral proteins activate PARIS-mediated tRNA degradation and viral tRNAs rescue infection Biochemistry.

166. Tesson, F., Huiting, E., Wei, L., Ren, J., Johnson, M., Planel, R., Cury, J., Feng, Y., Bondy-Denomy, J. and Bernheim, A. (2024) Exploring the diversity of anti-defense systems across prokaryotes, phages and mobile genetic elements. Nucleic Acids Res., 10.1093/nar/gkae1171.

167. Mayo-Muñoz, D., Pinilla-Redondo, R., Birkholz, N. and Fineran, P.C. (2023) A host of armor: Prokaryotic immune strategies against mobile genetic elements. Cell Rep., 42, 112672.

168. Georjon, H. and Bernheim, A. (2023) The highly diverse antiphage defence systems of bacteria. Nat. Rev. Microbiol., 21, 686–700.

169. Makarova, K.S., Timinskas, A., Wolf, Y.I., Gussow, A.B., Siksnys, V., Venclovas, Č. and Koonin, E.V. (2020) Evolutionary and functional classification of the CARF domain superfamily, key sensors in prokaryotic antivirus defense. Nucleic Acids Res., 10.1093/nar/gkaa635.

170. Li, H., Tan, Y., Basu, D., Corbett, K.D. and Zhang, D. (2025) Unveiling the multifaceted domain polymorphism of the Menshen antiphage system. Nucleic Acids Res., 53, gkaf357.

171. Rakesh, S. and Krishnan, A. (2026) Expanding the landscape of BREX diversity: uncovering multi-layered functional frameworks and identification of novel BREX-related defense systems. Nucleic Acids Res., 54, gkag035.

172. Paoletti, M. and Clavé, C. (2007) The Fungus-Specific HET Domain Mediates Programmed Cell Death in *Podospora anserina*. Eukaryot. Cell, 6, 2001–2008.

173. Haerty, W. and Golding, G.B. (2010) Low-complexity sequences and single amino acid repeats: not just “junk” peptide sequences. Genome, 53, 753–762.

174. Eme, L., Gentekaki, E., Curtis, B., Archibald, J.M. and Roger, A.J. (2017) Lateral Gene Transfer in the Adaptation of the Anaerobic Parasite Blastocystis to the Gut. Curr. Biol., 27, 807–820.

175. Gabaldón, T. (2020) Patterns and impacts of nonvertical evolution in eukaryotes: a paradigm shift. Ann. N. Y. Acad. Sci., 1476, 78–92.

176. Leger, M.M., Eme, L., Stairs, C.W. and Roger, A.J. (2018) Demystifying Eukaryote Lateral Gene Transfer (Response to Martin 2017 DOI: 10.1002/bies.201700115). BioEssays, 40, 1700242.

177. Richards, T.A., Soanes, D.M., Jones, M.D.M., Vasieva, O., Leonard, G., Paszkiewicz, K., Foster, P.G., Hall, N. and Talbot, N.J. (2011) Horizontal gene transfer facilitated the evolution of plant parasitic mechanisms in the oomycetes. Proc. Natl. Acad. Sci., 108, 15258–15263.

178. Srivastava, M., Begovic, E., Chapman, J., Putnam, N.H., Hellsten, U., Kawashima, T., Kuo, A., Mitros, T., Salamov, A., Carpenter, M.L., et al. (2008) The Trichoplax genome and the nature of placozoans. Nature, 454, 955–960.

