## Supplementary Figures for "The bacterial antiphage protein Tiamat is an ATPase and DNase whose homologs are found across the tree of life"

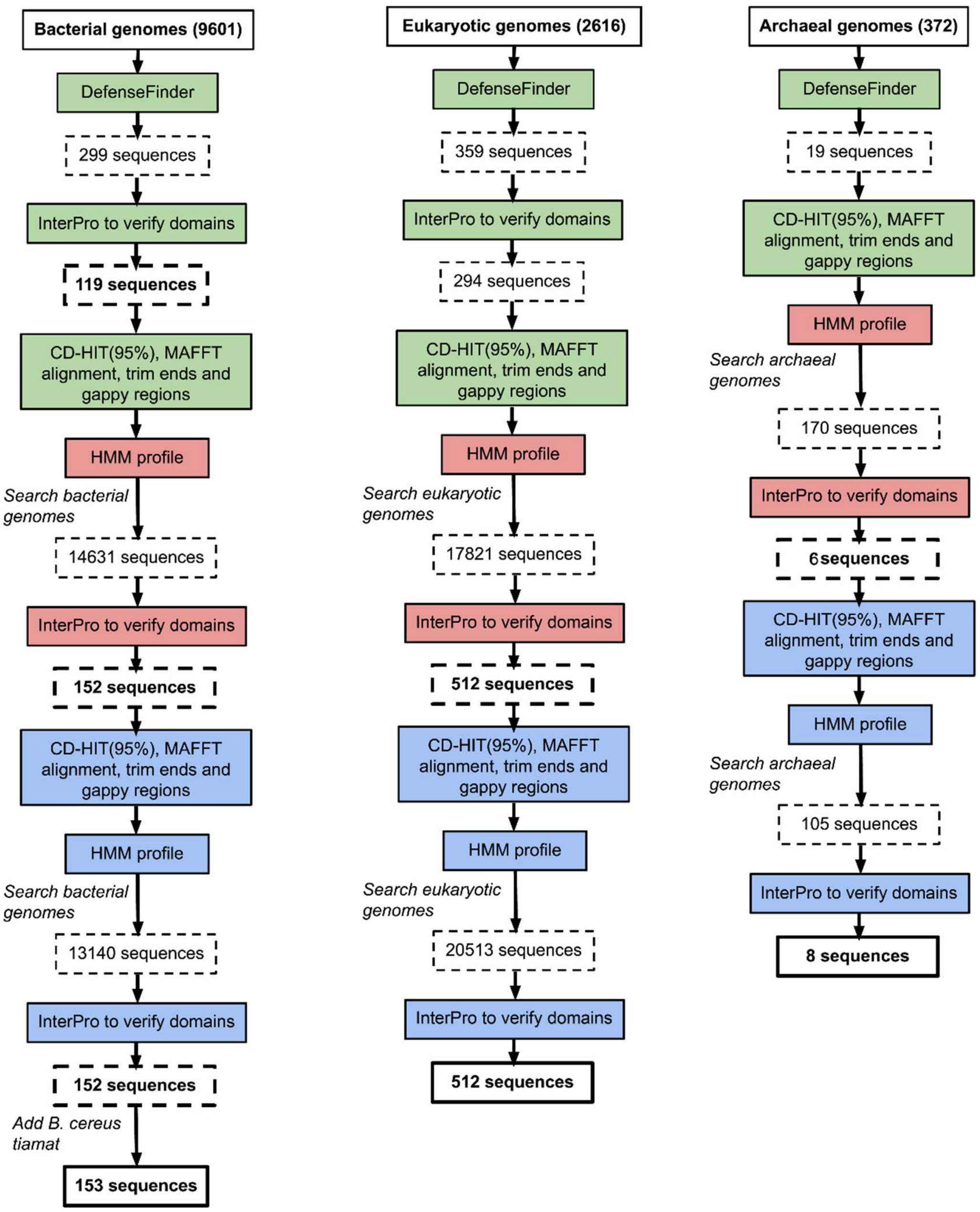


**Figure S1. Initial workflow for identifying and curating Tiamat homologs across bacterial, eukaryotic, and archaeal proteomes.** Flow charts summarizing the iterative bioinformatic pipeline used to identify, cluster, and verify Tiamat homologs across Bacterial (9,601 proteomes), Eukaryotic (2,616 proteomes), and Archaeal (372 proteomes) datasets from the UniProt reference proteome database. For each domain of life, candidate sequences initially detected by DefenseFinder were manually inspected using InterProScan to confirm the presence of expected domains. Verified sequences were clustered with CD-HIT

(95%), aligned with MAFFT, and trimmed to remove gappy or poorly aligned regions before building an initial HMM profile. The resulting profiles were then searched against their respective genomic datasets to recover additional homologs. Newly identified sequences were again filtered by domain composition and redundancy, and used to generate refined HMM profiles for a second search round. The process was repeated until no new sequences were found (2 iterations). Final representative sets of 153 bacterial, 512 eukaryotic, and 8 archaeal Tiamat homologs were retained for downstream analyses.

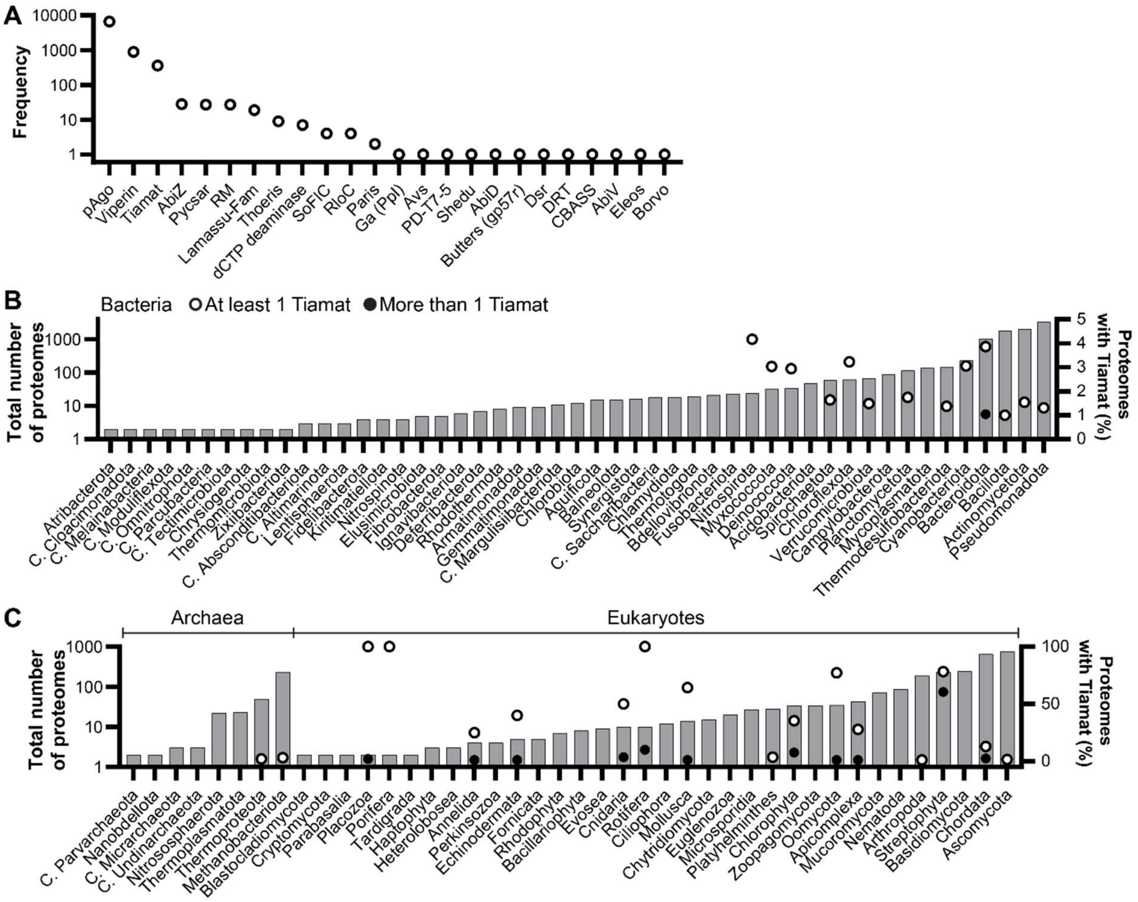


**Figure S2. Tiamat homologs are highly prevalent in distinct eukaryotic taxonomic groups.** (**A**) Number of bacterial and archaeal antiviral protein homologs identified in an initial search against UniProt reference eukaryotic proteomes. (**B**) The number of proteomes searched (left y-axis) from specific bacterial phyla is shown (UniProt reference proteomes). The percentage of proteomes with a minimum of 1 Tiamat homolog (open circle), and more than 1 Tiamat homolog (closed circle) are indicated (right y-axis). (**C**) The number of proteomes searched (left y-axis) from specific archaeal and eukaryotic phyla is shown. The percentage of proteomes with a minimum of 1 Tiamat homolog (open circle), and more than 1 Tiamat homolog (closed circle) are indicated (right y-axis). For clarity, only phyla for which more than 1 proteome was analyzed are shown, and zero frequency values are not plotted in panels B and C.

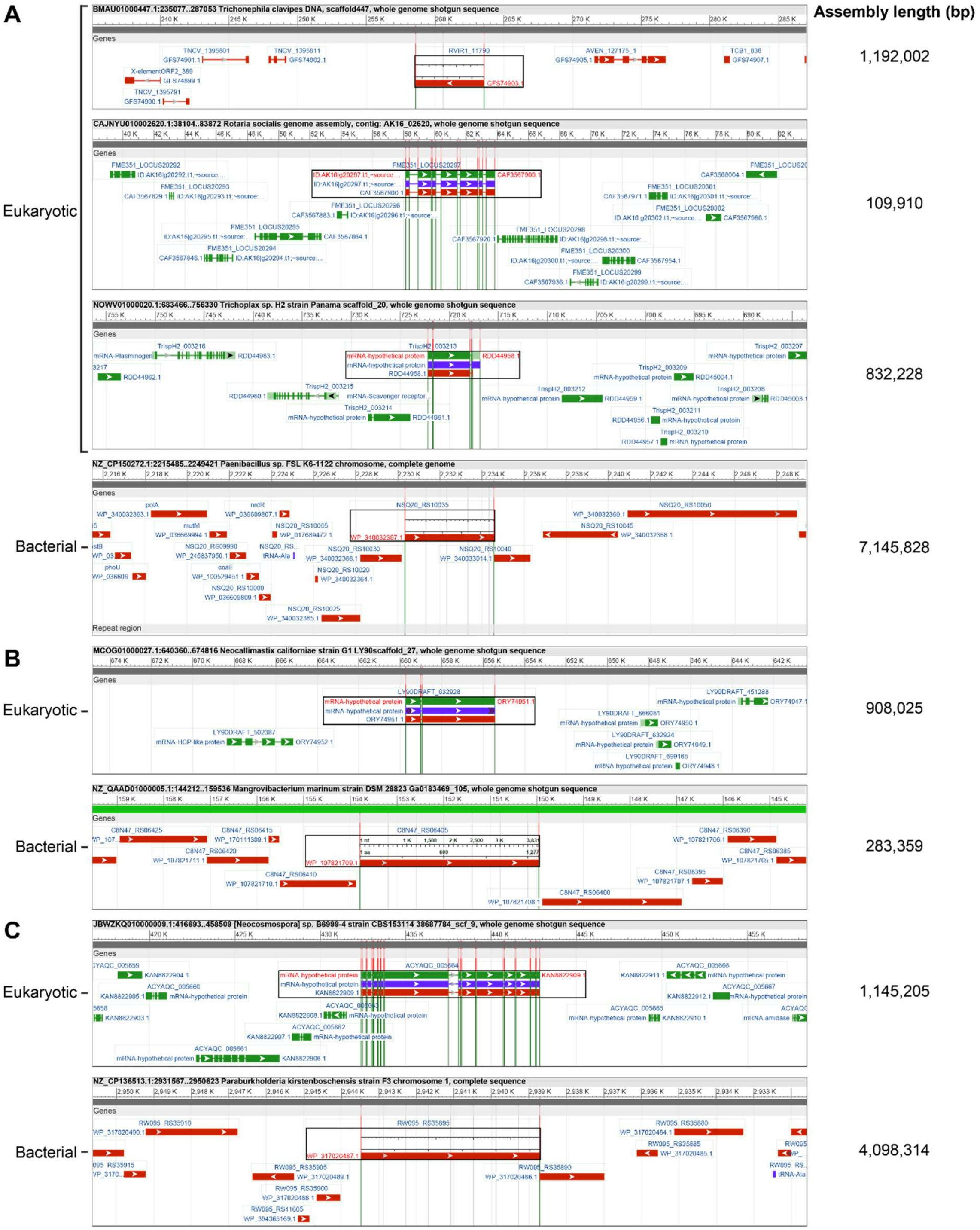


**Figure S3. Tiamat gene properties and genomic context suggests eukaryotic and prokaryotic Tiamat proteins that clade together are not sequence contaminations.** Graphical views of the Tiamat gene and surrounding genes of key representatives from clades that contained a mixture of eukaryotic and prokaryotic sequences. Data from sequences found in (**A**) Clade 4, (**B**) Clade 5, (**C**) Clade 6, are shown. Eukaryotic Tiamat genes either contain introns themselves or are next to intron-containing genes, and they neighbour eukaryotic genes. Bacterial Tiamat genes do not contain introns, and do not occur next to intron-containing genes, and they neighbour bacterial genes. Tiamat genes were found in scaffolds, contigs and complete genomes. The total length of each assembly is shown to the right of each example.

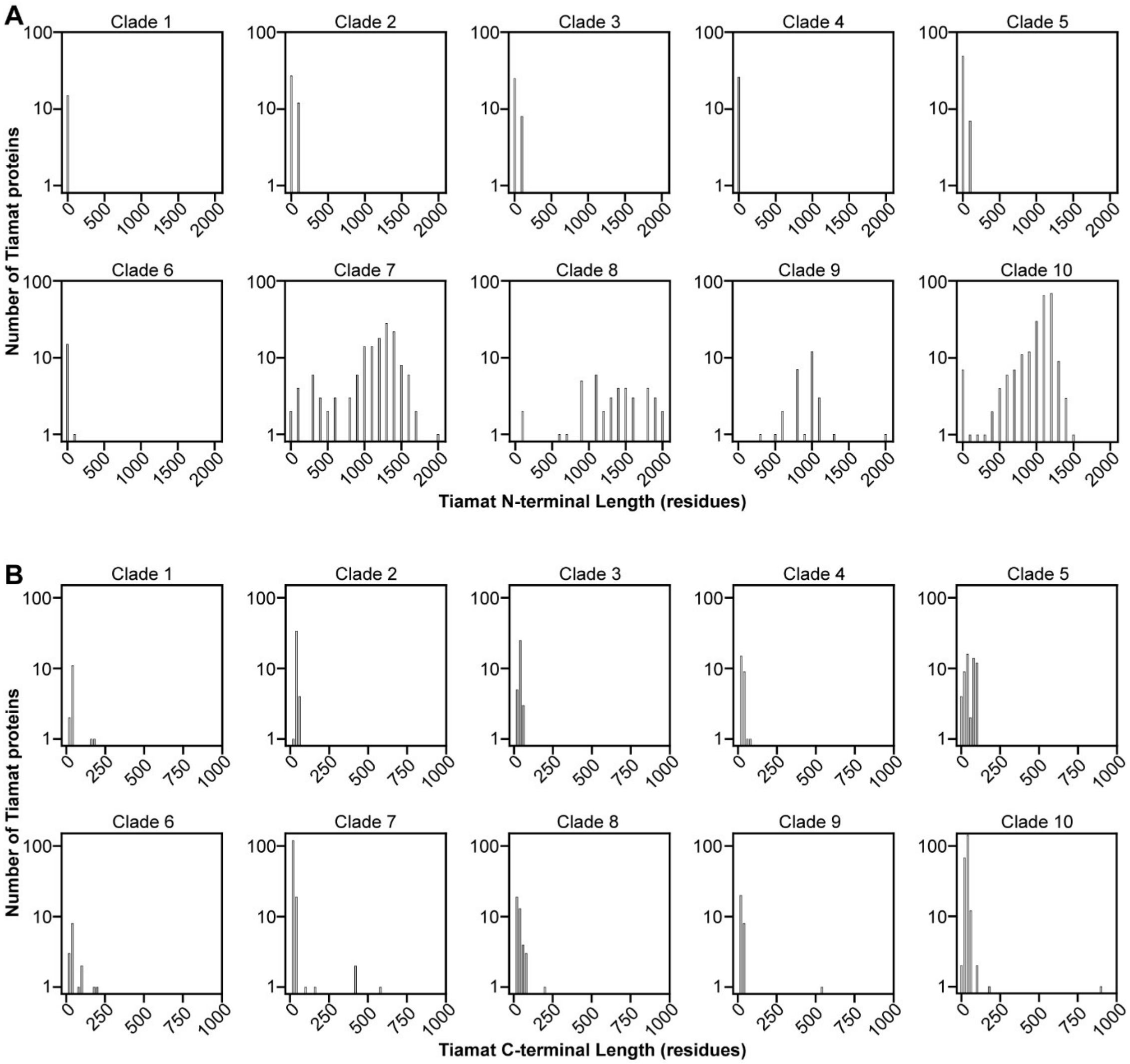


**Figure S4. Eukaryotic Tiamat proteins frequently have longer N-termini than bacterial or archaeal Tiamat proteins.** (**A**) Distribution of the lengths of extensions N-terminal to the ATPase domain of Tiamat proteins. (**B**) Distribution of the lengths of extensions C-terminal to the nuclease domain of Tiamat proteins. The sequences analyzed here originate from the UniProt reference proteome database.

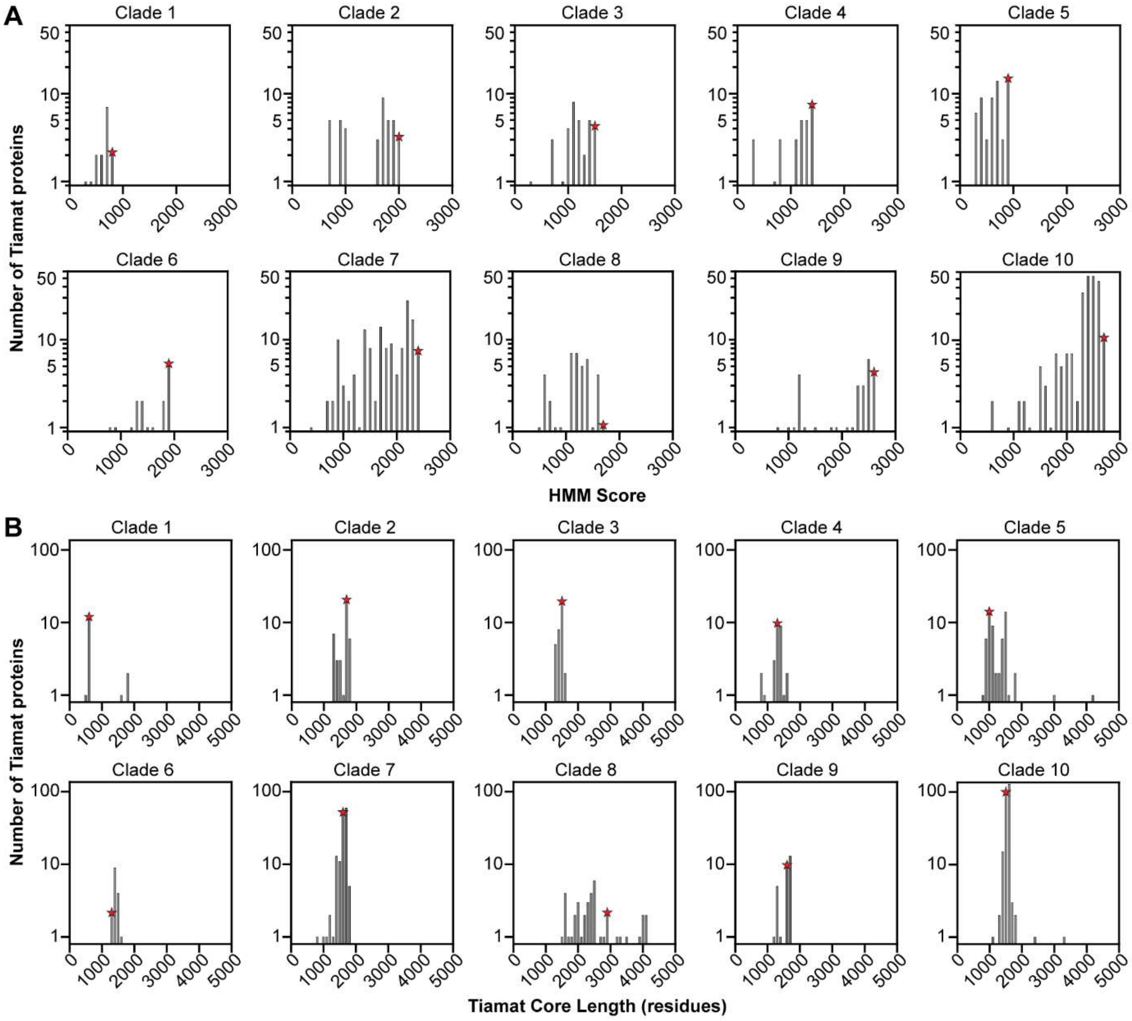


**Figure S5. HMM-guided selection of Tiamat representative proteins on per-clade basis.** (**A**) Distribution of HMM scores for Tiamat proteins within a given clade. Clade-specific HMMs of the central region of Tiamat proteins spanning the ATPase through to the nuclease domain were generated, which was then used to search and score each Tiamat protein within the same clade. A higher HMM score signifies that protein is most similar to most proteins in its clade. For each clade, the protein with the highest HMM score (red star) was chosen to be the representative for that clade. (**B**) Distribution of the lengths of the core region (ATPase domain through to PD-(D/E)xK domain) of Tiamat proteins within a given clade. The bin containing the representative protein for each clade is annotated (red star). The sequences analyzed here originate from the UniProt reference proteome database.

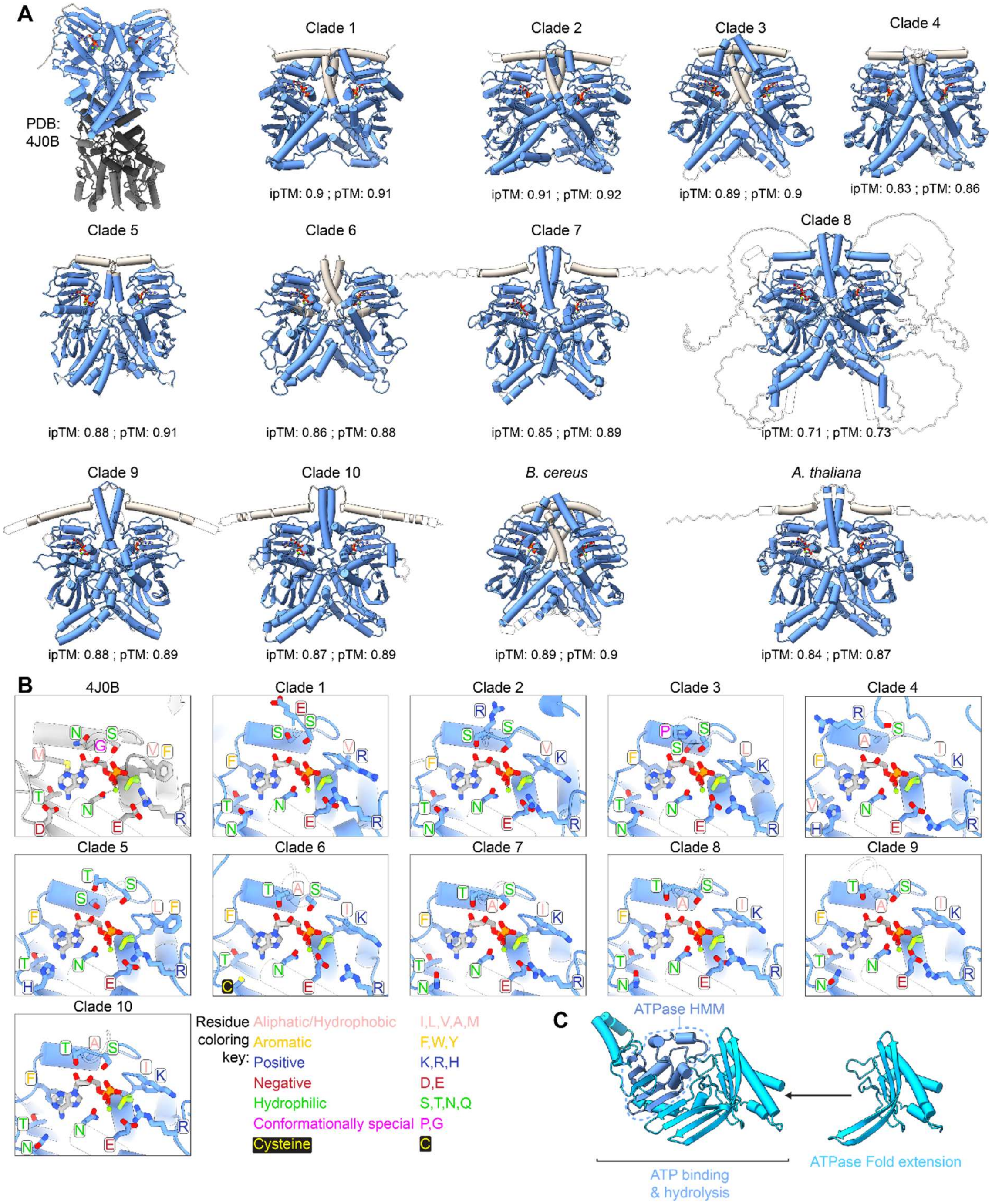


**Figure S6. Conservation of ATPase domain across Bacterial and Eukaryotic Tiamat representatives.** (**A**) Overviews of the experimentally determined structure of dimeric Hsp90 (PDB: 4J0B) and of AlphaFold3 predicted structures of dimers of the ATPase domain of each Tiamat clade representative protein, and of *B. cereus* and *A. thaliana* proteins. For Tiamat proteins with N-terminal extensions shorter than 50 residues, the entire N-terminus (20-42 residues; beige) was included in ATPase dimerization predictions. For Tiamat proteins with long N-terminal extensions, only the adjacent 40 residues were included. (**B**) Zoom-in of the active sites of all structures shown in panel A. The ADP, $BeF_3$, and the $Mg^{2+}$ ions from the experimentally determined structure are shown overlaid on all putative ATPase structures. Residue labels are colored by biochemical property, using the Zappo coloring scheme. (**C**) An example of how the ATPase annotation via HMM or InterProScan has been extended to include a region that may correspond with the "transducer" domain of well-characterized GHL proteins is shown.

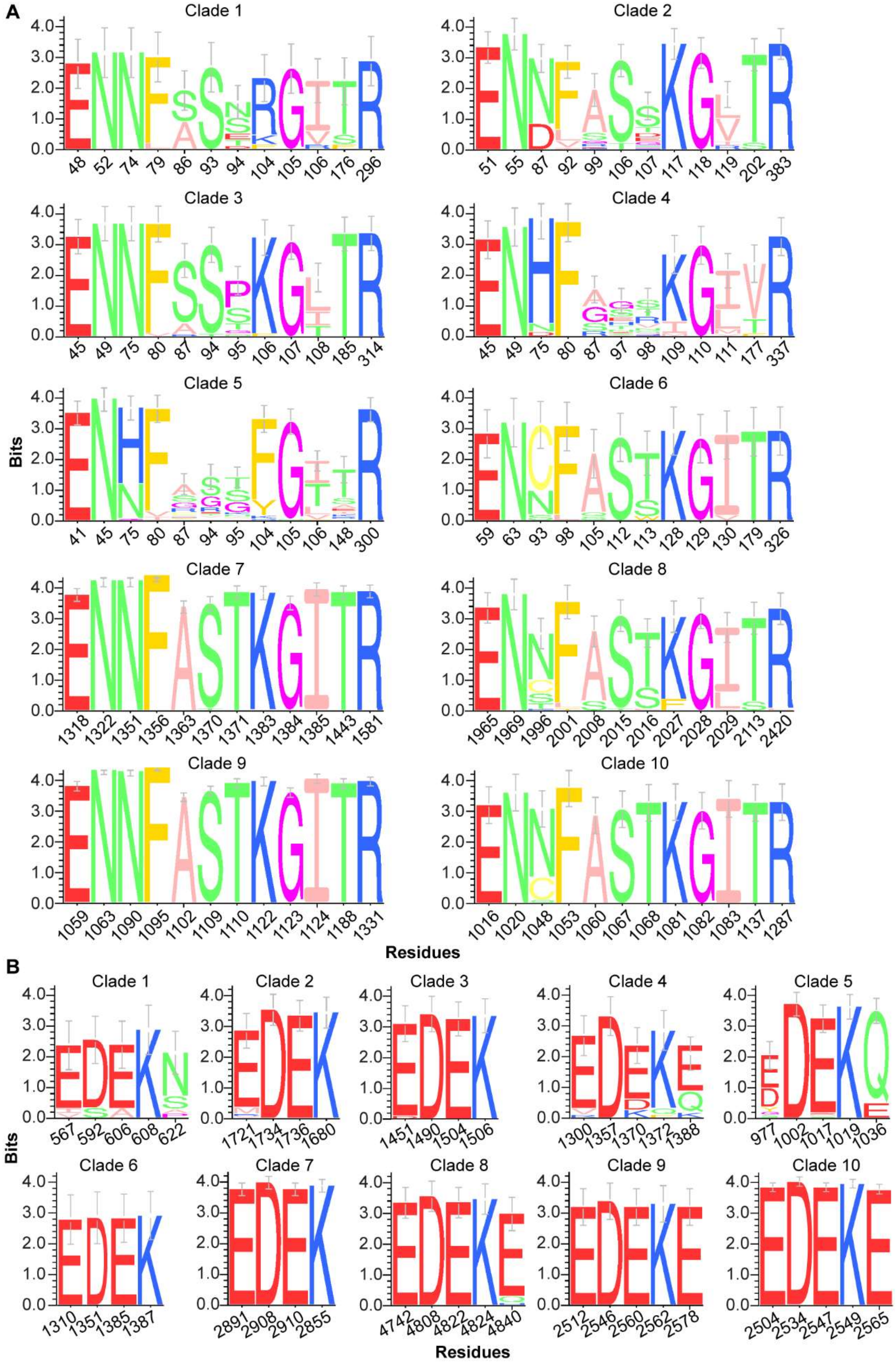


**Figure S7. ATPase and nuclease active site residues are conserved across representatives of Tiamat clades.** (**A**) Sequence logos depicting the conservation of key residues in the ATPase active site of Tiamat proteins, per clade. (**B**) Sequence logos depicting the conservation of key residues in the nuclease active site of Tiamat proteins, per clade. Residues are colored according to the "Zappo" coloring scheme, and are numbered according to the representative protein for each clade. The sequences analyzed here originate from the UniProt reference proteome database.

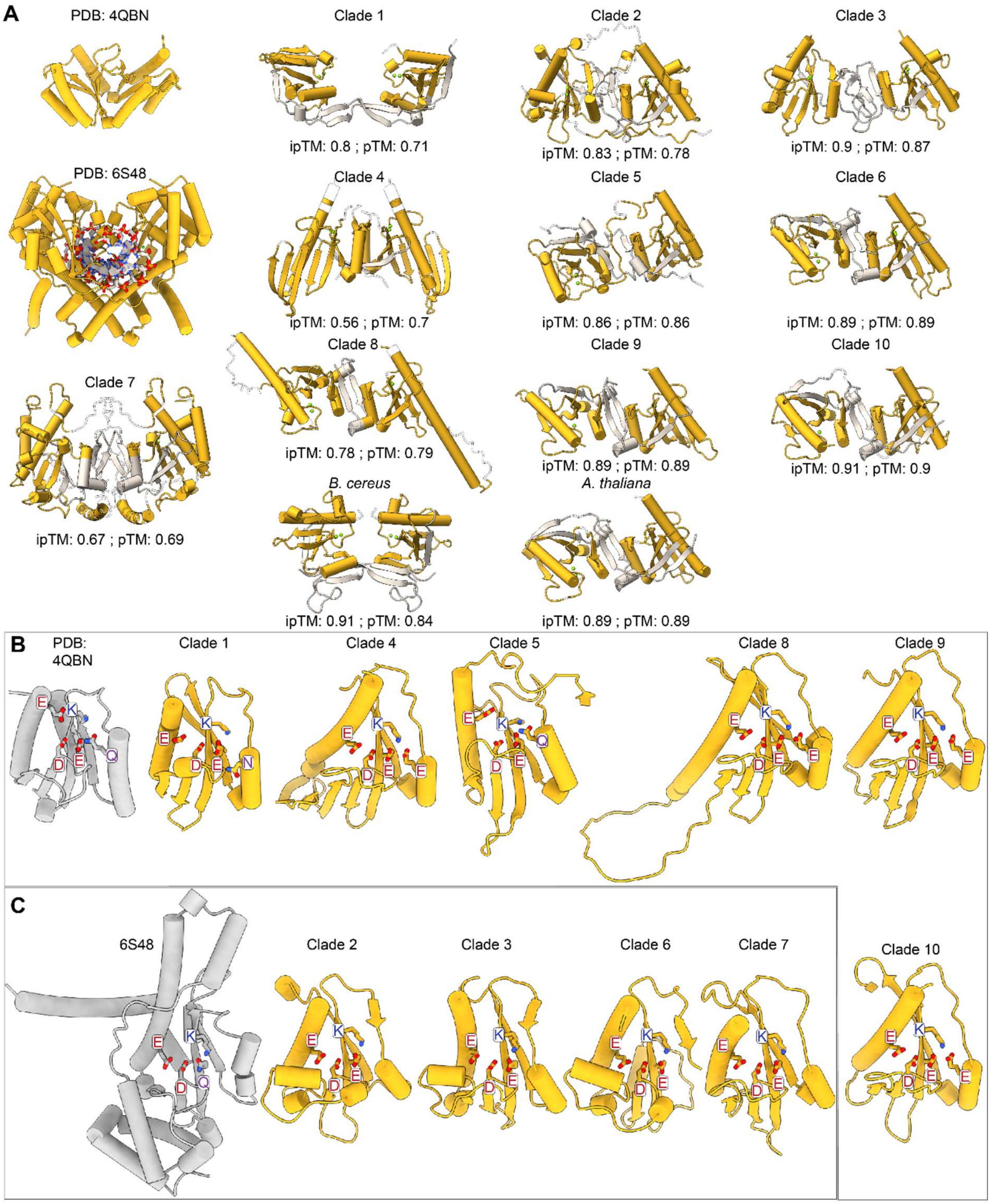


**Figure S8. Conservation of nuclease domain across Bacterial and Eukaryotic Tiamat representatives**. (**A**) Overviews of the experimentally determined structures of a dimeric bacterial Holliday junction resolvase (PDB:4QBN) and bacterial restriction endonuclease (PDB:6S48) that were most structurally similar to Tiamat nuclease domains, shown alongside predicted dimeric structures of the nuclease domains of each Tiamat clade representative protein, and of *B. cereus* and *A. thaliana* proteins. The entire C-terminus (12-66 residues) was included in nuclease dimerization predictions (beige). Three differing viewing angles are used between representative structures to best showcase the symmetry and dimerization of each example nuclease. (**B**) Experimentally determined structures for a monomer of a bacterial Holliday junction resolvase (PDB:4QBN) shown alongside similar putative PD-(D/E)XK nuclease domains from representative proteins from Tiamat homolog clades. (**C**) Experimentally determined structures for a monomer of a bacterial restriction endonuclease (PDB:6S48) shown alongside similar putative PD-(D/E)XK nuclease domains from representative proteins from Tiamat homolog clades.

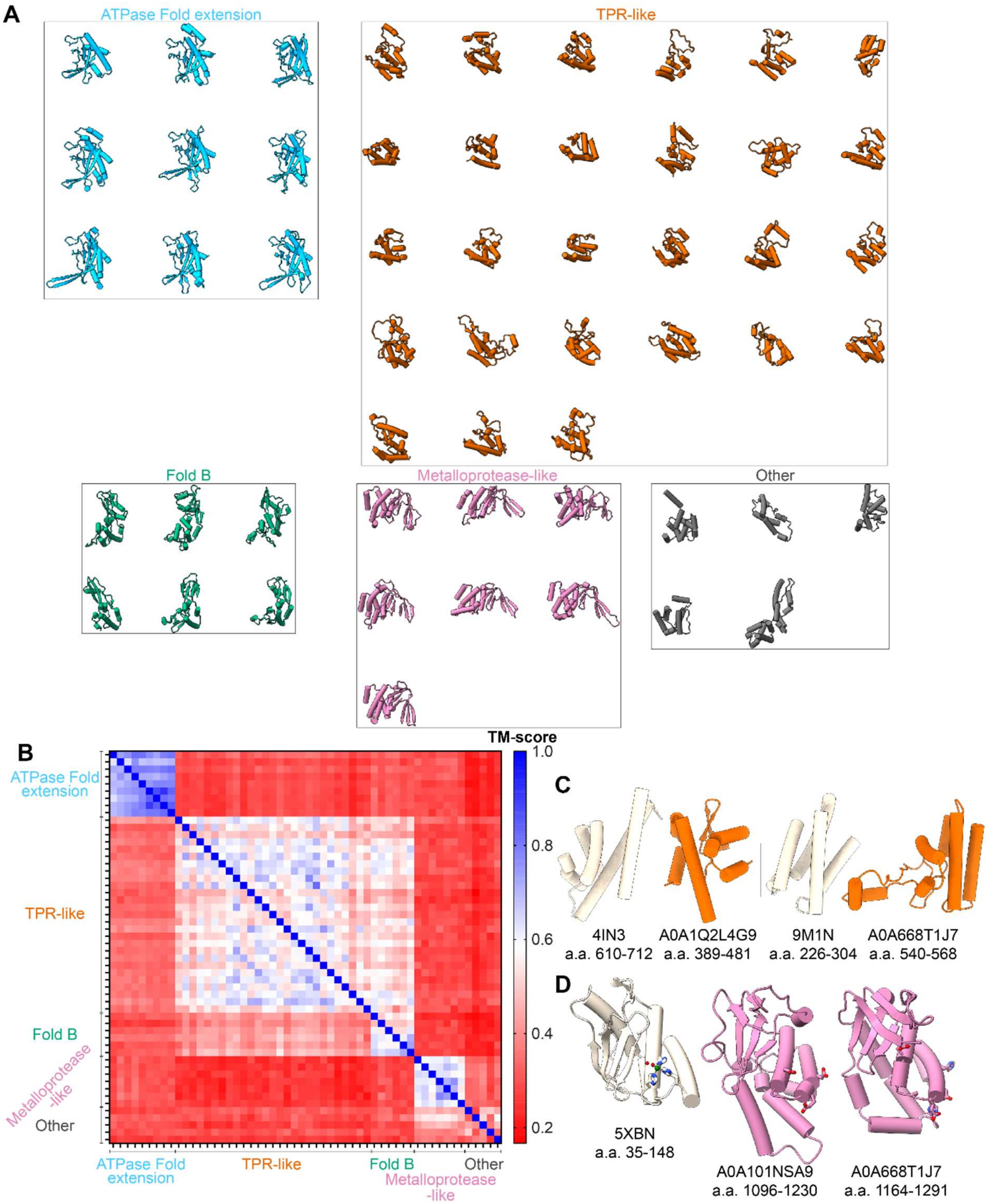


**Figure S9. The hypervariable region between the ATPase and NOV_C domains of Tiamat contain conserved protein folds.** (**A**) Gallery depicting broadly conserved folded protein domains identified in representative Tiamat proteins (see Figure 2b). (**B**) Quantitative assessment of similarity of putative protein domains via the TM-score metric (1, 2). Scores higher than 0.5 typically signify the two proteins are the same SCOP or CATH fold, whereas TM-scores below 0.17 suggest the two proteins are unrelated (2). (**C**) TPR-like folds in Tiamat proteins structurally align well to TPR domains in other proteins. Dali reports that the noted amino acid (a.a.) range from *Planococcus lenghuensis* Tiamat (A0A1Q2L4G9) matches *Saccharomyces cerevisiae* Bch1 (4IN3, chain D) with a Z-score of 5.2 and RMSD of 3.1 Å, and the noted amino acid range of *Chromis aereus* Tiamat (A0A668T1J7) matches the noted amino acids of human Tubulin-specific chaperone D (9M1N, chain D) with a Z-score of 6.5 and RMSD of 2.7 Å. The PDB ID or protein accession ID, and range of amino acids (a.a.) depicted are shown below each example protein structure. (**D**) Metalloprotease-like folds in Tiamat align well to a number of metalloprotease domains, and five of the seven representative folds from Tiamat proteins specifically align well to the Wss1 metalloprotease from *Saccharomyces cerevisiae*. Dali reports that the noted regions from *Streptomyces yokosukanensis* Tiamat (A0A101NSA9) and *C. aereus* Tiamat (A0A668T1J7) match to Wss1 (5XBN) with Z-scores of 4.2 and 6.2 and RMSDs of 4.3 Å and 2.3 Å respectively. Tiamat residues near the canonical metalloprotease active site position, and which commonly coordinate either active-site or structural zinc ions, are shown in “stick” representation (3, 4).

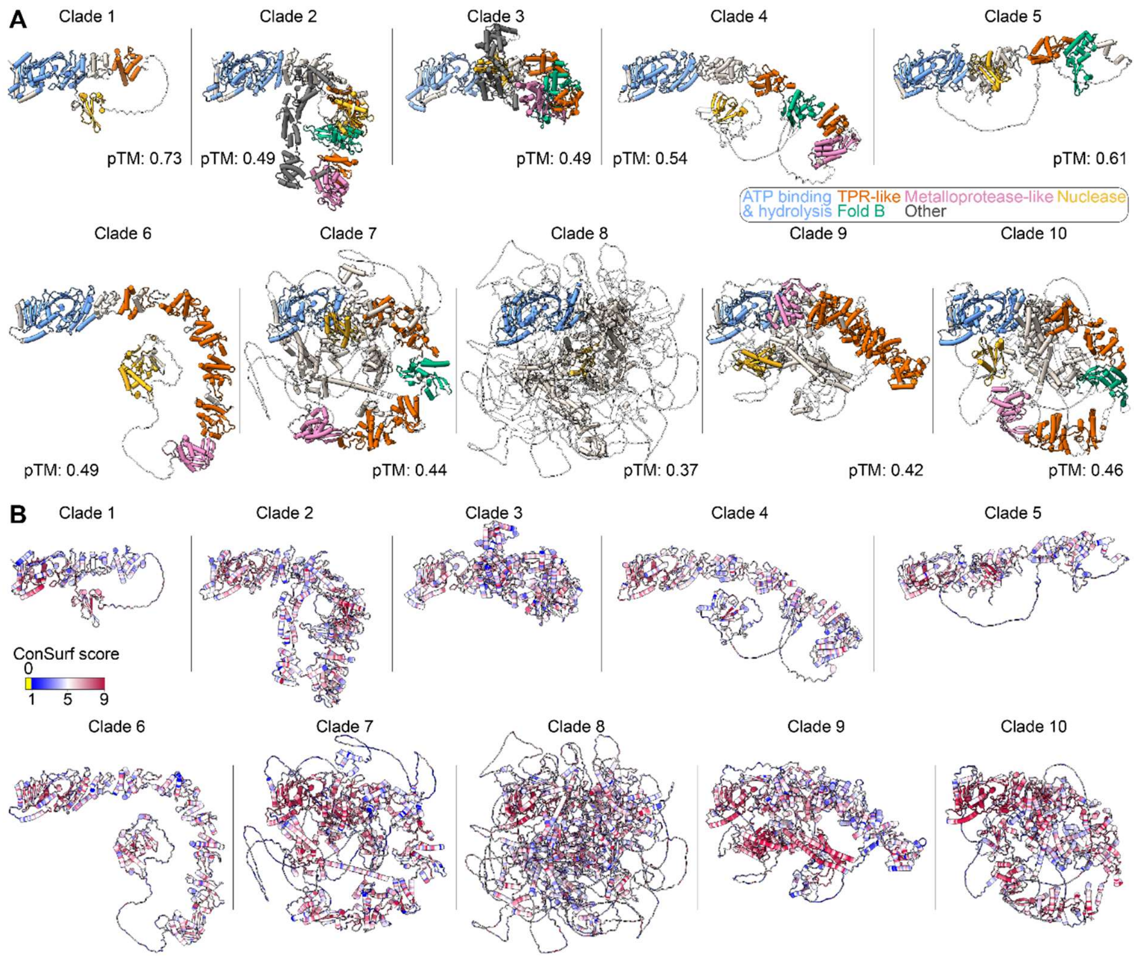


**Figure S10. Predicted structures and overall sequence conservation of Tiamat homolog representatives.** (**A**) Full-length predicted monomeric structures of the representative Tiamat proteins for all clades. 2-D schemes of these proteins are shown in main text Figure 2. The proteins are colored in beige, with putative biochemically active domains and annotated folds given additional colors. Regions of the predicted protein structures with a pLDDT score less than 50 are shown in 100% transparency. (**B**) The conservation of residues (ConSurf score) amongst Tiamat homologs within each clade is shown on the predicted structure of each clade representative (5).

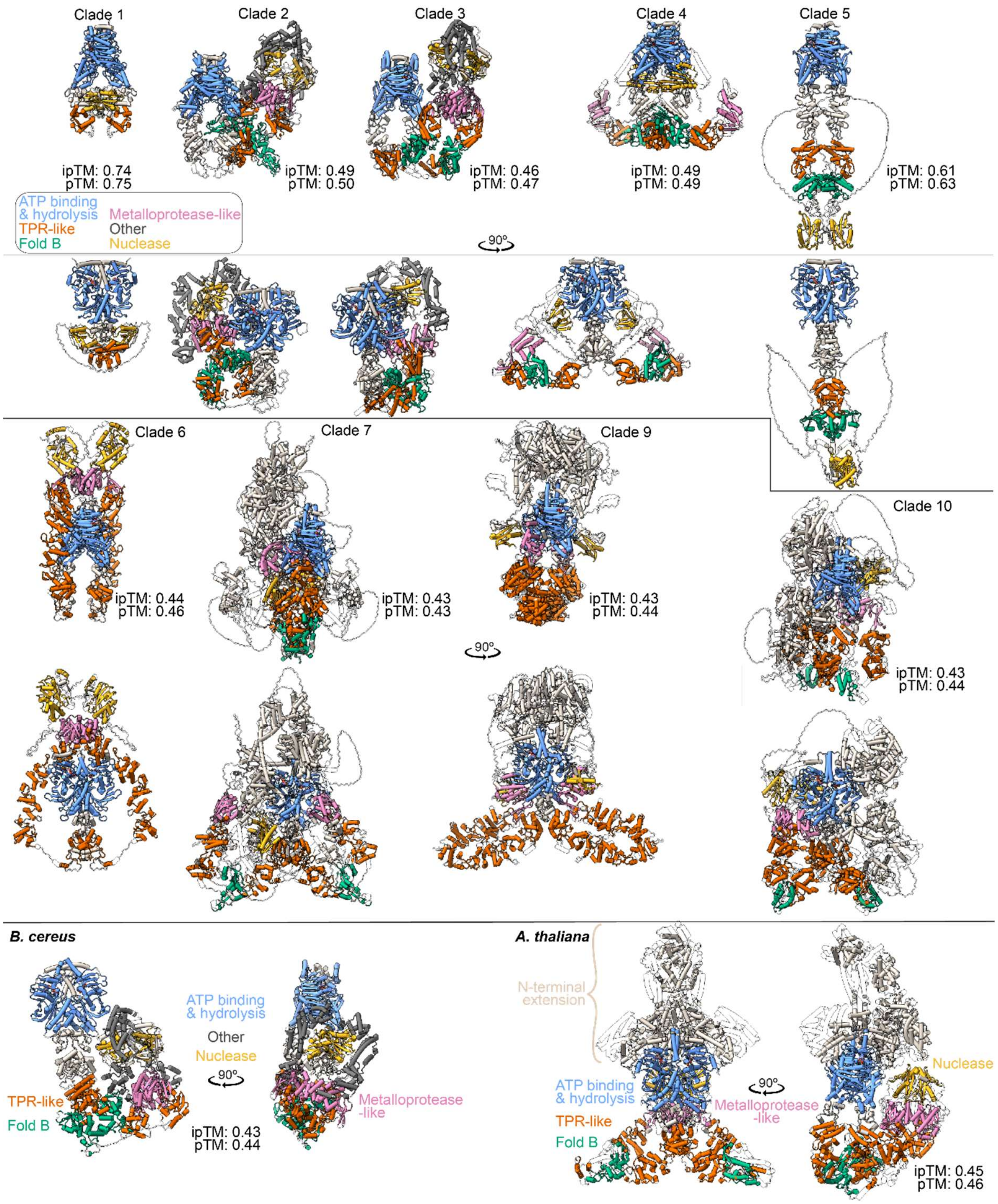


**Figure S11. Tiamat proteins are predicted to dimerize, but flexibility of the central region introduces uncertainty.** Structures of dimeric Tiamat proteins predicted by AlphaFold3 in the presence of two ATP molecules, and six $Mg^{2+}$ ions. 2-D schemes of these proteins are shown in main text Figure 2. The proteins are colored in beige, with putative biochemically active domains and annotated folds given additional colors. Regions of the predicted protein structures with a pLDDT score less than 50 are shown in 100%

transparency. Two views are shown for each protein (top, bottom) rotated by 90° relative to each other. The central region of Tiamat proteins are composed of several domains interweaved with flexible linkers. Dimers of the Clade 8 representative could not be computed.

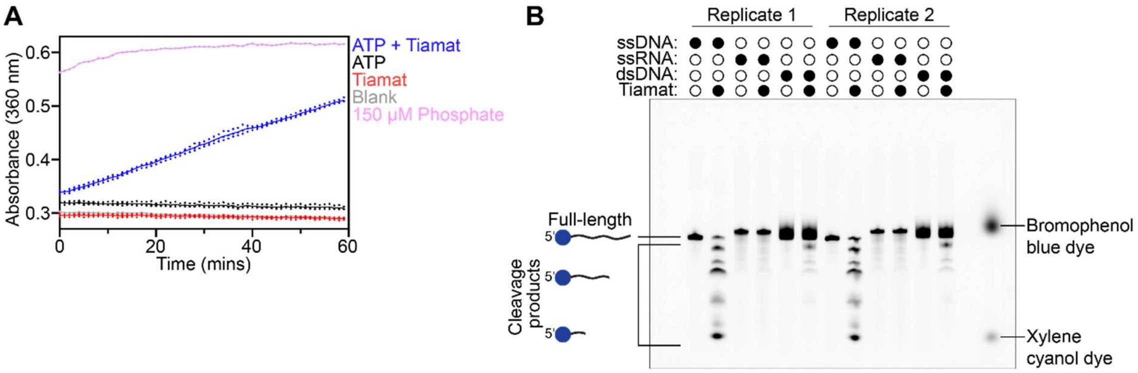


**Figure S12. *B. cereus* Tiamat is an ATPase and ssDNase.** (**A**) Raw ATPase activity assays show that ATP is hydrolyzed in the presence of Tiamat. “ATP + Tiamat”, “ATP” alone and “Tiamat” alone experiments were measured in triplicate. “Blank” was measured in duplicate. A single 150 μM phosphate-containing reaction was measured. Blank-subtracted values are plotted in Figure 3. Individual points for all assays and a line at the mean are shown. (**B**) Nuclease activity assays show that Tiamat preferentially cleaves Cy5-labelled ssDNA over dsDNA, and does not appear to cleave ssRNA. Nuclease assays were performed in duplicate. The bromophenol blue and xylene cyanol dyes are fluorescent at similar wavelengths to Cy5 and so appear in the right most lane, in which tracking dyes were added. A slightly overexposed image is shown here in comparison to Figure 3 so that the background signal in the gel can be seen.

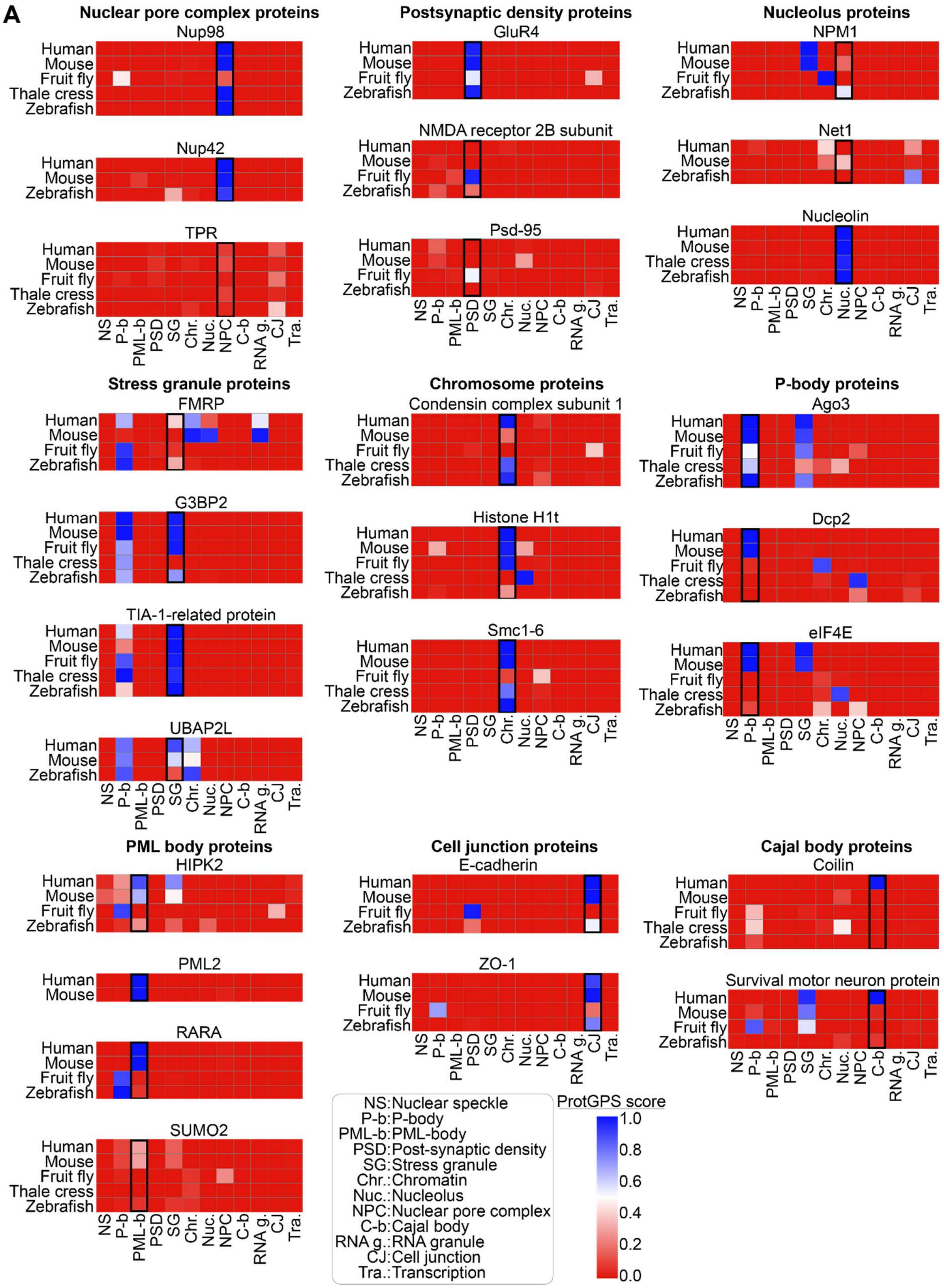


**Figure S13. ProtGPS predicts similar cellular localization for some but not all non-human protein homologs.** Heatmaps showing ProtGPS-predicted subcellular localizations for positive-control reference proteins experimentally validated to localize to specific cellular compartments across *Homo sapiens*, *Mus musculus*, *Drosophila melanogaster*, *Arabidopsis thaliana*, and *Danio rerio* (6–22). Prediction confidence scores (0.0 – 1.0) are represented by the color scale, with blue indicating the highest predicted likelihood of localization in a given compartment, and red indicating the lowest predicted likelihood. As a caveat, ProtGPS was trained on human proteins (23), and while many proteins that are characteristic signatures of subcellular locations such as the P-body or nuclear pore complex, are strongly conserved across eukaryotes it is possible that there is a higher false positive rate for annotation of non-human proteins.

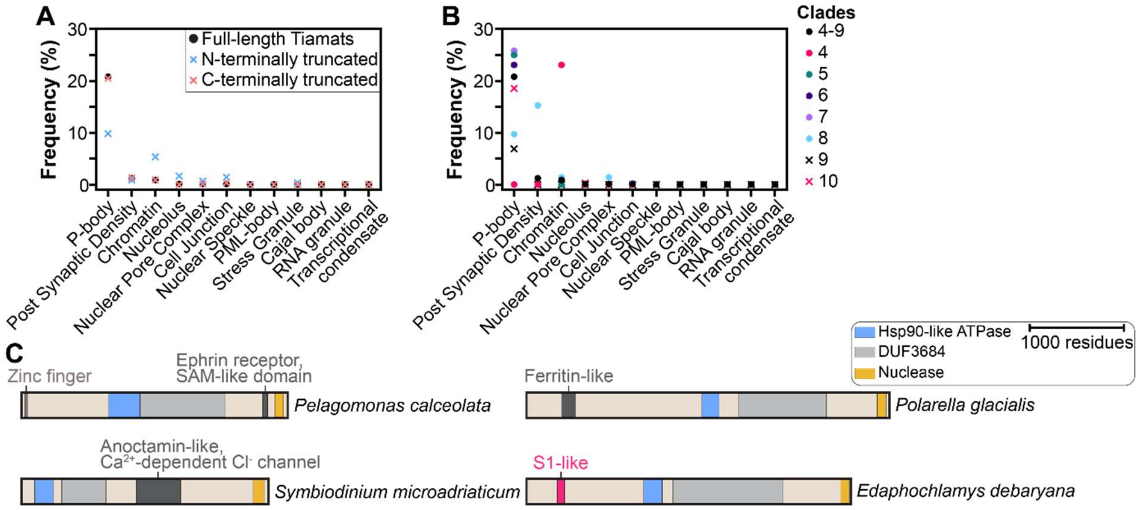


**Figure S14. The N-terminus of eukaryotic Tiamat proteins drives many ProtGPS-based localization predictions**. (**A**) ProtGPS prediction of the subcellular localization of full-length (black circles), N-terminally truncated (blue crosses), or C-terminally truncated (red crosses) eukaryotic Tiamat proteins. The N-terminally truncated Tiamat proteins have a decreased predicted propensity to localize to P-bodies and an increased propensity to localize to chromatin, relative to full-length Tiamat. Whereas C-terminally truncated Tiamat proteins have a similar predicted subcellular localization to full-length Tiamat proteins. (**B**) Analysis of subcellular localization of Tiamat proteins on a per-clade basis. 6-26% of Tiamat proteins within clades 5-10 are predicted to localize to the P-body. Clade 8 is enriched in proteins that are predicted to localize to the post-synaptic density (11 of 72 proteins). Clade 4 appears enriched in proteins that are predicted to localize to chromatin, though this clade has a small number of eukaryotic proteins (13 total), such that 3 proteins represent 23% of proteins in the clade. (**C**) Domain annotations of four of the eleven Tiamat proteins predicted to localize to the post-synaptic density. The sequences analyzed here originate from the UniProt reference proteome and RefSeq databases.

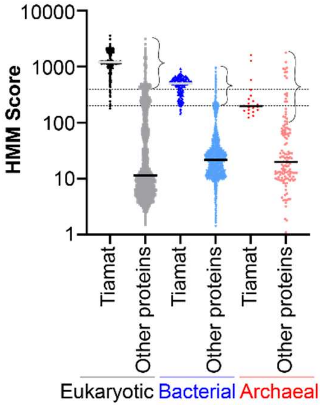


**Figure S15. Distribution of HMM scores guides downstream analysis of Tiamat-like homologs.** A search for Tiamat homologs in the UniProt and RefSeq databases was performed using custom HMMs built from eukaryotic, bacterial and archaeal sequences that yielded an initial list of proteins which was initially strictly curated to yield highly confident Tiamat homologs ("Tiamat" in graph) (see Figure S1 and Methods). Most proteins that did not pass initial curation have low HMM scores, such that median (horizontal grey or black bar across datapoints) values range from roughly 10-30. However, a fraction of proteins in this group of "Other proteins" that had been excluded from initial analyses possess high HMM scores. HMM score cutoffs of 400, 200 and 100 were used to select distantly related Tiamat-like proteins from eukaryotes, bacteria and archaea for downstream analyses respectively. Dashed lines are indicated at Y=400 and Y=200, and Y=100 is indicated on the y-axis. Curly brackets depict the proteins considered to be Tiamat-like for each domain of life.

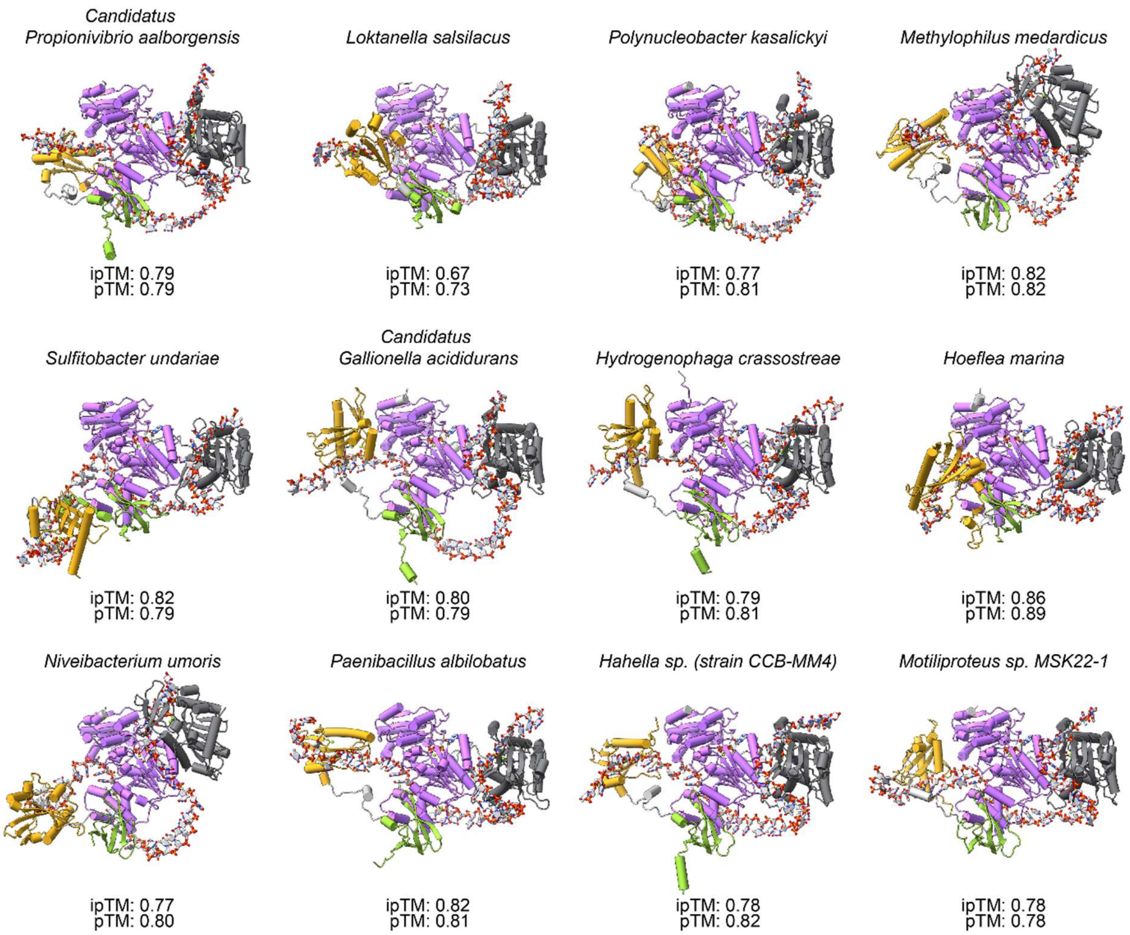


**Figure S16. Prediction of interactions between the C-terminus of Type I-G Tiamat proteins and adjacently encoded NERD-helicase.** Predicted structures of the C-terminus of Type I-G Tiamat proteins with the associated NERD-helicase protein, in the presence of one ATP, a 40-nt long T-rich DNA strand, and five $Mg^{2+}$ ions. In all cases the unique β-barrel fused to the C-terminus of Type I-G Tiamat proteins appears to recruit the NERD-helicase protein by binding the same surface of the helicase.

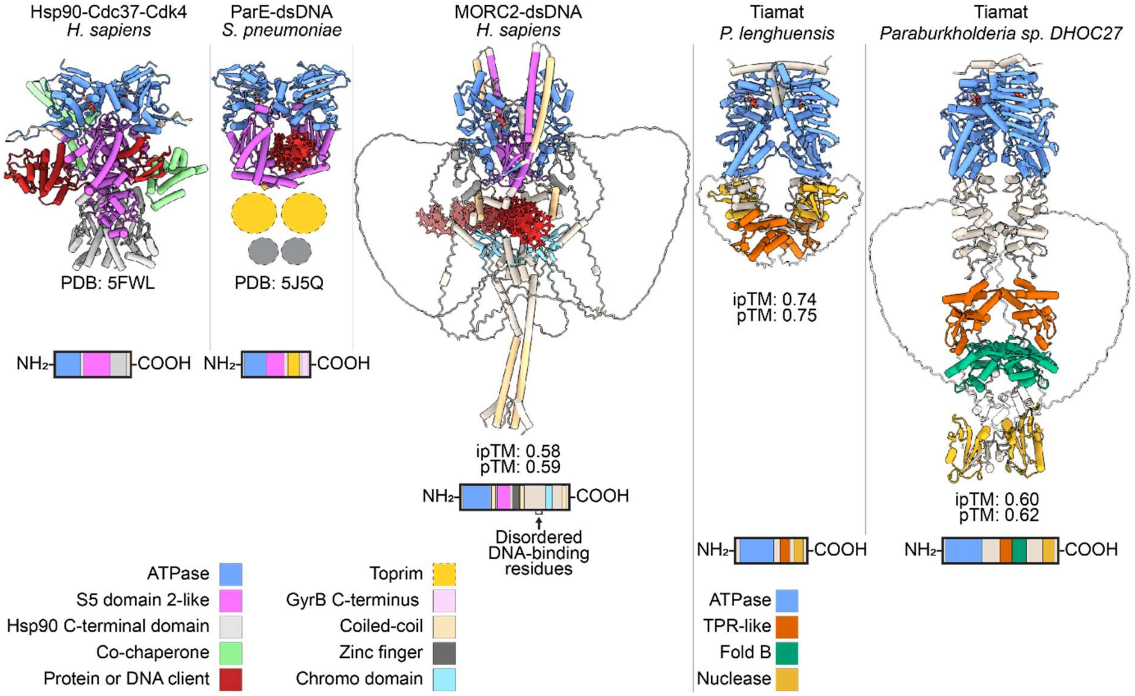


**Figure S17. Comparison of Tiamat to other GHKL ATPases suggests a mode of viral recognition.** GHKL ATPases such as the protein chaperone Hsp90, and the DNA type II topoisomerase ParE, bind client proteins or DNA in between two monomers. Clients are thought to bind to an open, ADP-bound state of the ATPase. Subsequent binding of ATP induces dimerization of the ATPase domains and closing of the dimer around the client protein or DNA. (**A**) Hsp90 works with its co-chaperone Cdc37 to bind to unfolded Cdk4 (24). Upon ATP hydrolysis, Hsp90 re-opens. Since the C-terminus of Hsp90 dimerizes too, the two Hsp90 monomers stay connected. If Cdk4 folds into its native state it is proposed to displace Cdc37 and dissociate from the chaperone. If Cdk4 does not refold, it will undergo successive ATP-powered cycles of Hsp90 closing and opening until it does. Cdk4 is bound by the so-called “middle domain” of Hsp90, annotated by InterProScan as S5 domain 2-like. (**B**) Topoisomerases change DNA topology by generating a transient DNA break, transferring a second duplex through this break, and rejoining the DNA. ParE is a type II topoisomerase which consists of an ATPase domain, a domain annotated as ribosomal protein S5 domain 2-like by InterProScan, and a C-terminal Toprim domain. Toprim is a nuclease domain found in housekeeping proteins like topoisomerases and primases, and in antiphage defense systems such as Gabija and PARIS (25–27). A structure of ParE is pictured trapping the DNA to be transferred (T-segment) across a transient double-stranded DNA break. The Toprim domain was not resolved in the original structure, and its relative position is indicated. (**C**) MORC proteins are involved in eukaryotic heterochromatin condensation and gene silencing (28). Transient opening of the MORC2 homodimer by separation of the two ATPase domains is proposed to allow for DNA substrates to enter and bind the central portion of MORC2. Since the C-terminal coiled-coil domains of MORC2 dimerize, the two MORC2 monomers remain connected (29). DNA binding to the central region of MORC2 may stimulate ATP binding and the dimerization of the ATPase domains, to clamp the DNA. DNA is predicted by AlphaFold3 to thread through the center of MORC2, however major DNA binding residues are disordered in this prediction and looped out into two "wings" (29, 30). (**D**) Predicted structures of Tiamat homologs suggests that the ATPase and nuclease domains may both dimerize. Therefore, similar to other GHKL ATPases, ADP-bound Tiamat may transiently open and form a separation between its monomers at its N-terminus to allow for a protein or DNA substrate to enter and bind its central lumen. Like other GHKL ATPases, subsequent ATP binding may trigger dimerization of the ATPase domains and clamping down of the two Tiamat monomers onto the substrate.

**REFERENCES**


1. Zhang,Y. and Skolnick,J. (2004) Scoring function for automated assessment of protein structure template quality. *Proteins Struct. Funct. Bioinforma.*, **57**, 702–710.

2. Xu,J. and Zhang,Y. (2010) How significant is a protein structure similarity with TM-score = 0.5? *Bioinformatics*, **26**, 889–895.

3. Vallee,B.L. and Auld,D.S. (1990) Zinc coordination, function, and structure of zinc enzymes and other proteins. *Biochemistry*, **29**, 5647–5659.

4. Jongeneel,C.V., Bouvier,J. and Bairoch,A. (1989) A unique signature identifies a family of zinc-dependent metallopeptidases. *FEBS Lett.*, **242**, 211–214.

5. Ashkenazy,H., Abadi,S., Martz,E., Chay,O., Mayrose,I., Pupko,T. and Ben-Tal,N. (2016) ConSurf 2016: an improved methodology to estimate and visualize evolutionary conservation in macromolecules. *Nucleic Acids Res.*, **44**, W344–W350.

6. Ferraiuolo,M.A., Basak,S., Dostie,J., Murray,E.L., Schoenberg,D.R. and Sonenberg,N. (2005) A role for the eIF4E-binding protein 4E-T in P-body formation and mRNA decay. *J. Cell Biol.*, **170**, 913–924.

7. Sen,G.L. and Blau,H.M. (2005) Argonaute 2/RISC resides in sites of mammalian mRNA decay known as cytoplasmic bodies. *Nat. Cell Biol.*, **7**, 633–636.

8. Liu,J., Valencia-Sanchez,M.A., Hannon,G.J. and Parker,R. (2005) MicroRNA-dependent localization of targeted mRNAs to mammalian P-bodies. *Nat. Cell Biol.*, **7**, 719–723.

9. Tishinov,K. and Spang,A. (2021) The mRNA decapping complex is buffered by nuclear localization. *J. Cell Sci.*, **134**, jcs259156.

10. Guan,D. and Kao,H.-Y. (2015) The function, regulation and therapeutic implications of the tumor suppressor protein, PML. *Cell Biosci.*, **5**, 60.

11. Gulyássy,P., Puska,G., Györffy,B.A., Todorov-Völgyi,K., Juhász,G., Drahos,L. and Kékesi,K.A. (2020) Proteomic comparison of different synaptosome preparation procedures. *Amino Acids*, **52**, 1529–1543.

12. Barki-Harrington,L., Elkobi,A., Tzabary,T. and Rosenblum,K. (2009) Tyrosine Phosphorylation of the 2B Subunit of the NMDA Receptor Is Necessary for Taste Memory Formation. *J. Neurosci.*, **29**, 9219–9226.

13. Matsubara,A., Laake,J.H., Davanger,S., Usami,S. and Ottersen,O.P. (1996) Organization of AMPA Receptor Subunits at a Glutamate Synapse: A Quantitative Immunogold Analysis of Hair Cell Synapses in the Rat Organ of Corti. *J. Neurosci.*, **16**, 4457–4467.

14. Meyer,C., Garzia,A., Mazzola,M., Gerstberger,S., Molina,H. and Tuschl,T. (2018) The TIA1 RNA-Binding Protein Family Regulates EIF2AK2-Mediated Stress Response and Cell Cycle Progression. *Mol. Cell*, **69**, 622-635.e6.

15. Maruri-López,I., Figueroa,N.E., Hernández-Sánchez,I.E. and Chodasiewicz,M. (2021) Plant Stress Granules: Trends and Beyond. *Front. Plant Sci.*, **12**, 722643.

16. Hirano,T. (2006) At the heart of the chromosome: SMC proteins in action. *Nat. Rev. Mol. Cell Biol.*, **7**, 311–322.

17. Engbrecht,M., Grundei,D., Dilger,A.M., Wiedemann,H., Aust,A.-K., Baumgärtner,S., Helfrich,S., Kergl-Räpple,F., Bürkle,A. and Mangerich,A. (2024) Monitoring nucleolar-nucleoplasmic protein shuttling in living cells by high-content microscopy and automated image analysis. *Nucleic Acids Res.*, **52**, e72–e72.

18. Geil,C., Schwab,M. and Seufert,W. (2008) A Nucleolus-Localized Activator of Cdc14 Phosphatase Supports rDNA Segregation in Yeast Mitosis. *Curr. Biol.*, **18**, 1001–1005.

19. Strambio-De-Castillia,C., Niepel,M. and Rout,M.P. (2010) The nuclear pore complex: bridging nuclear transport and gene regulation. *Nat. Rev. Mol. Cell Biol.*, **11**, 490–501.

20. Neugebauer,K.M. (2017) Special focus on the Cajal Body. *RNA Biol.*, **14**, 669–670.

21. Nizami,Z., Deryusheva,S. and Gall,J.G. (2010) The Cajal Body and Histone Locus Body. *Cold Spring Harb. Perspect. Biol.*, **2**, a000653–a000653.

22. Sang,Q., Wang,G., Morton,D.B., Wu,H. and Xie,B. (2021) The ZO-1 protein Polychaetoid as an upstream regulator of the Hippo pathway in Drosophila. *PLOS Genet.*, **17**, e1009894.

23. Kilgore,H.R., Chinn,I., Mikhael,P.G., Mitnikov,I., Dongen,C.V., Zylberberg,G., Afeyan,L., Banani,S.F., Wilson-Hawken,S., Lee,T.I., *et al.* (2025) Protein codes promote selective subcellular compartmentalization. 10.1126/science.adq2634.

24. Verba,K.A., Wang,R.Y.-R., Arakawa,A., Liu,Y., Shirouzu,M., Yokoyama,S. and Agard,D.A. (2016) Atomic structure of Hsp90-Cdc37-Cdk4 reveals that Hsp90 traps and stabilizes an unfolded kinase. *Science*, **352**, 1542–1547.

25. Antine,S.P., Johnson,A.G., Mooney,S.E., Leavitt,A., Mayer,M.L., Yirmiya,E., Amitai,G., Sorek,R. and Kranzusch,P.J. (2024) Structural basis of Gabija anti-phage defence and viral immune evasion. *Nature*, **625**, 360–365.

26. Aravind,L. (1998) Toprim--a conserved catalytic domain in type IA and II topoisomerases, DnaG-type primases, OLD family nucleases and RecR proteins. *Nucleic Acids Res.*, **26**, 4205–4213.

27. Burman,N., Belukhina,S., Depardieu,F., Wilkinson,R.A., Skutel,M., Santiago-Frangos,A., Graham,A.B., Livenskyi,A., Chechenina,A., Morozova,N., *et al.* (2024) A virally encoded tRNA neutralizes the PARIS antiviral defence system. *Nature*, **634**, 424–431.

28. Moissiard,G., Cokus,S.J., Cary,J., Feng,S., Billi,A.C., Stroud,H., Husmann,D., Zhan,Y., Lajoie,B.R., McCord,R.P., *et al.* (2012) MORC Family ATPases Required for Heterochromatin Condensation and Gene Silencing. *Science*, **336**, 1448–1451.

29. Fendler,N.L., Ly,J., Welp,L., Lu,D., Schulte,F., Urlaub,H. and Vos,S.M. (2025) Identification and characterization of a human MORC2 DNA binding region that is required for gene silencing. *Nucleic Acids Res.*, **53**, gkae1273.

30. Abramson,J., Adler,J., Dunger,J., Evans,R., Green,T., Pritzel,A., Ronneberger,O., Willmore,L., Ballard,A.J., Bambrick,J., *et al.* (2024) Accurate structure prediction of biomolecular interactions with AlphaFold 3. *Nature*, 10.1038/s41586-024-07487-w.
